# Concurrent category-selective neural activity across the ventral occipito-temporal cortex supports a non-hierarchical view of human visual recognition

**DOI:** 10.1101/2025.07.28.666781

**Authors:** Corentin Jacques, Jacques Jonas, Sophie Colnat-Coulbois, Bruno Rossion

## Abstract

Visual recognition is a fundamental human brain function, supported by a network of regions in the ventral occipito-temporal cortex (VOTC). This network is thought to be organized hierarchically, with definite processing stages increasing in invariance and time-course from posterior to anterior cortical regions. Here we provide a stringent test of this view by measuring category-selective neural activity to natural images of faces across the VOTC with electrophysiological intracerebral recordings in a large human sample (N=140; >11000 recording sites). Face-selective high frequency broadband (30-160 Hz) neural activity is distributed across the VOTC, with right-hemispheric dominance and regional peaks of activity. Crucially, while a progressive increase in degree of category-selectivity is found along the postero-anterior axis, neural activity occurs largely concurrently (∼100ms onset – ∼450ms offset) across all VOTC regions. These observations challenge the standard hierarchical view of neural organization of visual object recognition in the human association cortex, supporting alternative models of this key brain function.

## INTRODUCTION

Recognition, i.e., reproducible discrimination, of signals from the sensory environment is a fundamental function of all central nervous systems (Edelman, 2004). Vision is considered the dominant sensory modality for most primates, who have high visual acuity and excellent binocular vision for recognizing their environment(Jacobs, 1999). In the primate order, visual object recognition is supported by a wide bilateral network of brain regions, including subcortical structures and the occipital and temporal cortices(Conway, 2018; DiCarlo et al., 2012; Ungerleider and Bell, 2011). Based primarily on non-human primates research, this network is generally conceived as being organized in a hierarchical manner, with definite processing stages increasing progressively in complexity of representation from posterior to anterior cortical regions, leading ultimately to rich invariant visual representations readily available for memory associations and behavior planning (Conway, 2018; DiCarlo et al., 2012; Freiwald, 2020; Freiwald and Tsao, 2010; Grill-Spector et al., 2017; Hubel and Wiesel, 1968; Issa et al., 2018; Marr, 1982; Riesenhuber and Poggio, 1999; Tsao, 2014; Van Essen et al., 1992; Zeki and Shipp, 1988). In this hierarchical network, the selective response properties of neural populations in a given brain region are thought to stem from the combination of simpler responses from lower levels.

However, beyond the initial cascades of activities in early sensory brain structures (i.e., retina, lateral geniculate nucleus, primary visual cortices), whether visual object recognition is organized hierarchically remains largely unknown and disputed (Bullier, 2001; Eldridge et al., 2023; Hegdé and Felleman, 2007; Kravitz et al., 2013; Mumford, 1992; Rossion, 2022). This is particularly the case within the temporal association cortex, which is disproportionately enlarged in humans compared to other primates (Braunsdorf et al., 2021; Buckner and Krienen, 2013) and holds most category-selective ventral regions that are key for visual object recognition in our species (Bracci et al., 2017; Grill-Spector and Weiner, 2014; Rossion et al., 2018).

Here we provide important information to evaluate the hierarchical view of human visual object recognition through an extensive characterization of the time-course of category-selective brain activity along the human ventral occipito-temporal cortex (VOTC). To do so, we take advantage of the high spatial and temporal resolution provided by intracerebral electroencephalographic (iEEG) recordings in a particularly large sample of individual human brains (N=140) implanted from posterior to anterior regions of their VOTC (>11000 intracerebral recording sites). With iEEG frequency-tagging, we isolate category-selective high frequency (‘high frequency broadband’, 30-160 Hz) neural activity to natural images of faces – arguably the most familiar and ecologically valid stimulus in the human environment-, to characterize its time-course across the whole VOTC. This allows testing two essential features of a hierarchical organization (Bullier and Nowak, 1995; Schmolesky et al., 1998): (1) an increase in representation complexity, or abstraction, along the VOTC and (2) a progressive increase in the onset time of the earliest neural response at each level of the hierarchy, its inputs being driven by the output from the previous level.

Relative to natural images of various object shapes, we find consistent face-selective neural activity throughout the VOTC, with a progressive increase in face-selectivity, or abstraction, along the postero-anterior axis, reaching a high proportion of exclusive response to faces in the most anterior regions of the ventral temporal lobe. While these findings apparently support a hierarchical organization of visual recognition, face-selective neural activity occurs largely concurrently (∼100ms onset – ∼450ms offset latency) across the VOTC, challenging a hierarchical organization of visual object recognition in the human association cortex.

## RESULTS

Neural signal was measured in 140 human participants implanted with intracerebral electrodes (**Figure 1A**) while they viewed sequences of variable natural images of objects presented at a rapid periodic rate (6 Hz, or 167ms SOA, one fixation/image). Highly variable natural images of faces appeared every 5 stimuli (i.e., 6/5 = 1.2 Hz; **Figure 1B-C**). This stimulation mode has been validated in tens of studies with multiple recording techniques (e.g., Electroencephalography (EEG) (Rossion et al., 2015); Magnetoencephalography (MEG) (Hauk et al., 2021); intracerebral recordings(Jacques et al., 2022; Jonas et al., 2016a); functional magnetic resonance imaging (fMRI)(Gao et al., 2018)), providing high signal-to-noise ratio (SNR) face-selective activity. In this stimulation mode, the periodic modulation of the high frequency broadband (HFB, 30-160 Hz, **Figure 1D**) signal allows to objectively (i.e., at pre-determined frequencies (Norcia et al., 2015; Regan, 1989)) identify and quantify both the neural activity common to faces and nonface objects at 6 Hz and harmonics, as well as the face-selective (i.e., differential) activity identified at the frequency of face stimulation (1.2 Hz) and specific harmonics (e.g., 2.4 Hz, 3.6 Hz,…; **Figure 1E**) in the frequency-domain(Jacques et al., 2022; Retter and Rossion, 2016) (z > 3.1; *p* < 0.001). The use of HFB signal, considered as a local signal (relative to higher SNR phase-locked low frequency signals, (Jacques et al., 2022)) correlated with local population spiking activity (Manning et al., 2009) as well as BOLD signal (Jacques et al., 2016b), was intended to ensure maximal signal independence between VOTC regions investigated. Potential dependencies due to electrical field propagation were further limited by the use of local bipolar referencing.

**Figure 1.**
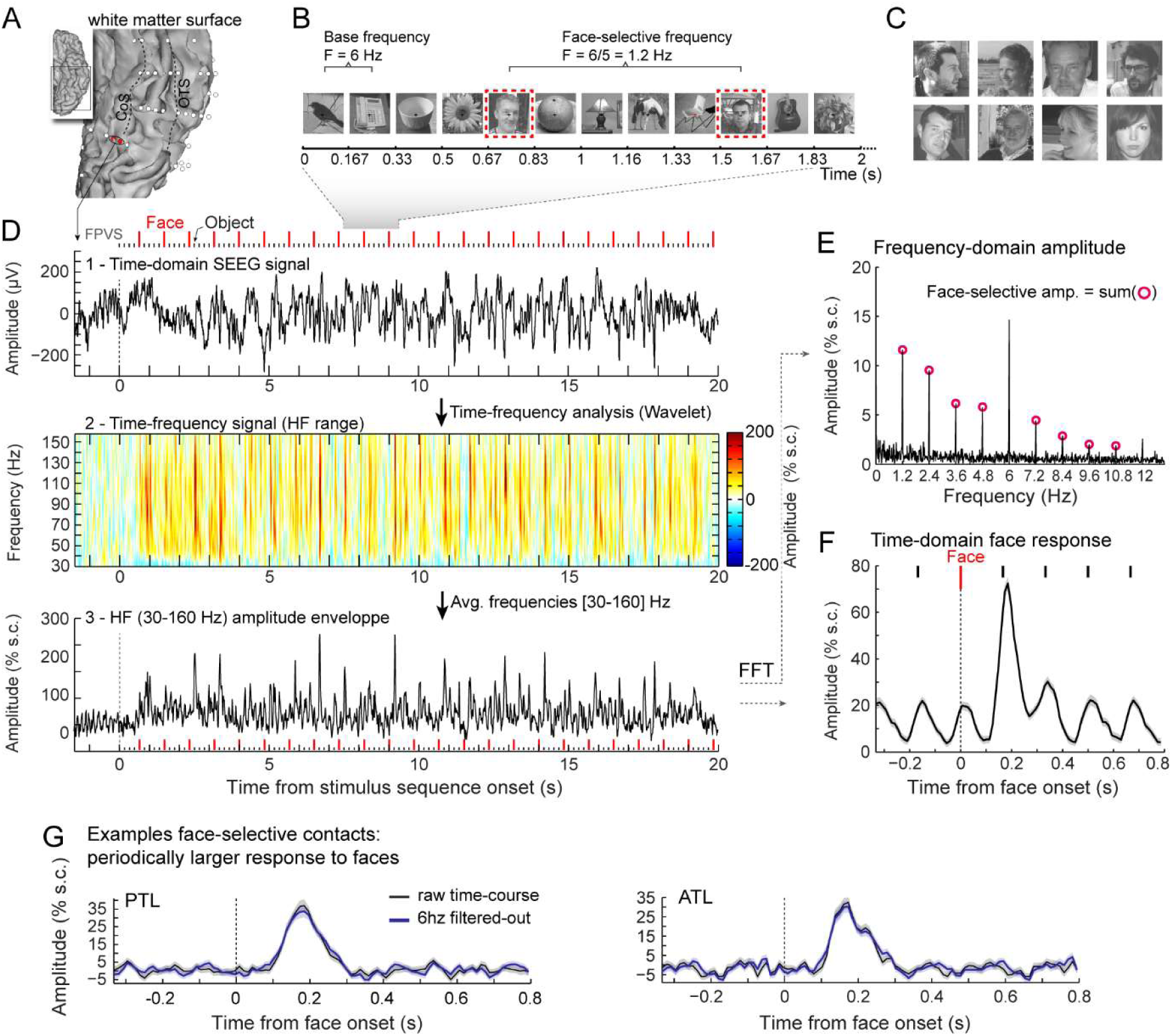
Recording and quantifying time-domain face-selective activity in the VOTC. **A.** SEEG (depth) electrode arrays (white circles) shown on the reconstructed white matter surface of one of the participants (ventral view of the left hemisphere). Electrodes penetrate both gyral and sulcal cortical tissues. **B.** The frequency-tagging paradigm to quantify face-selective neural activity: images of nonface objects appear at a rate of six stimuli per second (6 Hz) with variable face images presented every five stimuli (i.e., every 0.835s). Each stimulation sequence lasts for 65s (2s showed here). **C.** Representative examples of natural face images used in the study (actual images not shown for copyright reasons). **D**. Top: example raw intracranial EEG signal measured at the bipolar recording contacts shown in panel A (red). The signal is shown from -1.5 to 20 s relative to the onset of a stimulation sequence. The time-series displayed is an average of 2 sequences. Above the time-series, red vertical ticks indicate the appearance of face image every 0.835s and small black vertical ticks indicate the appearance of non-face objects every 0.167s. Middle: time by frequency representation of SEEG data in the HFB range (30-160 Hz). The plot shows the percent signal change at each frequency relative to a pre-stimulus baseline period (-1.6s to -0.3s). Periodic burst of HFB activity at the frequency of face stimulation (i.e., 1.2 Hz) are visible. Bottom: modulation of HFB amplitude over time obtained by averaging time-frequency signals across 30-160 Hz. **E.** HFB signal is transformed in the frequency domain to quantify face-selective amplitude as the sum of 12 face-selective frequency harmonics. **F.** Time-domain averaged HFB response to face images shows both the periodic response to non-face objects at 6 HZ (cycle duration = 0.167 s) and the larger face-selective response starting ∼0.1 s after face onset. Shaded area around the curve is standard error of the mean across face trials. **G.** Mean face responses from two separate example recording contacts (in PTL and ATL) in which the 6 Hz response to non-face objects has (black traces) or has not (blue traces) been filtered-out.

Face-selective activity can also be characterized in the time-domain, which allows visualizing periodic activity common to faces and nonface objects (every 0.167 s) and any significant deviation from this activity (i.e., face-selective activity; every 0.833 s; **Figure 1F**). In the time-domain, the face-selective signal can be isolated by filtering-out the 6 Hz signal common to face and non-face objects (i.e., notch filter at 6 Hz and harmonics) (**Figure 1G**) (e.g., (Jacques et al., 2016a; Quek and Rossion, 2017; Retter et al., 2020; Retter and Rossion, 2016; Rossion et al., 2015)).

Visualization of time-courses in contacts with significant face-selective activity reveals that face-selectivity is manifested either as a periodically larger response to faces compared to nonface objects (**Figure 1F,G**, **Figure 2**) or more rarely as a periodically smaller response to faces (**Figure S1**). Since category-selectivity is usually defined as a larger response for a given category compared to control categories in electrophysiological and neuroimaging studies, in the current study we focus on those larger activity for faces. Nevertheless, we provide illustrations for contacts showing response decrease in supplemental material **(Figures S1**). We identified 441 VOTC recording contacts in the gray matter and medial temporal lobe that showed a significant HFB face-selective response increase in 100 participants (among 8278 bipolar contacts located in gray matter or medial temporal lobe in the VOTC of 140 participants; 4680 in the left hemisphere and 3598 in the right hemisphere, **Table S1**). The proportion of face-selective contacts was significantly larger in the right (6.5%, 234/3598) than in the left hemisphere (4.5%, 207/4680, *p* < 0.001, 2-tailed permutation test).

**Figure 2:**
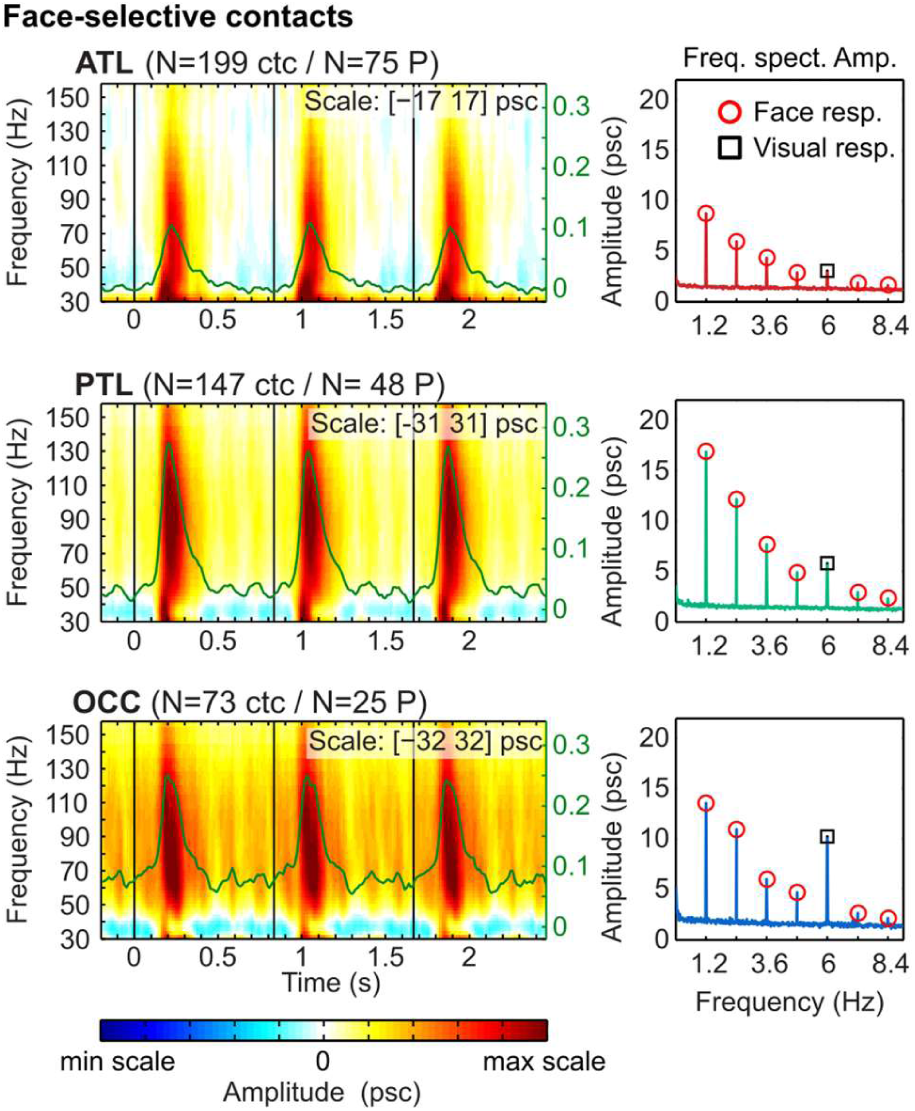
Time-domain face-selective periodic increases. Left column: Time by frequency response (percent signal change, psc; see scale value at the top of each plot, as well as number of recording contacts / participants, and color scale at the bottom) in the HFB range (30-160 Hz) averaged over face-selective contacts in each main VOTC region (OCC: occipital, bottom; PTL: Posterior temporal lobe, middle; ATL: anterior temporal lobe, top). For each recording contact, the time-frequency data was segmented in epochs of about 3 face cycles (i.e. 3 x 0.833 s), averaged by contacts and then averaged over the three groups of contacts. Frequency axis is on the left. Green traces are the HFB amplitude envelope obtained by averaging over the 30-160 Hz range. Amplitude (psc) axis is on the right, show in green. Right column: Frequency spectra averaged over the corresponding groups of recording contacts and showing the face-selective response (red circles) at multiples of 1.2 Hz (i.e., face stimulation frequency) and visual response at 6 Hz (black square, other harmonics not shown). This highlights the sharp decrease in general visual response (i.e., 6Hz) relative to face-selective response from posterior to anterior VOTC.

Face-selective contacts displayed in the Talairach space for group visualization and analyses (**Figure 3A**), indicate that HFB face-selective activity was distributed across the VOTC, from the occipital lobe (OCC) to the anterior temporal lobe (ATL, 95% of contacts located in [-92 to -5] TAL y coordinate, **Figure 3A**), although not reaching the temporal pole (TP). Face-selective activity was also measured in subcortical structures of the medial temporal lobe (MTL, amygdala- AMG). The bulk of the activity stretched from the IOG, through the FG and adjacent sulci (OTS and COS), up to the antFG+ (antFG and surrounding sulci: antOTS and antCOS). The highest number and proportion of face-selective contacts (**Figure 3B,C**, contacts labeled according to each participants individual anatomy as in(Jacques et al., 2022; Jonas et al., 2016a), **Figure S2**) were observed in bilateral latFG and antOTS as well as right IOG.

**Figure 3:**
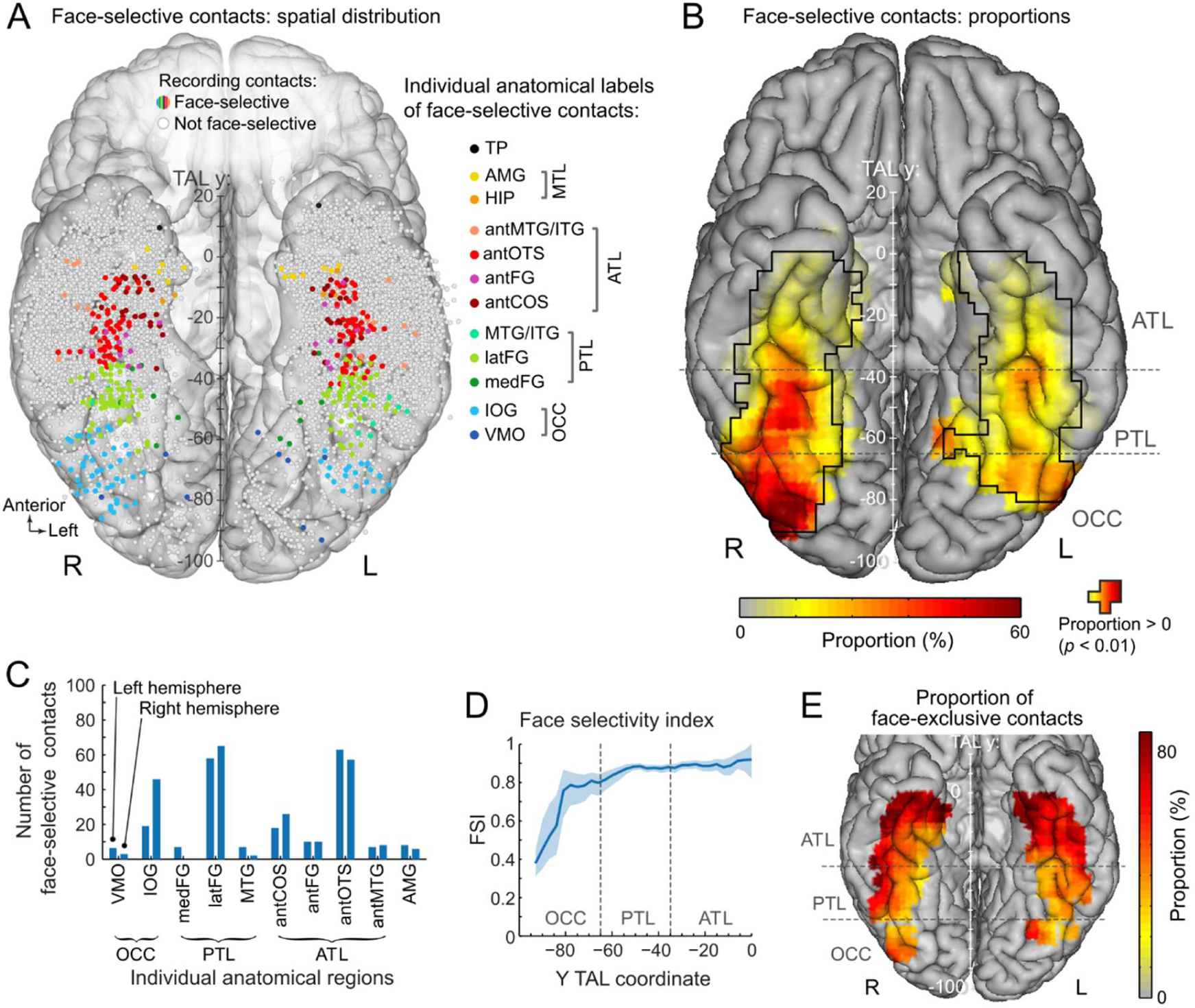
Spatial organization and increase in abstraction of face-selective HFB activity in VOTC. **A**. Map of all VOTC recording contacts across the 140 individual brains displayed in the Talairach space in a transparent reconstructed cortical surface of the Colin27 brain (ventral view). Each circle represents a single recording contact. Face-selective contacts are color-coded according to their anatomical location in the original individual anatomy. White-filled circles correspond to contacts without significant face-selective activity. Values along the y-axis of the Talairach coordinate system (postero-anterior) are shown near the interhemispheric fissure. **B**. VOTC maps of the local proportion of face-selective contacts relative to the number of recorded contacts. Black contour lines delineate local proportions significantly above zero (*p* < 0.01, percentile bootstrap). **C.** The number of face-selective contact is show for each anatomical region (region defined in each individual participant) and hemisphere. **D**. Face-selectivity index (FSI) along the postero-anterior axis collapsed along the X dimension (medio-lateral). The shaded area shows the 99% confidence interval computed using a percentile bootstrap. **E**. Map of the proportion of face-exclusive (i.e., face-selective without significant response to nonface-objects at 6Hz and harmonics) relative to face-selective contacts across VOTC.

### Increase in selectivity (‘abstraction’) from posterior to anterior VOTC

Quantifying separately the amplitude of the face-selective activity (i.e., 1.2Hz and harmonics, excluding harmonics of 6 Hz) and the amplitude the general visual response (i.e., manifested for all presented images, at 6Hz and harmonics) for each contact allows computing an index of face-selectivity (face-selectivity index – FSI) that reflects the magnitude of the face-selective response relative to the overall visual responsiveness of the cortex around the recording contact. The FSI varies from 0 (no face-selective response) to 1 (only face-selective response, no general visual response) and can also be thought of as a proxy for the level of abstraction exhibited by the local neural population (i.e., the degree to which the neural population represents faces independently of generic visual input). FSI increased from posterior VOTC to anterior VOTC: OCC = 0.72, 99% confidence interval: [0.65 0.79]; PTL = 0.88, [0.86 0.89]; ATL = 0.88, [0.86 0.9]; OCC vs. PTL: *p* < 0.0002, 2-tailed bootstrap test; PTL vs. ATL: p = 0.45; OCC vs. ATL: *p* < 0.0002. Along the same lines, computing the FSI as a function of the position along postero-anterior axis (Y Talairach dimension, collapsing contacts over both hemispheres, **Figure 3D**) confirms a strong increase in the FSI from posterior to anterior VOTC (from <0.4 to >0.9), most pronounced over the posterior half of the VOTC. Additionally, we also quantified the proportion of face-selective contacts that are ‘face-exclusive’ (i.e. face-selective contacts that do not display any significant response to non-face objects measured at 6 Hz and harmonics with z < 1.64; *p* > 0.05). Face-exclusive contacts composed 38% of all face-selective contacts (169/441) with a marked increase in proportion from OCC (18%), to PTL (26%) and ATL (51%), reaching ∼80% in the most anterior parts of ATL (**Figure 3E**).

### Concurrent face-selective activity across VOTC

To achieve the most comprehensive and robust possible overview of the time-course of face-selective activity in VOTC, we started by examining the 3 main VOTC regions (i.e., OCC, PTL, ATL, excluding TP and MTL for which there were only few significant contacts), including all significant face-selective contacts showing a response increase (**Figure 3A**). The time-course was characterized by 4 parameters: (1) latency of response onset and (2) offset (**Figure S3**), as well as (3) response overlap and (4) response correlation across VOTC regions (**Figure S4**). The analyses focus on comparing these parameters to specifically test the hierarchical processing hypothesis.

We first considered onset latencies of face-selective signals (**Figure 4**, see **Figure S5** for non-normalized time-series, **Figure S1** for contacts exhibiting face-selective response decrease). Since onset latency estimates can vary substantially as a function of the method used (Rousselet, 2025), we report onset latencies computed with 4 different methods (**Figure S3**): (a) z-score, (b) delta slope, (c) broken stick, (d) percent-of-peak. Methods (a) and (b) rely on statistical thresholds while (c) and (d) do not. To maximize signal to noise ratio and statistical power we collapsed recording contacts across hemispheres (see **Figures S6, S7** and **Table S2** for left and right hemisphere presented separately, **Figure S11** for time-domain responses in individual participants). Statistical analyses were preformed using hierarchical percentile bootstrap (2000 iterations, see methods) and permutation tests (10000 permutations) to account for the partially nested structure of the data set, where each participant potentially contributes more than one contact per region or more than one region. Median onset latencies were extremely similar across VOTC main regions, ranging from 95 ms to 130 ms across regions and methods (**Figure 4A,B, Table 1**). For instance, using the z-score method, onset latencies were 110 ms (95% hierarchical bootstrap confidence interval: [79-134] ms) for OCC (N = 25 participants; 2.9 contacts per participant), 95 ms ([65-118] ms) for PTL (N = 48 participants; 3.1 contacts/p) and 100 ms ([73-132] ms) for ATL (N = 75P; 2.7 contacts/P). Within each method, despite overall differences of latency across methods, the range of median onset latencies across regions was only 15 ms, 8 ms, 9 ms and 12 ms respectively for methods a to d (**Figure 4B**). Statistically comparing regions (OCC vs. PTL, PTL vs. ATL, OCC vs. ATL)revealed no significant differences of onset latency with any of the four methods (all *p*’s > 0.89, two-tailed hierarchical permutation test, fdr-corrected, (Benjamini and Hochberg, 1995), **Table 1**). Effect sizes (Cohen’s d) for all comparisons ranged from 0 to 0.27 (median = 0.055, **Table 1**), indicating very small to small effects. In addition, to determine whether VOTC regions activate concurrently (in parallel) or serially, we performed equivalence testing using the hierarchical bootstrap approach (2,000 iterations). Equivalence bounds and region of practical equivalence (ROPE) were defined to account for conduction delays between regions (i.e., ATL is further away from early visual cortex than OCC) and physiological variability corresponding to a small effect (Cohen’s d = 0.199). Across methods, the median overlap between the bootstrap distributions of differences (between regions) and the ROPE was 85.5%, 85% and 85% for OCC-PTL, PTL-ATL and OCC-ATL respectively, **Table 1**). Moreover, Bayes factor (Cauchy prior with scale = 0.5) were all above 3 (**Table 1**, except for OCC-PTL in the zscore method where PTL onset latency was ahead of OCC), providing moderate to strong support for equivalence across all comparisons (i.e., the measured differences are at least 3 times more likely under equivalence than under difference). These observations support a concurrent rather than a serial activation of VOTC regions, with onset latency differences between regions largely accounted for by simple conduction delays due to increased distance from early visual cortex along the VOTC.

**Figure 4:**
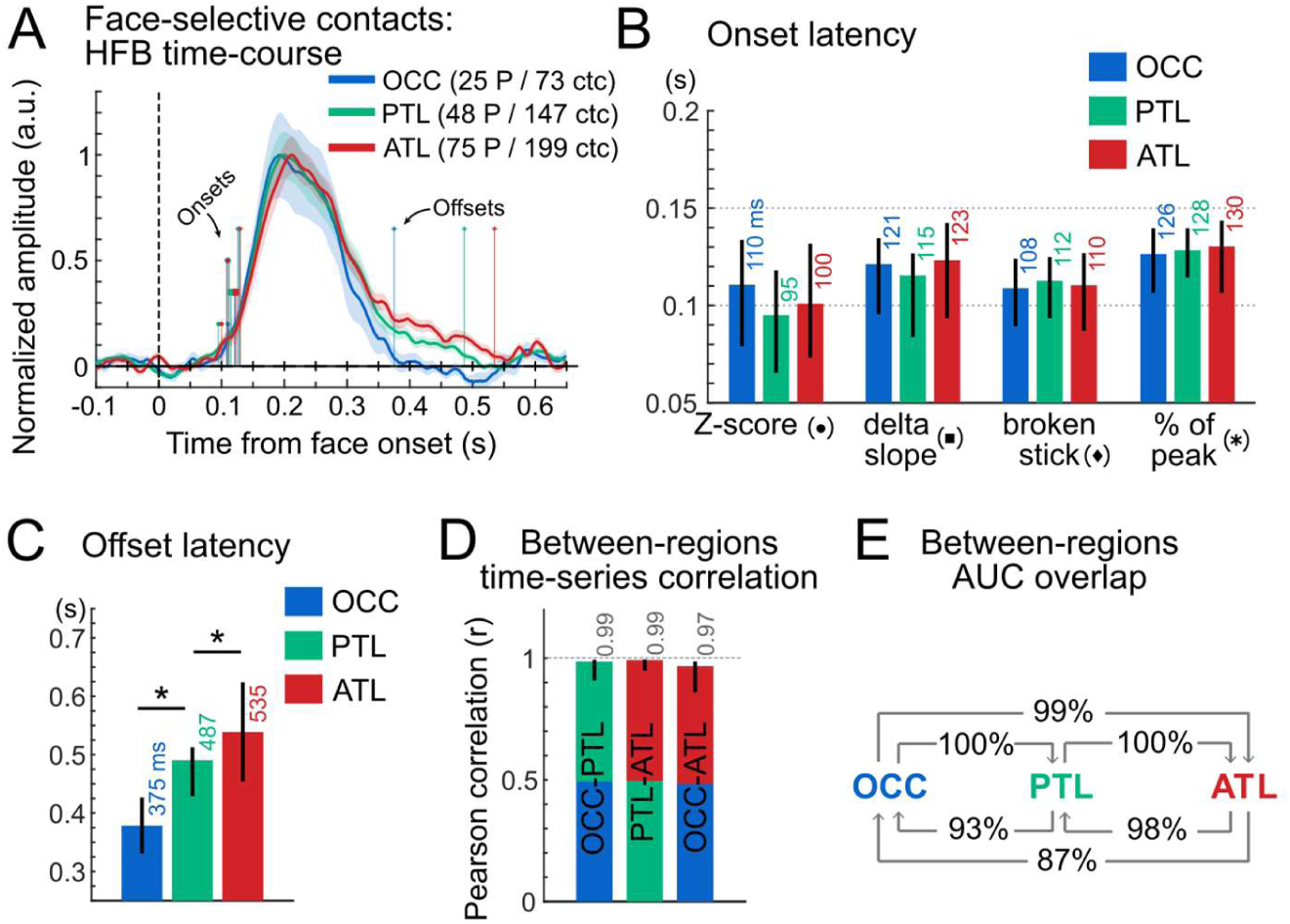
Concurrent face-selective activity across the VOTC. **A.** Mean time-domain face-selective HFB activity in each VOTC region (OCC, PTL, ATL) collapsed across hemisphere. HFB time-series were filtered to remove the general visual response at 6 Hz and harmonics, leaving only face-selective signals. The maximum amplitude of each averaged waveform was normalized to 1 for visualization purposes only (see Figure S5 for non-normalized waveforms). Shaded area represents the standard error of the mean between participants. Colored vertical lines indicate onset latencies across four different methods (see markers’ legend in panel B) and offset latencies (estimated at 5% of peak amplitude) for each VOTC region. **B**. Onset latency for each VOTC main region and for 4 latency estimation methods, together with 95% confidence interval (percentile bootstrap). **C**. Offset latency in each region. **D**. Correlation of time-series between regions and 95% confidence intervals. **E**. Between-region area under the curve (AUC) overlap. The arrow shows the directionality of the computed overlap. For instance, the arrow from ATL to OCC indicates the percentage of the total AUC of ATL (measured between onset and offset latencies) occupied by the AUC of OCC (determined between max(onset(OCC,ATL)) and min(offset(OCC,ATL))).

**Table 1 :**
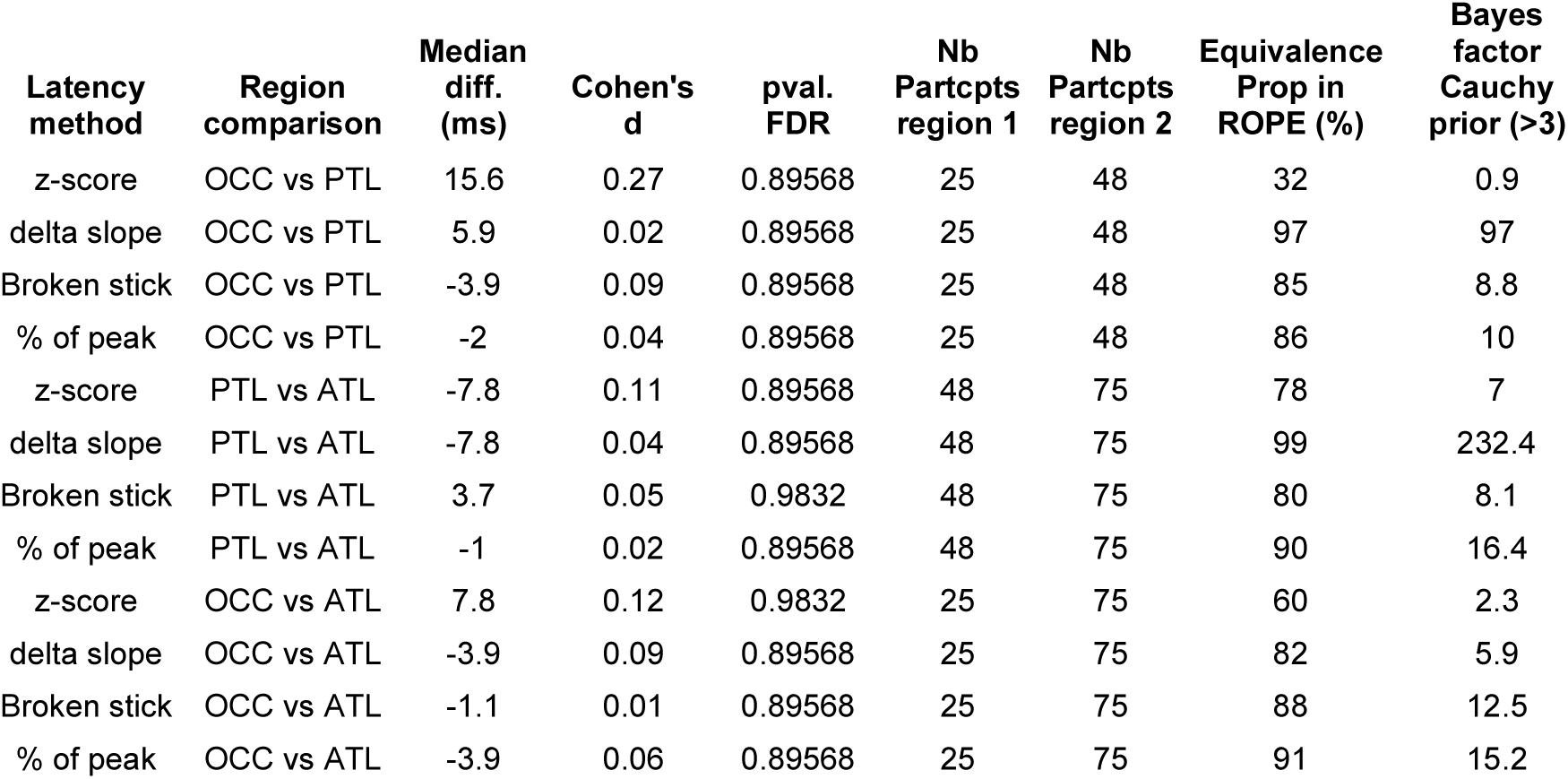
Statistical comparisons of face-selective onset latencies across regions (OCC vs PTL, PTL vs ATL, OCC vs ATL) using permutation tests et equivalence testing (proportion of difference in ROPE and Bayes factor) for 4 different onset estimation methods. Table contains the median latency difference between regions (Median diff.), effect size (Cohen’s d) of the difference, p-value of the permutation test (pval. FDR), number of participants included in each region.

| Latency method | Region comparison | Median diff. (ms) | Cohen's d | pval. FDR | Nb Partcpts region 1 | Nb Partcpts region 2 | Equivalence Prop in ROPE (%) | Bayes factor Cauchy prior (>3) |
| --- | --- | --- | --- | --- | --- | --- | --- | --- |
| z-score | OCC vs PTL | 15.6 | 0.27 | 0.89568 | 25 | 48 | 32 | 0.9 |
| delta slope | OCC vs PTL | 5.9 | 0.02 | 0.89568 | 25 | 48 | 97 | 97 |
| Broken stick | OCC vs PTL | -3.9 | 0.09 | 0.89568 | 25 | 48 | 85 | 8.8 |
| % of peak | OCC vs PTL | -2 | 0.04 | 0.89568 | 25 | 48 | 86 | 10 |
| z-score | PTL vs ATL | -7.8 | 0.11 | 0.89568 | 48 | 75 | 78 | 7 |
| delta slope | PTL vs ATL | -7.8 | 0.04 | 0.89568 | 48 | 75 | 99 | 232.4 |
| Broken stick | PTL vs ATL | 3.7 | 0.05 | 0.9832 | 48 | 75 | 80 | 8.1 |
| % of peak | PTL vs ATL | -1 | 0.02 | 0.89568 | 48 | 75 | 90 | 16.4 |
| z-score | OCC vs ATL | 7.8 | 0.12 | 0.9832 | 25 | 75 | 60 | 2.3 |
| delta slope | OCC vs ATL | -3.9 | 0.09 | 0.89568 | 25 | 75 | 82 | 5.9 |
| Broken stick | OCC vs ATL | -1.1 | 0.01 | 0.89568 | 25 | 75 | 88 | 12.5 |
| % of peak | OCC vs ATL | -3.9 | 0.06 | 0.89568 | 25 | 75 | 91 | 15.2 |

Second, estimating the latency at which neural activity returned to baseline level (i.e., offset latency) with the percent-of-peak method (i.e. when does the signal goes back to under 5% of the baseline-to-peak amplitude), indicate a gradual increase in the offset latency (**Figure 4A,C**) from OCC (375ms, [331-427] ms) to PTL (487 ms, [429-513] ms) and ATL (535 ms, [454-624] ms), with all statistical comparisons across the 3 regions reaching significance (*p* = 0.026, two-tailed permutation test, fdr-corrected). Together with the similar onset latencies across main VOTC regions, this indicates that overall, response duration increases as a function of postero-anterior VOTC location.

Third, we examined the temporal overlap between two given regions by computing the percentage of the area under the response curve (AUC) of the overlap between region A and B (determined using onset and offset latencies) relative to the total AUC of region A (see methods, **Figure S4**). Interestingly, despite the differences in offset latencies across VOTC, on average when comparing two regions, the AUC that falls within the temporal overlap between these regions represents 96% of the total AUC for each region, ranging from 87% (percentage of the ATL AUC occupied by the AUC of its temporal overlap with the OCC, see **Figure 4E**) to 100% (e.g., percentage of the PTL AUC occupied by the AUC of its temporal overlap with the ATL, **Figure 4E**). This indicates that, even in case of relatively large differences in offset latency (e.g., 375 ms for OCC and 534 ms for ATL), the portion of the ATL AUC during non-overlapping window (e.g., from 126 to 130 ms and 375 to 535 ms for OCC vs. ATL) is negligible (13%) compared to the AUC observed during the temporal overlap (130 ms to 375 ms, 87% of the ATL AUC occupied by the AUC of its temporal overlap with the OCC). Moreover, these observations indicate that the vast majority of the response in a region located anteriorly (e.g., ATL) to another region (e.g., PTL) takes place during the time-window of the posterior region.

Fourth, as an additional measure of temporal overlap we considered the correlations between the mean time-series recorded in each region, which were highly correlated (**Figure 4D**), ranging from Pearson’s r=0.96 when comparing OCC to ATL, to r=0.99 when comparing PTL to ATL Importantly, statistically comparing the correlations calculated between-regions (computed using random halves of the data to match sample size of within-region correlations) to correlations *within* each region (using split halves of the data, within OCC: r = 0.95; within PTL: r = 0.99; within ATL: r = 0.98) revealed no significant difference (*p*’s > 0.35, one-tailed percentile boostrap test, fdr-corrected). This indicates that the time-domain neural signals measured across different VOTC regions are not significantly more different than signals measured from the same region.

In a separate analysis (**Figure S8**), we also fully replicate the above observations but focusing on timing properties in 3 core face-selective regions in VOTC: IOG, latFG and antFG+ (the latter being composed of antFG, antOTS, antCOS). In addition, timing analyses performed on low frequency event-related potentials (ERPs) confirms that beyond early visual retinotopic cortex (V1-hV4, posterior to the transverse Collateral Sulcus, (Brewer et al., 2005; Winawer and Witthoft, 2015)) where responses start up to 40 ms before responses measured in contiguous anterior regions, concurrent face-selective activity is measured across the whole VOTC (**Figure S9**, **Table S3**).

In addition to the group analyses described above, we performed a within-participants comparison of onset latencies in participants that had face-selective contacts in at least 2 different main VOTC regions (e.g., OCC and PTL). While within-participants statistics allow disregarding between-participants variability, it was done here at the expense of sample size since only a small subset of participants have face-selective contacts located in multiple face-selective regions. Onset latencies were computed for individual recording contacts and averaged by region within participants. These comparisons did not reveal significant differences in onset latencies between main VOTC regions collapsed across hemispheres, irrespective of the methods used to estimate onset latencies (OCC minus PTL: N=13, median difference across methods = 0 ms, all *ps* > 0.25, 2-tailed permutation test, fdr-corrected; PTL minus ATL, N=19, median difference across methods = -17 ms, *ps* > 0.05; OCC minus ATL, N=9, median difference across methods = -19 ms, all *ps* > 0.11). Furthermore, very small effect size (OCC minus PTL, median Cohen’d across methods = 0.02) and small-to-medium effect sizes for OCC minus ATL (median Cohen’d across methods = 0.4) and PTL minus ATL (median Cohen’d across methods = 0.57) were observed. Nevertheless, these effects were more compatible with equivalence than difference when accounting for conduction delay between regions (see above). Across methods, the median overlap between the distributions of differences (between regions) and the ROPE was 75%, 73% and 81% for OCC-PTL, PTL-ATL and OCC-ATL respectively. Similarly, median Bayes factor (Cauchy prior with scale = 0.5) were all above 3 (OCC-PTL = 8.5; PTL-ATL = 3.6; and OCC-ATL = 6.2), providing moderate to strong support for equivalence across all comparisons.

### Concurrent functional connectivity between face-selective regions

Last, we estimated the functional connectivity between pairs of main face-selective regions (IOG, latFG, antFG+) by correlating single-trial face-selective amplitude measured in different VOTC regions (IOG, latFG, antFG+), across time. Our use of local bipolar referencing of electrophysiological signals ensures that responses in each region are maximally local and that any significant correlation between regions cannot be attributed to the reference itself. Directionality in connectivity (e.g., connectivity from IOG to latFG or the reverse) was estimated by correlating face-selective amplitude measured at different time-lags (-150 to 150ms) between the compared regions (Kadipasaoglu et al., 2017). While we used Pearson correlations as in (Kadipasaoglu et al., 2017) for the main analysis, examining the coupling between regions using mutual information (using the method in (Ince et al., 2017)) which is sensitive to both linear and non-linear relationship, yielded a very similar pattern of results (**Figure S10**). The properties of the functional connectivity patterns were largely similar across pairs or regions (**Figure 5**), although correlations were weaker between IOG and antFG (**Figure 5**). Overall, correlations were present before response onset, manifested by weak scattered yet largely symmetrical small correlation clusters. Stronger, more consistent and clustered between-region functional connectivity appeared at response onset, i.e., around 100-120ms, and was maximal at response peak between 150 and 200 ms (see **Figure 4A**), remaining significant for the remaining of the response. Most importantly, correlations measured after response onset were consistently centered on the diagonal (0 ms lag), further supporting concurrent activity of the VOTC regions and suggesting that these regions receive common correlated concurrent inputs possibly from lower-level visual cortex. Despite correlations being centered on the diagonal, correlation coefficients symmetrically spread around the diagonal, resulting either from reciprocal connections or from short (<10-30ms, likely due to temporal smoothing caused by the time-frequency analyses) or longer (up to 150ms) duration of temporal auto-correlation of the electrophysiological signal. These broader/longer correlations around the diagonal, which were most pronounced between IOG-latFG and latFG-antFG (**Figure 5**, left and middle), appeared after 200 ms and were likely related to sustained and stable face-selective neural responses at these latencies which are correlated in amplitude across regions.

**Figure 5:**
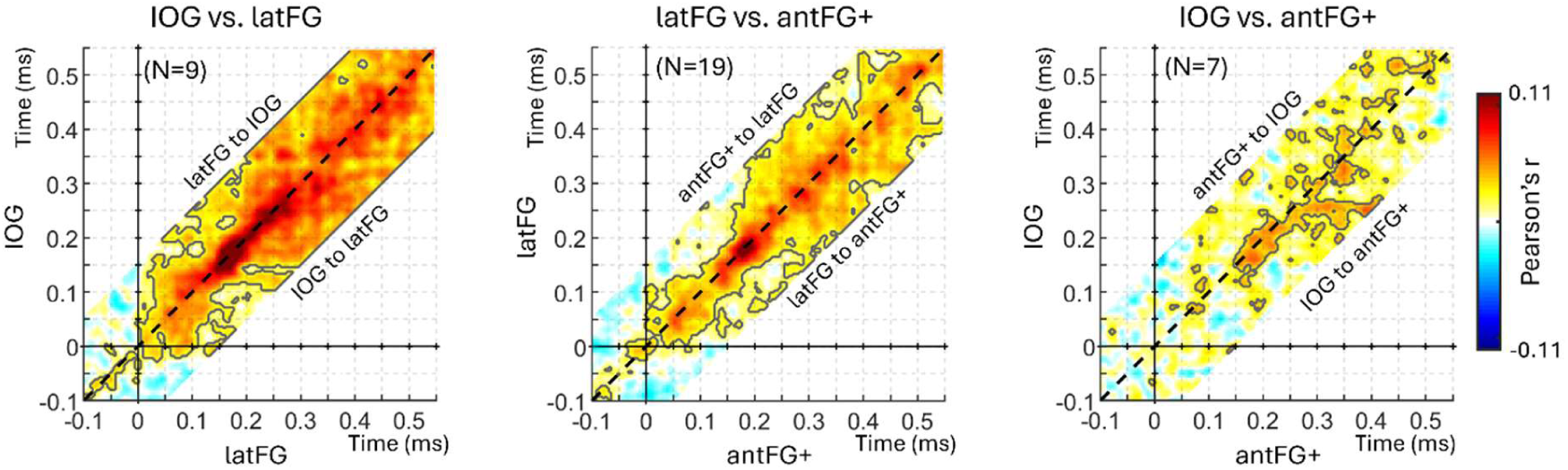
Concurrent functional connectivity between face-selective regions. Each plot shows the group-averaged Pearson’s correlations between single-trials face-selective amplitude measured at two distinct face-selective regions (left: IOG and latFG; middle: latFG and antFG+; right: IOG and antFG+) computed at each time point, representing functional connectivity between pairs of regions. Black contour lines indicate significant positive correlations (*p* < 0.01, fdr-corrected). Correlations were computed across -150 to 150 ms lags between regions to infer direction of connectivity. The black dashed diagonal line represents a 0 ms time-lag between regions. Correlations centered above the diagonal would indicate that face-selective activity in the more anterior region (e.g., latFG in left panel) correlates with but precedes activity in the posterior region (e.g., IOG in left panel), suggesting an information flow from anterior to posterior, and the reverse for correlations centered below the diagonal.

### Mapping face-selective onset latency in Talairach space

Next, we aimed to explore the onset latencies of face-selectivity HFB signal at a more local scale, taking advantage of the dense sampling of the VOTC to compute maps in the Talairach space. Again, this was computed by collapsing across the left and right hemispheres to increase sample size. Moreover, we only included local VOTC volumes (i.e., ‘voxels’) with at least 10 contacts. While overall onset latencies varied between methods, the pattern of latencies across VOTC were similar (**Figure 6A**). Specifically, face-selective response latency was lowest in regions exhibiting the most consistent face-selectivity: IOG (z-score method: IOG: mean = 94 ms, range = [60 - 108ms], **Figure 6A**) and laFG (mean = 97 ms, range = [91 - 116ms]), as well as the posterior part of antFG+ area (y-coordinates between -40 and -20 mm; mean = 94 ms, range = [61 - 106ms]). Between and around those regions, latency was slightly higher (e.g.: anterior part of antFG+: mean = 126 ms, range = [75 – 138ms]; posterior FG: mean = 113 ms, range = [102 - 130ms]), especially in the antMTG region (mean = 137 ms, range = [100 - 157ms]). Examining the postero-anterior profile of face-selective response latency (**Figure 6B**) reveals relatively constant latencies (∼90 to 120ms depending on the method) in the OCC (Talairach Y coordinates: -90 to -65 mm]), PTL (Talairach Y coordinates: -65 to -40 mm) and posterior portion of ATL (Y < -20 mm), after which the response latency increased(i.e., 15-35 ms increase). However, this increase was within the bounds of what would be expected by chance if the antero-posterior position had no influence on face-selective response onset latency (**Figure 6B**, randomization test where the pairing between each recording contact location and its time-domain response was shuffled), and accounting for the expected conduction delays between posterior and anterior VOTC (distance > 60 mm, axonal conduction speed likely between 1.7 and 5.3 m/s (Lemarechal et al., 2022; van Blooijs et al., 2023), **Figure 6B**), even with direct cortico-cortical connections from earlier visual cortical areas.

**Figure 6:**
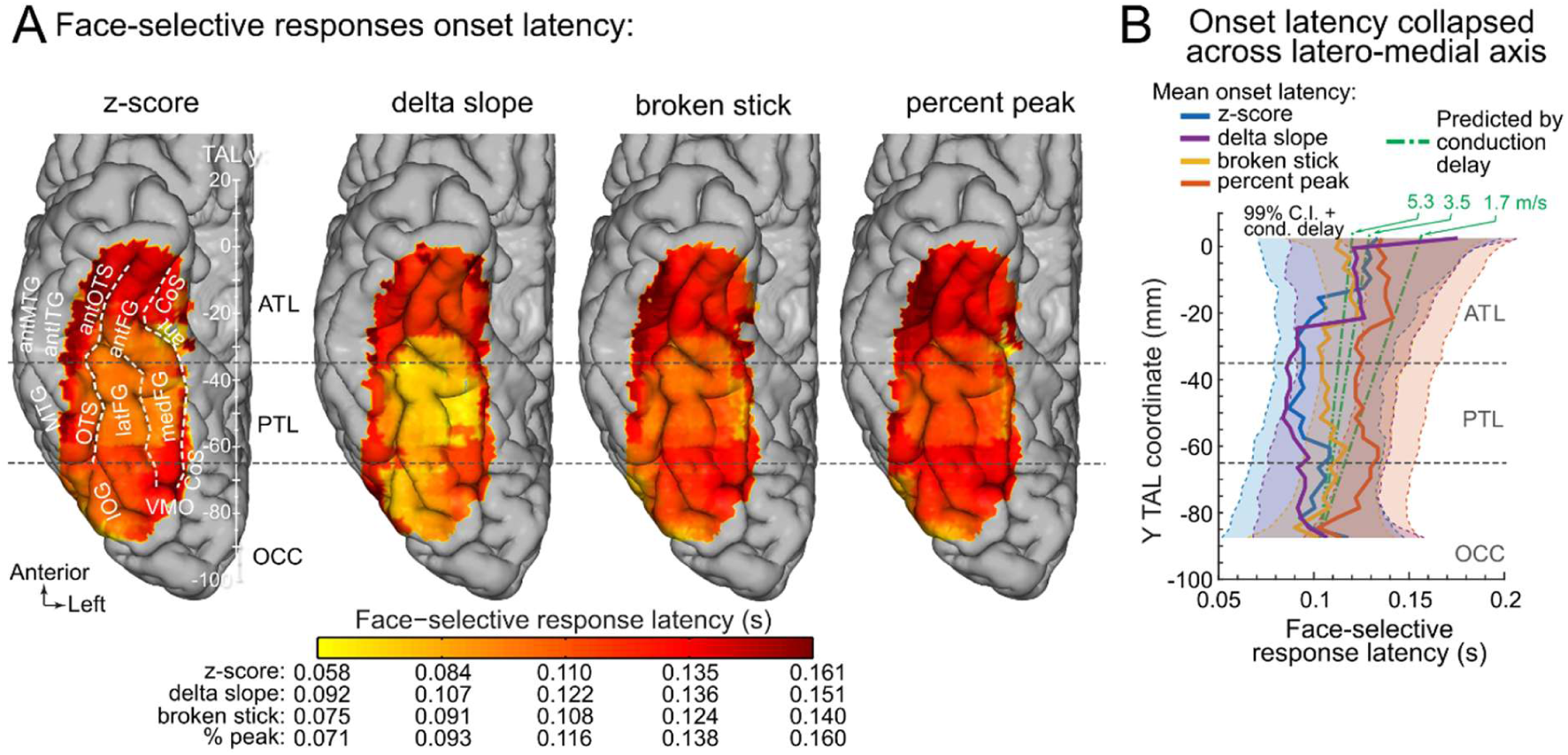
Mapping concurrent face-selective response onset latency in VOTC. **A**. Face-selective onset latency map across VOTC, collapsed across hemispheres and displayed for 4 different methods to estimate response onset latency (see methods). **B.** Variation of face-selective response latency along the postero-anterior axis for 4 estimation methods. For each method, each data point represents the onset latency measured from the time-series averaged over contacts collapsed across the medio-lateral X dimension within 20 mm segments (in the Y dimension). Thick lines are estimated onset latencies and shaded areas show the 99% confidence intervals expected under the null hypothesis that the postero-anterior location has no influence on the onset latency, accounting for the expected conduction delays between posterior and anterior VOTC. Green lines show the expected increase in response onset latency based on simple axonal conduction delays (Lemarechal et al., 2022; van Blooijs et al., 2023) due to increasing distance from the occipital region, with reference to the latency averaged over the OCC region for the ‘zscore’ method. Lines show mean expected conduction velocity for direct cortico-cortical connections (∼3.5 m/s) and +/- 1 std (1.7 and 5.3 m/s).

## Discussion

Here we provide a large-scale mapping of the time-course of neural activity supporting visual recognition in the human VOTC with electrophysiological intracerebral recordings. Faces are used as the key target category for visual recognition not only because of their highly significant social value for humans, but also due to their wide distribution of representation across the (bilateral) VOTC, allowing to compare the time-course of category-selectivity across multiple regions. In addition, the cortical network of face-selective regions is the most extensively studied within the occipital and temporal lobes and is thought to reflect the manifestation of a canonical set of operations that reveal general principles of how primate visual recognition works (Connor, 2010; Conway, 2018; DiCarlo et al., 2012; Freiwald, 2020; Grill-Spector et al., 2017) and is implemented in artificial visual recognition systems (i.e., deep neural networks; e.g., (Grill-Spector et al., 2018).

We report two major findings. First, a progressive increase in face-selectivity along the postero-anterior axis. Second, a concurrent face-selective neural activity, with similar onset latencies (∼100ms) and largely overlapping and correlated time courses across the whole VOTC, spanning around 90 mm of cortical territory along the postero-anterior axis. Our observations suggest that while the human VOTC may apparently be functionally organized in a hierarchy of regions with increasingly more abstract representations, these regions of the association cortex do not appear to be activated sequentially (or even in cascade) but rather concurrently.

### 1. Time course of VOTC face-selective activity and speed of face categorization

The onset of category-selectivity for faces identified here in humans at around 100 ms at the group level is early, supporting an initial fast feedforward sweep of activity for rapid face categorization (Cauchoix et al., 2014; Crouzet and Thorpe, 2010; Retter et al., 2020; Serre et al., 2007). This onset latency, with a peak of activity reached at about 200 ms, is in line with multiple sources of evidence (speed of gaze to lateralized natural images of faces(Crouzet and Thorpe, 2010); scalp EEG studies (Retter and Rossion, 2016; Retter et al., 2020; Rossion and Caharel, 2011; Rousselet et al., 2008); intracranial EEG studies(Allison et al., 1999; Kadipasaoglu et al., 2017; Liu et al., 2009); single neuron activity in human face-selective fusiform regions (Laurent et al., 2025; Quian Quiroga et al., 2023)). Here we extend these findings by providing comprehensive measurements of response onset latencies across the whole VOTC. Importantly, we do not measure the absolute response onset latency to face images (Cao et al., 2025; Jacques et al., 2016b; Regev et al., 2018; Schrouff et al., 2020). Although these are generally similar to our measure of face-selective onset latencies in these regions, investigating category-selective activity is key because it may be temporally dissociated from general visual responses in the same brain areas (Jiang et al., 2011) and is critically related to visual recognition function (Jonas and Rossion, 2021; Rangarajan et al., 2014; Volfart et al., 2022).

Beyond onset latencies, we also report response durations that are in line with values obtained from the scalp (EEG) with the same paradigm (about 420ms (Retter and Rossion, 2016; Retter et al., 2020; Rossion et al., 2015)). Despite a slight increase in duration of the sustained low amplitude part of the time-series along postero-anterior VOTC axis, the overwhelming majority of signal amplitude (87 to 100%) and variability (r = 0.97 to 0.99) occurs concurrently, i.e., it temporally overlaps across brain regions. Given the timing similarity between the present intracerebral data and scalp EEG data obtained in neurotypical individuals with the same approach (Retter and Rossion, 2016; Rossion et al., 2015), there appears to be no delay potentially attributable to the specific population tested here, i.e., patients with intractable refractory epilepsy (Allison et al., 1999). While we cannot exclude overall lower response amplitudes of face-selective activity in this specific population relative to normal individuals and/or potential shifts of hemispheric lateralization at the individual level depending on hemispheric seizure localization, this would not affect our conclusions. Finally, whereas face-selectivity is measured here during temporally fixed face stimulation (every 833 ms), observers perform an orthogonal task, are unaware of the periodicity of face stimulation and unable to determine it due to the rapid base stimulation rate. Most importantly, with this fast stimulation mode approach, both the amplitude and time course of face-selective activity are unaffected by the temporal distance between face stimuli/ratio of faces *vs.* objects (Retter & Rossion, 2016), or the temporal periodicity or predictability of face presentation (i.e., identical time-courses and amplitudes of face-selective activity for periodically or nonperiodically embedded faces in nonface object trains; (Quek and Rossion, 2017))

### 2. A progressive posterior to anterior gradient of category-selective abstraction in the VOTC

Our findings indicate a progressive increase in category-selectivity from posterior to anterior VOTC. More specifically, while category-selective populations of neurons in posterior VOTC regions tend to respond to all visual stimuli, as evidenced by high 6 Hz amplitude, this common visual response gradually decreases along the postero-anterior axis (**Figures 2**, **3D**), leading to a high proportion of exclusive response to faces in the most anterior regions of the ventral temporal lobe (**Figure 3E**). The postero-anterior increase in face-selectivity is only partially in line with fMRI observations, which have reported both larger face-selectivity in the fusiform gyrus (‘Fusiform Face Area’, FFA) than in the posteriorly located inferior occipital gyrus (IOG; ‘Occipital Face Area’, OFA)(Tsao et al., 2008; Weiner and Grill-Spector, 2010), but also opposite findings (Rossion et al., 2012) (see also recent evidence (Chen et al., 2023) of larger face-selectivity in the posterior than middle fusiform gyrus), potentially depending on the general size/amplitude of activity in a given ROI and how selectivity indexes are computed. Most importantly, fMRI studies are limited to comprehensively explore the VOTC due to the large magnetic susceptibility artifacts (low SNR) in the ventral ATL (Ojemann et al., 1997; Rossion et al., 2024). To the best of our knowledge, previous intracranial studies have not described such a comprehensive pattern of variation in face-selectivity across the whole VOTC, due to limited sampling (spatially or limited to the gyral surface), lack of mapping and/or face-selective response quantification, or focus on medio-lateral variations of selectivity (Allison et al., 1999; Engell and McCarthy, 2014; Jacques et al., 2016b; Jonas et al., 2016a; Kadipasaoglu et al., 2017, 2016; Rangarajan et al., 2014; Schrouff et al., 2020; Vidal et al., 2010).

This increase in selectivity for faces, as well as the increase in the proportion of face-exclusive activity in the postero-anterior VOTC axis, cannot be attributed to a mere reduced ability of anterior regions to generate responses at a fast presentation rate (6 Hz) (Jonas et al., 2016a). Rather, these observations suggest an increase in ‘abstraction’ of visual neural representations, with a large proportion of neuronal populations in anterior VOTC regions exhibiting similar activity to different faces images independently of the context in which they appear, while being much less (or not at all) responsive to other complex visual stimulation. These findings are compatible with standard hierarchical models of visual processing, such as proposed originally by (Hubel and Wiesel, 1962) and extended to a hierarchy of stages for building increasingly complex invariant visual object or face representations (Connor, 2010; Conway, 2018; DiCarlo et al., 2012; Duchaine and Yovel, 2015; Fairhall and Ishai, 2007; Freiwald, 2020; Freiwald & tsao, 2010; Grill-Spector et al., 2017; Grill-Spector et al., 2018; Issa et al., 2018; Marr, 1982; Riesenhuber and Poggio, 1999; Tsao, 2014; Van Essen et al., 1992), as well as with the view that this hierarchy is implemented anatomically along the postero-anterior axis of the human VOTC.

### 3. Concurrent face-selectivity across VOTC: evidence for non-hierarchical organization of visual association cortex

While the increase in category-selectivity from posterior to anterior VOTC as described above is compatible with a hierarchical model of visual recognition, the remarkably similar time-course of face-selective visual neural activity across human VOTC regions is difficult to reconcile with the temporal properties implied by such model. According to the classical hierarchical view of visual processing (Conway, 2018; DiCarlo et al., 2012; Duchaine and Yovel, 2015; Fairhall and Ishai, 2007; Freiwald, 2020; Grill-Spector et al., 2018 Grill-Spector et al., 2017; Issa et al., 2018; Marr, 1982; Riesenhuber and Poggio, 1999; Tsao, 2014; Van Essen et al., 1992), representations at a given level of the hierarchy are built from combination of inputs received from a lower level, implying serial ordering and thus a time difference between successive levels (Bullier, 2001; Bullier and Nowak, 1995; DiCarlo et al., 2012; Freiwald & Tsao, 2010; Schmolesky et al., 1998; Thorpe and Fabre-Thorpe, 2001).

Hierarchical models do not imply that a process should be completed at a given stage before the next one is initiated. Yet, even in a hierarchical cascade of cortical processes and representations (DiCarlo et al., 2012), and regardless of acknowledged feedback/reentrant loops (Edelman and Gally, 2013; Issa et al., 2018; Kar et al., 2019; Kravitz et al., 2013; Lamme and Roelfsema, 2000) there should be delays in onset times, and only partial temporal overlap between increasing processing level of the hierarchy(Bullier and Nowak, 1995; Freiwald & Tsao, 2010; Issa et al., 2018; Schmolesky et al., 1998). Based on findings from electrophysiological as well as cortico-cortical evoked potential (CCEP) measurements, the delay between each level should be at least 10-30ms depending on (multi)synaptic transmission delays and conduction delays in short- or long-range connections(Conner et al., 2011; Freiwald & Tsao, 2010; Lemarechal et al., 2022; Nowak and Bullier, 1997; Schmolesky et al., 1998; van Blooijs et al., 2023). Importantly, this standard view of hierarchical processing is also how human, and non-human primate, cortical recognition of faces is generally conceived (Connor, 2010; Conway, 2018; Duchaine and Yovel, 2015; Fairhall and Ishai, 2007; Grill-Spector et al., 2017; Issa et al., 2018; Schweinberger and Neumann, 2016; Tsao, 2014), with up to 50 ms time difference postulated between ‘stages’ in humans (Sadeh et al., 2010; Schweinberger and Neumann, 2016), and computationally modeled (e.g., deep neural networks; Grill-Spector et al., 2018).

In contrast to this hierarchical view, the present findings of concurrent face-selective activity across the human VOTC support a non-hierarchical organization of this recognition process. More specifically, our observations suggest that, rather than being successively activated along a posterior-axis, multiple face-selective (clustered) populations of neurons spread across the whole VOTC receive direct and parallel sensory inputs from low-level retinotopic visual areas (e.g., V1-hV4), through so-called bypass pathways. Such pathways have long been described in anatomical studies of the visual system of macaque monkeys (Conway, 2018; Distler et al., 1993; Eldridge et al., 2023; Van Essen and Maunsell, 1983; Zeki and Shipp, 1988;). In humans, specifically for face recognition, both lesion studies and time-resolved fMRI investigations have provided indirect evidence for bypass cortical pathways, reporting face-selective activity in the midFusiform gyrus (i.e., ‘Fusiform face Area’, FFA) in the absence of (lesion: (Rossion et al., 2003; Steeves et al., 2006; Weiner et al., 2016)), or before (Jiang et al., 2011), any category-selective activity in the posteriorly located IOG. Moreover, while an intracranial recording study with a limited sample focusing largely on occipital and posterior temporal face-selective regions has provided mixed outcomes regarding the onset of selective response to faces (delayed onset in the left latFG relative to IOG in n=4; no delay in the right hemisphere n=3), CCEP in the same study(Kadipasaoglu et al., 2017) also suggested independent and parallel signal propagations between early visual areas and both face-selective regions (i.e. latFG/FFA and IOG). Critically, this study did not report face-selective onset latencies in the ATL, where ∼200 face-selective contacts were measured in the current study. Further evidence from Diffusion Tensor Imaging (DTI) suggest independent connections between early visual cortex and face-selective regions in the IOG and post/mid-fusiform gyrus(Bryant et al., 2019; Finzi et al., 2021; Kim et al., 2006; Wang et al., 2020; Weiner et al., 2016). More generally, human V1 even appears to show (hominoid-specific) connectivity with lateral, inferior, and anterior temporal regions, beyond the retinotopically organized cortical areas(Bryant et al., 2019).

While these observations provide an anatomical basis for concurrent onset times in face-selective regions of the human VOTC, the present electrophysiological data, collected from a particularly large sample and with an objective face-selective measure, go beyond by showing for the first time that (1) even the most anterior temporal population of neurons (ATL) respond selectively to faces as early as posterior VOTC regions (taking into account minimal fiber conduction delays) and (2) that all of these regions’ time-courses overlap in time remarkably. The current findings also support direct connections between early visual cortex and face-selective regions around the anterior fusiform gyrus, anterior OTS and anterior COS (antFG+) in the ATL (likely via the same fiber bundles as for posterior face-selective regions: the inferior longitudinal fasciculus and inferior fronto-occipital fasciculus) where DTI measurements are limited by magnetic susceptibility artifacts. Importantly, while we report a slight overall increase in response onset latency in the ATL (e.g., **Figure 6B**) compared to more posterior regions, this increase can be accounted for by the predicted conduction delay given the large distance of the ATL from early visual areas in the posterior occipital cortex and the likely reliance on long range white matter fibers (∼1.7 to 5.3 m/s or mm/ms (Lemarechal et al., 2022; van Blooijs et al., 2023)). This further supports the view that there is no need to postulate mandatory intermediate hierarchical stage/relays between posterior and anterior regions. A number of factors can affect transmission speed, as determined using CCEP in humans, among which the number of fibers connecting different regions(Conner et al., 2011) or the myelin density of these connections (Bryant et al., 2019; Glasser and van Essen, 2011), which are likely to vary between face-selective regions (Finzi et al., 2021; Natu et al., 2019; Wang et al., 2020). Together with concurrent onset latencies, the large overlap in time-courses suggests parallel category-selective processing, likely to be initiated independently by low-level sensory inputs from early visual cortex, as well as ongoing reentrant interactions (i.e., dynamic recursive/recurrent exchange of signals (Edelman and Gally, 2013; Kar et al., 2019; Lamme and Roelfsema, 2000)) between these regions for ∼300ms(Issa et al., 2018; Kadipasaoglu et al., 2017). Moreover, our finding of differences across face-selective regions in the duration of the low amplitude sustained part of the response may reflect different patterns of connectivity between these regions and other (sub)cortical regions to which they are connected to. In particular, the longest duration in the ATL may potentially be linked to the specific connectivity of this region with medial temporal lobe and other structures involved in semantic or episodic memory formation or retrieval (Persichetti et al., 2021).

### 4. Implications for models of visual recognition

How would sensory inputs be recognized as faces in such a non-hierarchical human system? In a nutshell, through synaptic low-level sensory inputs originating from early visual cortex (e.g., V1) successfully triggering (post-synaptic) activity of sufficiently large face-selective clustered populations of neurons in parallel throughout the VOTC. Note that despite temporal overlap, and concurrent process, there are genuine differences in degree of selectivity, as disclosed here, as well as receptive fields sizes, foveal bias and ipsilateral sensitivity across face-selective human VOTC regions (Finzi et al., 2021; Kay et al., 2015). These functional differences between regions can be advantageous for visual recognition since they increase the probability of variable incoming sensory inputs from early visual cortices to successfully trigger clustered face-selective populations of neurons (‘patches’) interconnected as chains (Gordon et al., 2026) along the VOTC. Importantly, in such a non-hierarchical system, there is no requirement for normalization processes across definite successive stages to build ‘invariant’ representations from inputs varying in size, orientation, lighting, etc. Instead, through hebbian learning mechanisms (Fuster, 1995; Hebb, 1949; Markram et al., 2011), views that are more commonly experienced (e.g., full-front faces at conversational distances) would be represented by larger populations of neurons than rarely seen views, providing a faster accumulation of category-selective neural activity (i.e., evidence accumulation) in this network and a more efficient recognition function for sensory inputs corresponding to these views (Perrett et al., 1998).

To be clear, such a non-hierarchical organization of the human ventral cortical face network corresponds fully to a feedforward/bottom-up view of visual recognition (at least for recognition of relatively clear views of faces) enriched by concurrent reentrant exchanges of signals within VOTC regions for a few hundreds of milliseconds. That is, putative ‘top-down’ signals from population of neurons in parietal or prefrontal cortices(Bar, 2003; Kar and DiCarlo, 2021) are not necessary for fast automatic recognition of faces. Yet, this simple view rests on a conceptual reinterpretation of functional organization of the human VOTC: rather than constituting a series of processing stages with increasing levels of complexity to derive visual representations (i.e., build the most veridical images of the world), face-selective populations of neurons are conceptualized as memory representations (‘cortical memories’ (Fuster, 1995)) distributed throughout the VOTC. That is, these clustered populations of neurons, constrained by cytoarchitecture-specific white matter connections from birth (Kubota et al., 2025; Mahon, 2022), have learned (from early developmental stages(Kosakowski et al., 2022)) through Hebbian mechanisms to discharge selectively to faces, with temporal synchrony strengthening their connections (Hebb, 1949; Markram et al., 2011). Recognition, which ‘simply’ involves successful – (often) concurrent - matching of sensory inputs to these cortical memories in the human association cortex is achieved in a bottom-up fashion, with the ‘up’ (i.e., memory) component being even critical in recognizing ambiguous or degraded sensory inputs as faces (Cavanagh, 1991).

#### Summary and conclusions

In summary, our large-scale human intracerebral investigation of category-selective activity in human VOTC reveals an increase in selectivity/abstraction in face-selective activity from posterior to anterior VOTC, in line with a hierarchical view of visual recognition, yet with response timing indicating that category-selective neural processes occur largely concurrently across the whole VOTC. These findings suggest that visual recognition is supported by category-selective neural populations spread across the VOTC receiving direct concurrent inputs from early visual cortex, without the need to rely on intermediate and successive hierarchical relays.

## MATERIALS AND METHODS

### Participants

The study included 140 participants (71 females, mean age: 33.0±9.2 years; 123 right-handed, 3 ambidextrous) undergoing clinical intracerebral evaluation with depth electrodes (stereotactic electroencephalography or SEEG, **Figure 1A**) for refractory partial epilepsy. Participants were included in the study if they had at least one intracerebral electrode implanted in the ventral occipito-temporal cortex. All participants gave written consent to participate to the study, which was approved by a national human investigation committee certified by the French Ministry of Health (Institutional Review Board: IORG0009855)

#### Fast periodic visual stimulation paradigm

A well validated fast periodic visual stimulation (FPVS) paradigm with natural images was used to elicit face-selective neural activity in iEEG with high signal to-noise ratio (SNR) (see(Rossion et al., 2015) for the original description of the paradigm in EEG; see(Jonas et al., 2016a; Rossion et al., 2018) for its validity in iEEG).

##### Stimuli

Two hundred grayscale natural images of various non-face objects (from 14 non-face categories: cats, dogs, horses, birds, flowers, fruits, vegetables, houseplants, phones, chairs, cameras, dishes, guitars, lamps) and 50 grayscale natural images of faces were used(Jacques et al., 2022; Jonas et al., 2016a; Rossion et al., 2015). Each image contained an unsegmented object or face near the center, these stimuli differing in terms of size, viewpoint, lighting conditions and background. Images were equalized for mean pixel luminance and contrast, but low-level visual cues associated with the faces and visual objects remained highly variable, naturally eliminating the systematic contribution of low-level visual cues to the recorded face-selective neural activity(Gao et al., 2018; Rossion et al., 2015).

##### Procedure

Participants viewed continuous sequences of natural images of objects presented at a fast rate of 6 Hz (i.e., stimulus onset asynchrony of 167ms) through sinusoidal contrast modulation. This relatively fast rate allows only one fixation per stimulus and is largely sufficient to elicit maximal face-selective activity(Retter and Rossion, 2016). Images of faces appear periodically as every 5th stimulus, so that neural activity that is common to faces and nonface stimuli is manifested at 6 Hz and harmonics, while differential (i.e., selective) reliable activity to faces are expressed at 1.2 Hz (i.e., 6 Hz/5) (see **Figure 1B,C**). All images were randomly selected from their respective categories (face and nonface), with the constrain that no image could be immediately repeated. A stimulation sequence lasted 70 s: 66 s of stimulation at full-contrast flanked by 2 s of fade-in and fade-out, where contrast gradually increased or decreased, respectively. During a sequence, participants were instructed to fixate a small black cross which was presented continuously at the center of the stimuli and to detect brief (500 ms) color-changes (black to red) of this fixation-cross. Among the 140 participants, participants viewed either 2 sequences (69 participants), 3 sequences (6 participants), 4 sequences (55 participants), 5 sequences (1 participant), 6 sequences (3 participants), 8 or more sequences (6 participants). No participant had seizures in the 2 hours preceding the recordings.

#### Intracerebral electrode implantation and SEEG recording

Intracerebral electrodes (Dixi Medical, Besançon, France) were stereotactically implanted within the participants’ brains for clinical purposes, i.e., to delineate their seizure onset zones (Talairach and Bancaud, 1973) and to functionally map the surrounding cortex for potential epilepsy surgery. Each 0.8 mm diameter intracerebral electrode contains 5-15 independent recording contacts of 2 mm in length separated by 1.5 mm from edge to edge (**Figure 1A**). A total of 997 electrode arrays were implanted in the VOTC of the 140 participants. These electrodes contained 11121 individual recording contacts in the VOTC (i.e., in the gray/white matter or medial temporal lobe-MTL; 6295 and 4826 contacts in the left and right hemisphere respectively). Intracerebral EEG was sampled at either 500 or 512 Hz and referenced to either a midline prefrontal scalp electrode or an intracerebral contact in the white matter. SEEG signal was re-referenced offline to bipolar reference to limit dependencies between neighboring contacts(Hagen et al., 2025). Specifically, the signal at a given recording contact was computed as the signal measured at that contact (i.e., with the recording reference) minus the signal at the directly adjacent contact located more medially on the same SEEG electrode array. Since SEEG field potentials are computed using pairs of adjacent contacts, each electrode array contains 1 contact less than in the original recording. All subsequent analyses were performed on bipolar-referenced signal in the set of bipolar contacts as described just above.

##### Contact localization in the individual anatomy

The position of each contact relative to brain anatomy was determined in each participant’s own brain by coregistration of the post-operative CT-scan with a T1-weighted MRI of the patient’s head. Anatomical labels of bipolar contacts were determined using the anatomical location of the ‘active’ contact. In cases where the active contact was in the white matter and the ‘reference’ contact was in the gray matter, the active contact was labeled according to the anatomical location of the reference contact. Bipolar contacts in which both the active and reference contacts were in the white matter were excluded from analyses. To accurately assign an anatomical label to each contact, we used the same topographic parcellation of the VOTC as in(Jacques et al., 2022; Jonas et al., 2016b) (**Table S1, Figure S2**).

#### SEEG signal processing and analyses

##### High frequency broadband (HFB) preprocessing

Segments of iEEG corresponding to FPVS stimulation sequences were extracted (74-second segments, -2s to +72s, **Figure 1D**, top) and notched-filtered to remove 50Hz line noise and 2 harmonics (100 and 150Hz). Variation in signal amplitude as a function of time and frequency was estimated by a Morlet wavelet transform applied on each SEEG 74-second segment from frequencies of 30 to 160 Hz, in 2 Hz increments (**Figure 1D**, middle). The number of cycles (i.e., central frequency) of the wavelet was adapted as a function of frequency from 2 cycles at the lowest frequency to 9 cycles at the highest frequency. The temporal smoothing resulting from the wavelet transform was minimal: wavelet of 20 ms of full width at half maximum (FWHM) across the frequency range (i.e. median of FWHM computed at each frequency bin). A simulation of HFB signals with a known onset time and realistic signal-to-noise ratio indicates that the theoretical underestimation of onset latency due to the wavelet analysis is around 5-10 ms, which is on par with the estimation based on the half width at half maximum (= FWHM/2 = 20/2 ms). The wavelet transform was computed on each time-sample and the resulting amplitude envelope was downsampled by a factor of 6 (i.e., to a 85.3 Hz sampling rate) to save disk space and computation time. However, for timing analyses on face-selective contacts, the original recording sampling rate (500 or 512 Hz) was used to preserve temporal resolution. Resulting time-by-frequency SEEG segment were normalized to obtain, for each frequency bin, the percentage of signal change (PSC) generated by the stimulus onset relative to the mean amplitude in a pre-stimulus time-window (-1600 ms to -300 ms relative to the onset of the stimulation sequence). The PSC was then averaged across frequencies (between 30 Hz and 160 Hz) to obtain 74-seconds segments of time-varying HFB amplitude envelope (**Figure 1D**, bottom).

##### HFB frequency-domain processing

For each recording contact, 74s segments (i.e., 2-8 per participant) of HFB PSC envelope were averaged in the time-domain and cropped to contain an integer number of 1.2 Hz cycles beginning 2 seconds after the onset of the FPVS sequence (right at the end of the fade-in period) until approximately 68 seconds, before stimulus fade-out (79 face cycles ≈ 65.8 s). An FFT was then applied to the resulting cropped HFB envelope to obtain the amplitude spectrum in the frequency domain (**Figure 1E**). The frequency-tagging approach used here allows identifying and separating two types of neural activity: (1) a general visual response occurring at the base stimulation frequency (6 Hz) and its harmonics, as well as (2) a face-selective activity at 1.2 Hz and its harmonics(Jacques et al., 2022; Jonas et al., 2016a; Rossion et al., 2018). Face-selective activity significantly above noise level at the face stimulation frequency (1.2 Hz) and its harmonics (2.4, 3.6 Hz, etc.) were determined as follows(Hagen et al., 2025; Jacques et al., 2022): (1) the FFT spectrum was cut into 4 segments centered at the face frequency and harmonics, from the 1st until the 4th (1.2 Hz until 4.8 Hz), and surrounded by 25 neighboring bins on each side; (2) the amplitude values in these 4 segments of FFT spectra were summed; (3) the summed FFT spectrum was transformed into a Z-score. Z-scores were computed as the difference between the amplitude at the face frequency bin and the mean amplitude of surrounding bins divided by the standard deviation of amplitudes in the surrounding bins. A contact was considered as showing a face-selective response in HFB if the Z-score at the frequency bin of face stimulation exceeded 3.1 (i.e., *p* < 0.001 one-tailed: signal>noise).

##### HFB time-domain preprocessing

Recording contacts with a significant face-selective response (based on frequency-domain analyses) were further processed in the time-domain. Starting from the 74-seconds HFB amplitude segments (see ‘HFB preprocessing’), time-series were processed in the following way: (1) an FFT notch filter (filter width = 0.07Hz) was applied to remove the general visual response at 6Hz and 3 additional harmonics (i.e. 6, 12, 18, 24 Hz); (2) time-series were segmented in 1.17 s epochs centered on the onset of each face (i.e. [-2 to + 5] 6Hz-cycles relative to face onset) in the FPVS sequences; (3) resulting epochs were averaged (see **Figure 1F** for an example averaged time-course without the notch filtering of the general visual response). Unless noted otherwise, averaged time-series per contact were baseline-corrected by subtracting the mean amplitude in a [-0.166 to 0 s] time-window relative to face onset. Considering time-series for each contact revealed that face-selective activity manifest either as a periodically *larger* response to faces compared to non-face objects, or as a periodically *smaller* response to faces (**Figure S1**). Response increase/decrease was defined as a function of whether the amplitude in the 0.15s to 0.35s time-window was respectively larger or smaller than the amplitude in the time-window just preceding the onset of the face (1 cycle at 6Hz, i.e. [-0.167s to 0s]). During the FPVS sequences, images are presented using a sinusoidal modulation of contrast, rather than an abrupt onset (**Figure 1A**). Hence, to account for the delay with which the images become visible/perceivable in this setting, the face onset time was shifted forward by 33 ms, as established from a direct comparison of sine wave to square wave/abrupt stimulus presentation in a full EEG study (Retter and Rossion, 2016) and a subset of participants of the present study. This corresponds to 4 frames at 120Hz screen refresh rate and to a face contrast of about 35% of the maximal(Retter and Rossion, 2016).

##### Group visualization and analyses in Talairach space

For group mapping and visualization, anatomical MRIs were spatially normalized to determine the Talairach (TAL) coordinates of VOTC intracerebral contacts. The cortical surface used to display group contact locations and maps was obtained from segmenting the Collin27 brain from AFNI(Cox, 1996), which is aligned to the TAL space. We used TAL transformed coordinates to compute maps of the local proportion of face-selective intracerebral contacts across the VOTC. Local proportion of contacts was computed in volumes (i.e., ‘voxels’) of size 15 x 15 x 100 mm (respectively for the X: left – right, Y: posterior – anterior, and Z: inferior – superior dimensions) by steps of 3 x 3 x 100 mm over the whole VOTC. A large voxel size in the Z dimension enabled collapsing across contacts along the inferior-superior dimension. For each voxel, we extracted the following information across all participants in our sample: (1) number of recorded contacts located within the voxel across all participants; (2) number of significant face-selective contacts. From these values, for each voxel we computed the proportion of face-selective contacts as the number of significant contacts within the voxel divided by the total number of recorded contacts in that voxel. Then, for each voxel we determined whether the proportion of significant contacts was significantly above zero using a percentile bootstrap procedure, as follows: (1) within each voxel, sample as many contacts as the number of recorded contacts, with replacement; (2) for this bootstrap sample, determine the proportion of significant contacts and store this value; (3) repeat steps (1) and (2) 2,000 times to generate a distribution of bootstrap proportions; and (4) estimate the *p*-value as the fraction of bootstrap proportions equal to zero.

##### Quantification of response amplitudes

Amplitude quantification was performed on face-selective contacts. We first computed baseline-subtracted amplitudes in the frequency domain as the difference between the amplitude at each frequency bin and the average of 48 corresponding surrounding bins (up to 25 bins on each side, i.e., 50 bins, excluding the 2 bins directly adjacent to the bin of interest, i.e., 48 bins). Then, for each contact, face-selective amplitude was quantified as the sum of the baseline-subtracted amplitudes at the face frequency from the 1st until the 14th harmonic (1.2 Hz until 16.8 Hz), excluding the 5th and 10th harmonics (6 Hz and 12 Hz) that coincided with the base frequency(Jonas et al., 2016a). General visual response amplitude was quantified separately as the sum of the baseline-subtracted amplitudes at the base frequency from the 1st until the 3rd harmonic (6 Hz until 18 Hz).

##### Face-selectivity index

We computed an index of face-selectivity for face-selective contacts by taking the ratio of the face-selective amplitude (i.e., at 1.2Hz and harmonics, see above) to the sum of the face-selective and general visual amplitude (i.e., 6Hz and harmonics). This index provides an additional quantification of face-selectivity by taking into account the magnitude of response to non-face stimuli. It varies from 0 (no face-selective response) to 1 (face-selective response only, no general visual response). Face-selectivity indices were either computed at the scale of main VOTC regions or computed in Talairach space as a function of the posterior-anterior axis (Y TAL dimension). In the latter case, face-selectivity indices were computed along the Y dimension in segments of 15 mm and by steps of 3 mm, collapsing contacts across the X (lateral-medial, collapsed across both hemispheres) and Z (inferior-superior) dimensions. Face-selective and general visual activity were first averaged across contacts (i.e., within regions or voxels) before computing the index. This avoided obtaining indices outside the [0 - 1] range if, for instance, the denominator is smaller than 1.

##### HFB response timing parameters

In a first group analyses, the timing of face-selective activity was characterized on HFB time-domain responses using 4 parameters: (1) latency of response onset and (2) offset (**Figure S3**), as well as (3) response overlap and (4) response correlation across VOTC regions (**Figure S4**). Onset latency was estimated using 4 different methodologies (**Figure S3**): (a) z-score, (b) delta slope, (c) broken stick, (3) percent peak. In the z-score method (a), the HFB time-series was converted to z-score values by subtracting the mean amplitude in the baseline window of the time-series (i.e. before face onset: [-0.166 to 0s]) and dividing by the standard deviation of the amplitude in the same baseline window. Z-scores were converted to p-values which were FDR-corrected (Benjamini and Hochberg, 1995). Onset latency was the first time-point after 40ms (which was taken as a lower bound for physiologically plausible response latencies after face onset) at which *p* < 0.05 (two-tailed), for at least 30 ms. In the delta slope (b) method, the slopes of the variation in HFB amplitude are computed at each time point within a 25 ms sliding window. Onset latency is defined as the time (with a lower bound of 40 ms) at which the slope exceeds 2.32 times the mean and standard deviation of the slopes in the baseline window (i.e. an ‘acceleration’ of the electrophysiological signal manifesting the onset of the neural response), for a duration of at least 30 ms. In the broken stick method (c), a regression-based method (Mordkoff and Gianaros, 1999), onset latency is defined as the intersection point (lower bound set to 40 ms) of two line segments (with variable slopes) best fitted (least square error fit) to the HFB signal between the start of the baseline window (-166ms) and the end of the ramping-up portion of the signal before the first peak. In the percent peak (d) method, onset latency is defined as the first point, after 40 ms, that rises above 20% of the amplitude difference between the baseline window and the peak (maximum amplitude between 0 and 0.8 s), for at least 30 ms.

Offset latency was defined using the ‘percent peak’ approach, as the point in time, after the onset latency, where the signal reaches below 5% of the amplitude difference between baseline and peak (**Figure S3**).

The first two timing parameters of face-selective response, onset and offset latencies, were quantified per main VOTC region using a hierarchical bootstrapping approach to respect the nested structure of the data (region > participants > contacts > trials). For each bootstrap iteration and each region, we first sampled participants with replacement. Within each sampled participant, we then sampled contacts and then trials within sampled contacts, with replacement. Resampled trials, then contacts within participants, then participants within a region, were successively averaged to obtain a bootstrapped region-level response from which we derived onset latency (4 different methods) and offset latency. We obtained bootstrap distributions of onsets/offsets using 2000 bootstrap iterations per region, allowing to compute the median and 95% confidence interval for these 2 parameters.

As a third timing parameter, we estimated the overlap between the mean time-series for pairs of regions (e.g., region A and B, computed by averaging trials, then contacts within a participant, then the participants, as in the hierarchical bootstrap – see below) within hemispheres (**Figure S4A)**. The overlap is asymmetrical and is calculated separately for region A and region B as the ratio between the area under the curve (AUC) of the overlap between regions A and region B (i.e., summing/integrating the amplitude between the maximum of onset A and B and the minimum of offset A and B) and the total AUC for region A (for overlap of region A to B) or for region B (for overlap of region B to A).

Fourth, for each signal type we computed Pearson correlations between the time-series obtained in the three main VOTC regions, within each hemisphere (**Figure S4B)**. For each pair of regions, we determined whether the between-region correlation coefficient was significantly lower than within-regions correlations using a bootstrapping approach: (1) for each region, compute participants’ level mean time-series by averaging responses across contacts within each participant; (2) compute the *between-region correlation* by repeatedly (5000 times) randomly sampling half of the available participants’ mean time-series from each of the two regions, averaging the time-series for each region, calculate the Pearson correlation between these averaged time-series and take the mean over the obtained 5000 correlation coefficient; (2) compute the within-region correlation for each region in a similar manner by repeatedly (5000 times) randomly splitting the available participants’ time-series in two groups, average the time-series for each group, correlate time-series across the two groups and take the mean over the obtained 5000 correlation coefficient.

#### Statistical analyses: hierarchical permutation tests and equivalence testing

Statistical significance of latency differences between main VOTC regions was assessed using a hierarchical permutation test. We use a stratification approach to partition participants into a paired set (i.e. participants that had recording contacts in the two regions compared) and unpaired set (participants with contacts in a single region). For paired participants, the region labels were randomly swapped within subject (i.e., exchanging the signals from the two regions), thereby preserving all participant-, contact-, and trial-level structure while breaking the association between region and latency estimates. For unpaired participants, participants were randomly reassigned between regions while preserving the original group sizes, to generate pseudo-groups under the null hypothesis of no regional difference. In each permutation, signals were averaged across trials, then contacts, then participants within each permuted group and latency was computed and stored from the resulting region-level signals. We performed 10000 permutations to obtain a distribution of regional differences of latencies under the null hypothesis and determine the p-value as the fraction of the null distribution larger or smaller than the observed (non-permuted) difference.

In addition, to statistically assert whether onset latencies measured across main VOTC regions (OCC, PTL, ATL) were consistent with a concurrent (parallel) face-selective activation, we used Bayesian equivalence testing, relying on two separate metrics: (1) the percentage of differences in region of practical equivalence (ROPE), and (2) the Bayes factor using Cauchy prior. We did not rely on a strict Two one-sided Tests (TOST) procedure as these are notoriously under-powered and require much larger samples to be meaningful (Riesthuis, 2024). Equivalence bounds for ROPE were defined by combining two components: (1) a component of physiological variability and (2) a component reflecting expected delays in response onset between regions attributed to neural conduction delay, given the differential distances separating early visual cortex (EVC) from posterior face-selective regions (e.g. IOG) vs. anterior regions (ATL) and assuming signal mainly travels between VOTC regions through major postero-anterior axis fiber bundles of the Inferior longitudinal fasciculus (ILF) or the inferior fronto-occipital fasciculus (IFOF). Physiological variability corresponded to expected measurement noise and between-subject variability. Physiological equivalence bound was obtained by multiplying a Cohen’s d of 0.2 (conventional threshold for a negligible effect) with the (pooled) between-subjects variability in onset latency (computed for each region using a jackknife procedure and a correction factor of [N-participants – 1] to the jackknife standard deviation). For conduction delay bounds, the expected latency difference between regions under parallel activation depends on: (1) the distance between each region (considering the estimated origin of the signal is the same for all regions - EVC), and (2) neural conduction velocity. For inter-region distance we determined, for each region, the 5 and 95 percentile of the Talairach y-coordinate distribution and defined the maximal distance bounds as the distance between the y-coordinate corresponding to 5% of region 1 (e.g. OCC) to the coordinate corresponding 95% of region 2 (e.g. PTL). This resulted in the following maximum distance values: OCC-PTL: [54]mm; PTL-ATL: [54]mm; OCC-ATL: [83]mm. For conduction velocity, we used a constant value of 3.5 m/s, based on median axonal conduction velocity for cortico-cortical connections (Lemarechal et al., 2022; van Blooijs et al., 2023), which was more conservative than using a range of values (e.g. 1.7 to 5.3 m/s based on (Lemarechal et al., 2022)). Maximum expected conduction delay was computed as [maximum distance / conduction speed] (e.g. for OCC to PTL: 0.054 / 3.5 = 15 ms). Resulting equivalence bounds (ROPE) were asymmetrical given than one region (e.g. OCC) is always closer to the source (EVC) than the other region (e.g. PTL) and were defined as [-1*physiologial_bound +1*physiologial_bound+max_conduction_delay]. For instance, using the z-score method to measured onset latencies, ROPE was [-12 to 27] ms for OCC to PTL, meaning that under equivalence, OCC can be activated up to 27 ms earlier than PTL (maximum conduction delay + noise), while allowing for some instances where OCC activates later (up to 12ms, due to noise only).

For each pair of region compared, we used a bootstrap procedure to (1) define the percentage of differences between regions that fall within the ROPE, (2) compute the bayes factor using a Cauchy distribution (scale = 0.5) to estimate the proportion of the prior distribution in ROPE, and the bootstrap distribution to estimate the proportion of the posterior distribution in ROPE. The bootstrap distribution was obtained using a hierarchical stratified bootstrap procedure that naturally respects the nested structure of the data, that accommodates for unequal numbers of participants, contacts, and trials across regions, as well as partially overlapping participants samples across regions.

For each bootstrap iteration, with first sample participants with replacement within each stratum (i.e. paired vs unpaired participants samples). For paired participants, sampling was performed jointly across regions to preserve the dependency structure, whereas unpaired participants were sampled independently within each region. Within each sampled participant, contacts were then resampled with replacement, and within each contact, trials were resampled with replacement. For paired participants, trial resampling was performed using identical trials across sampled contacts with a participant to preserve trial-level covariance. Resampled trials were averaged at the contact level, contact-level signals were averaged within participant, and participant-level signals were averaged to obtain a region-level response. Onset latency was then estimated from this averaged signal for each region using one of the 4 methods defined above (‘HFB response timing parameters). This procedure was repeated across 2000 bootstrap iterations to obtain a distribution of latency estimates for each region that respects the structure of data at iteration-level. Latency differences between regions were computed at each iteration, yielding a bootstrap distribution of differences which was used to compute percentage of differences in ROPE and posterior distribution for the Bayes factor.

##### Functional connectivity between VOTC regions

Connectivity between regions was computed using participants that had face-selective contact(s) in at least 2 of the dominant face-selective regions in the same hemisphere: right/left IOG, latFG, antFG+. We used largely the same approach as in (Kadipasaoglu et al., 2017), except for the statistics. Starting from the single-trial HFB time-domain activity per recording contact, we performed the following steps: (1) dowsampling of the time-series by a factor of 2 to save computation time; (2) baseline-correction of each trial by subtracting the mean amplitude in the -166 to 0 ms; (3) subtract from each single-trial, the mean across trials to obtain trial-by-trial variability around the mean. The single-trial amplitudes were correlated between pairs of electrodes at each time point from -100 to 550 ms to compute the instantaneous functional connectivity between regions. To explore the directionality in the connectivity, we also computed the across-trial correlations after introducing variable temporal lags between regions (from -150 to 150 ms). For each pair of contacts, we lagged the time-series at one of the contact before computing the correlation. This allows determining directionality between regions if the amplitude of face-selective activity at a given time point in one region correlates with amplitude in the other region at an earlier or later time-point. Correlations were summarized in lagged-correlation matrices (**Figure 5**). In each participant and each pair of regions, correlations were computed across all possible pairs of face-selective contacts, averaged across contacts and then averaged across participants to obtain group estimates. Statistics were performed using a randomization test to determine the null distribution, by running 1000 times the same analyses steps after having randomized the order of trials between pairs of electrodes being correlated. Group-level correlations were tested against the null distributions at p < 0.01 fdr-corrected.

##### HFB onset latency map

In addition to computing timing parameters across main VOTC regions, we computed a group map of local face-selective response onset latency across VOTC in Talairach space, collapsed across hemisphere. We collected HFB time-series from contiguous face-selective contacts located within voxels of 20 x 20 x 100 mm (swept across VOTC in steps of 3 x 3 x 100 mm; see above ‘*Group visualization and analyses in Talairach space’)*, averaged collected time-series across contacts within-participant and then across participants, and computed onset latency of face-selective response within the current voxels using the 4 methods described above (see ‘*HFB response timing parameters’,* **Figure S3**). To prevent noisy measurements, we only considered voxels with at least 10 contacts. In a similar analysis, we also examined the postero-anterior variation of face-selective response latency by averaging HFB time-series from contacts located in voxels of 20 mm (along the Y dimension) collapsing contacts across the X (lateral-medial) and Z (inferior-superior) dimensions. Voxels were swept across the postero-anterior axis in steps of 3 mm. We used a randomization procedure to determine whether the measured postero-anterior profile of face-selective response onset latency deviate from what would be expected under the (null) hypothesis that the postero-anterior VOTC location has no influence on the onset latency, accounting for potential effect of conduction delay. Specifically, we built a random null distribution by repeatedly (5000 times) computed the postero-anterior profile of onset latency after having randomly shuffled the association between contacts (and associated averaged time-series) and their TAL coordinates. The 99% confidence interval was built by summing the low and high bounds of the interval obtained from the random coordinate shuffling respectively with the expected fast (5.3 m/s) and slow (1.7m/s) conduction delays starting at 0 ms at the most posterior point of our antero-posterior onset latency profile (**Figure 6B**). P-values were determined for each voxel as the fraction of the obtained random distribution that was higher than the actual measured onset latencies and were corrected for multiple comparisons using FDR(Benjamini and Hochberg, 1995).

## Acknowledgments/Funding

This research was supported by ERC AdG HUMANFACE 101055175

## Supporting Information for

**Supplementary Table 1:**
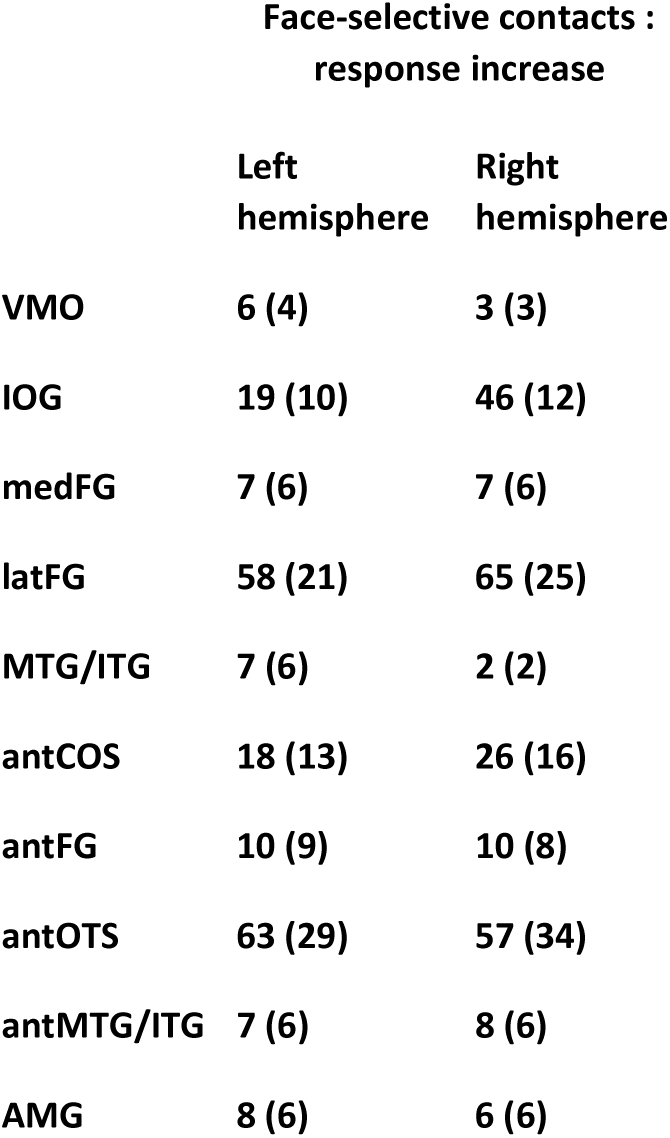
Number of contacts showing significant face-selective activity increase in each anatomical region. The corresponding number of participants in which these contacts were found is indicated in parenthesis. Acronyms: VMO: ventro-medial occipital cortex; IOG: inferior occipital gyrus; medFG: medial fusiform gyrus and collateral sulcus; latFG: lateral fusiform gyrus and occipito-temporal sulcus; MTG/ITG: inferior and middle temporal gyri; antCoS: anterior collateral sulcus; antOTS: anterior occipito-temporal sulcus; antFG: anterior fusiform gyrus; antMTG/ITG: anterior middle and inferior temporal gyri.

**Supplementary Figure 1:**
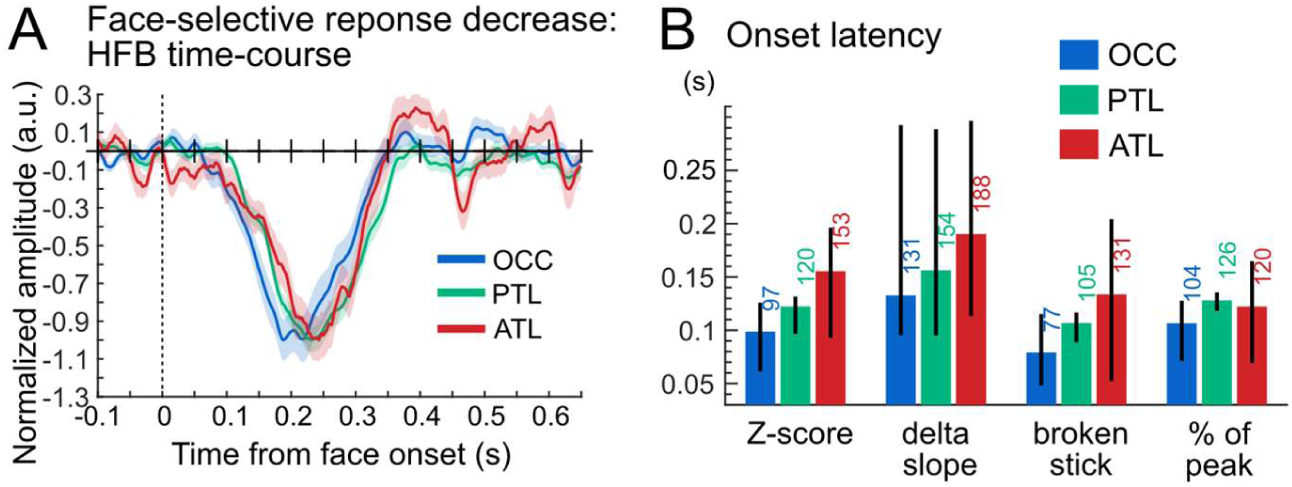
Response timing in contacts showing face-selective response decrease. **A.** Time-domain face-selective HFB activity averaged by main VOTC region (OCC, PTL, ATL). HFB time-series were notch-filtered to remove the general visual response at 6 Hz and harmonics, leaving only face-selective signals. The maximum amplitude of each averaged waveform was normalized to -1 for visualization purposes only. Shaded area represents the standard error of the mean between participants. **B**. Onset latency for each VOTC main region and for 4 latency estimation methods, together with 95% confidence interval (percentile bootstrap).

**Supplementary Figure 2:**
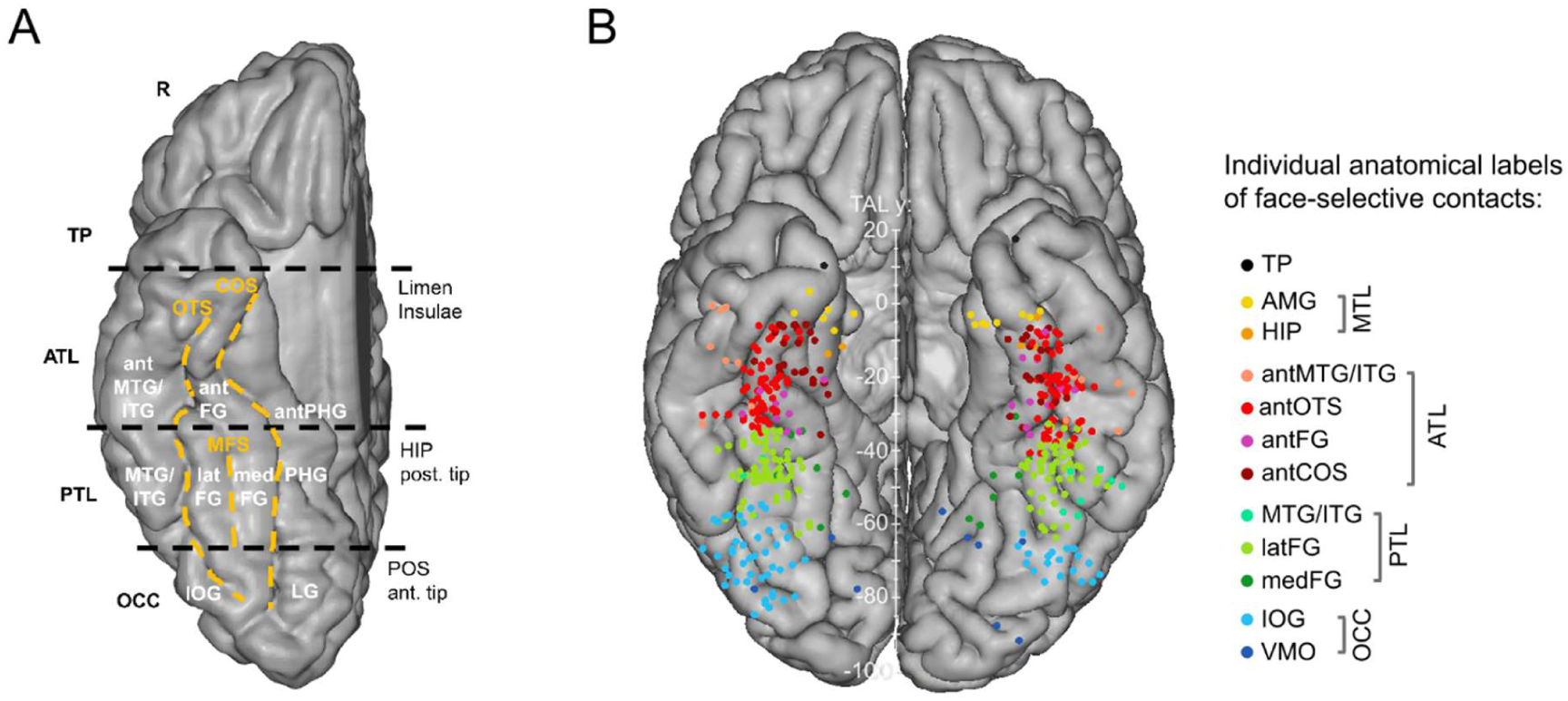
VOTC anatomical parcellation scheme and face-selective recording contacts anatomical labels. **A**. Anatomical regions were defined in each individual hemisphere according to major anatomical landmarks. The ventral temporal sulci (COS, OTS, and midfusiform sulcus, i.e., MFS) serve as medial/lateral borders of regions, whereas three coronal reference planes containing anatomical landmarks (OCC-PTL: anterior tip of the parieto-occipital sulcus, i.e., POS, PTL-ATL: posterior tip of the hippocampus, i.e., HIP, ATL-TP: limen insulae) serve as an anterior/posterior boundary for each region. Electrode contacts were considered to be in the ATL if they were located anteriorly to the posterior tip of the hippocampus and posteriorly to the limen insulae. The schematic locations of these anatomical structures are shown on a reconstructed cortical surface of the Colin27 brain. Acronyms: TP: temporal pole; ATL: anterior temporal lobe; PTL: posterior temporal lobe; OCC: occipital lobe; PHG: parahippocampal gyrus; CoS: collateral sulcus; FG: fusiform gyrus; ITG: inferior temporal gyrus; MTG: middle temporal gyrus; OTS: occipito-temporal sulcus; CS: calcarine sulcus; IOG: inferior occipital gyrus; LG: lingual gyrus; ant: anterior; lat: lateral; med: medial. **B.** Map of all face-selective recording contacts and displayed in the Talairach space. Each circle represents a single face-selective contact color-coded according to its anatomical location in the original individual anatomy (see legend on the right).

**Supplementary Figure 3:**
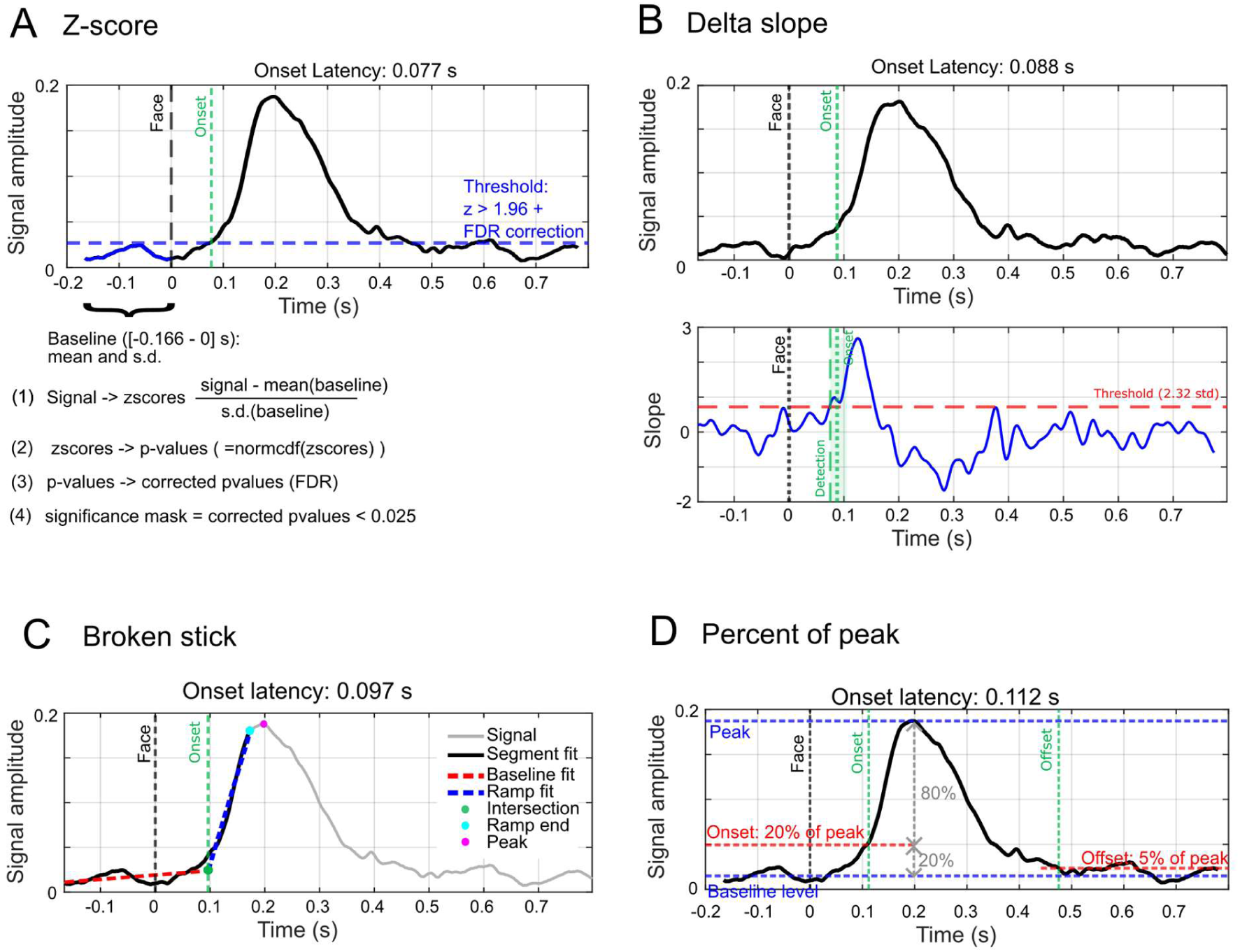
Estimating Onset and offset latencies. The onset latencies of face-selective activity were characterized using 4 different methods. The offset latency was characterized with one methods (i.e., percent of peak). **A.** In the *z-score* method, the HFB time-series was converted to z-score values by subtracting the mean amplitude in the baseline window of the time-series (i.e. before face onset: [-0.166 to 0s]) and dividing by the standard deviation of the amplitude in the same baseline window. Z-scores were converted to p-values which were FDR-corrected (Benjamini and Hochberg, 1995). Onset latency was the first time-point after 40ms (which was taken as a lower bound for physiologically plausible response latencies after face onset) at which *p* < 0.05 (two-tailed), for at least 30 ms. **B.** In the *delta slope* method, the slopes of the variation in HFB amplitude are computed at each time point within a 25 ms sliding window (see bottom plot). Onset latency is defined as the time (with a lower bound of 40 ms) at which the slope exceeds 2.32 times the mean and standard deviation of the slopes in the baseline window (i.e. an ‘acceleration’ of the electrophysiological signal manifesting the onset of the neural response), for a duration of at least 30 ms. **C.** In the *broken stick* method, a regression-based method (Mordkoff and Gianaros, 1999), onset latency is defined as the intersection point (lower bound set to 40 ms) of two line segments (with variable slopes) best fitted (least square error fit) to the HFB signal between the start of the baseline window (-166ms) and the end of the ramping-up portion of the signal before the first peak. **D.** In the *percent peak* method, onset latency is defined as the first point, after 40 ms, that rises above 20% of the amplitude difference between the baseline window and the peak (maximum amplitude between 0 and 0.8 s), for at least 30 ms. Offset latency was defined using the ‘percent peak’ approach, as the point in time, after the onset latency, where the signal reaches below 5% of the amplitude difference between baseline and peak.

**Supplementary Figure 4:**
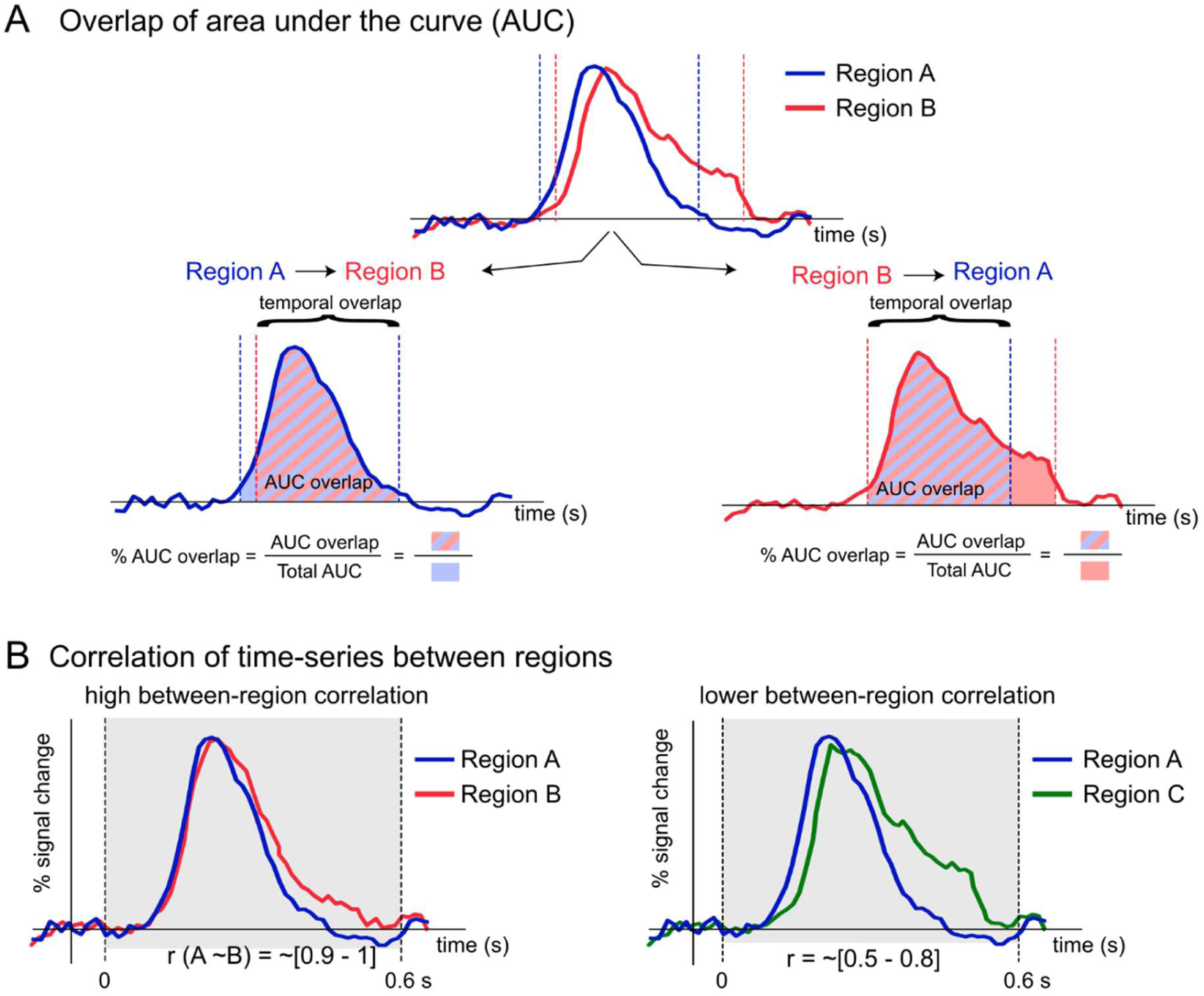
**Additional parameters for timing analyses.** The timing of face-selective activity was also characterized by computing response overlap and response correlation across VOTC regions. **A.** A parameter estimated the overlap between time-series for pairs of regions (e.g., regions A and B). The overlap is asymmetrical and is calculated separately for region A and region B as the ratio between the area under the curve (AUC) of the overlap between regions A and region B (i.e., summing the amplitude between the maximum of onset A and B and the minimum of offset A and B) and the total AUC for region A (for overlap of region A to B) or for region B (for overlap of region B to A). **B.** For the last timing parameter, Pearson correlations were computed between the time-series of pairs of regions using data between 0 and 0.6 s relative to face onset (gray shaded area). This provided an estimate of the similarity between the response function of the two regions compared. The shape of the response is more similar between region A and B than between region A and C. Between region correlations were compared against within-region correlations.

**Supplementary Figure 5:**
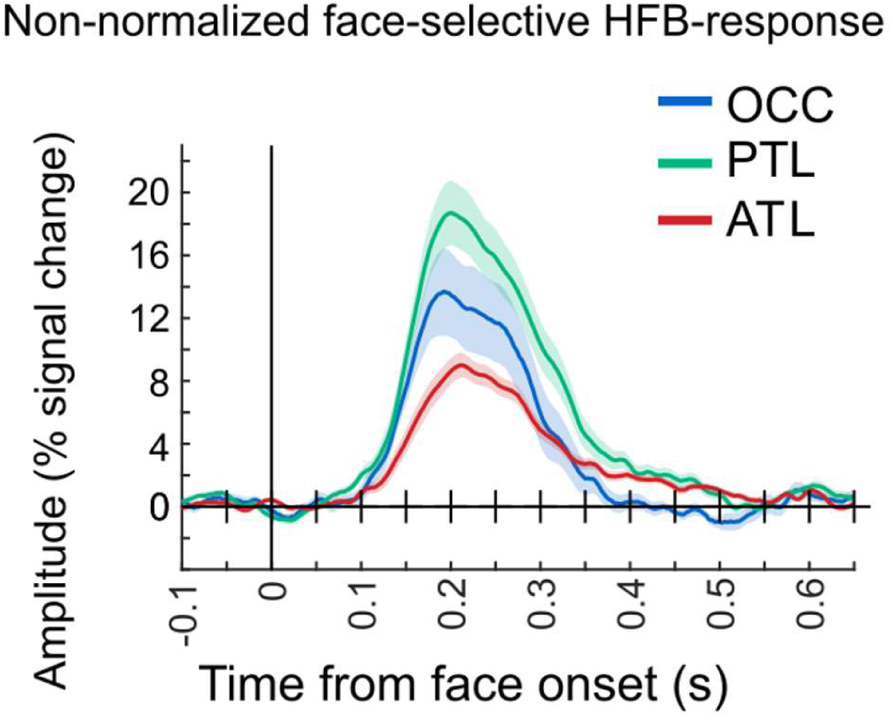
non-normalized time-courses. Time-domain face-selective HFB activity in contacts showing response increase, averaged by main VOTC region (OCC, PTL, ATL) across hemisphere. HFB time-series were notch-filtered to remove the general visual response at 6 Hz and harmonics, leaving only face-selective signals. Time-series are original non-normalized versions of Figure 4 of the main text. Shaded area represents the standard error of the mean between participants.

**Supplementary Figure 6:**
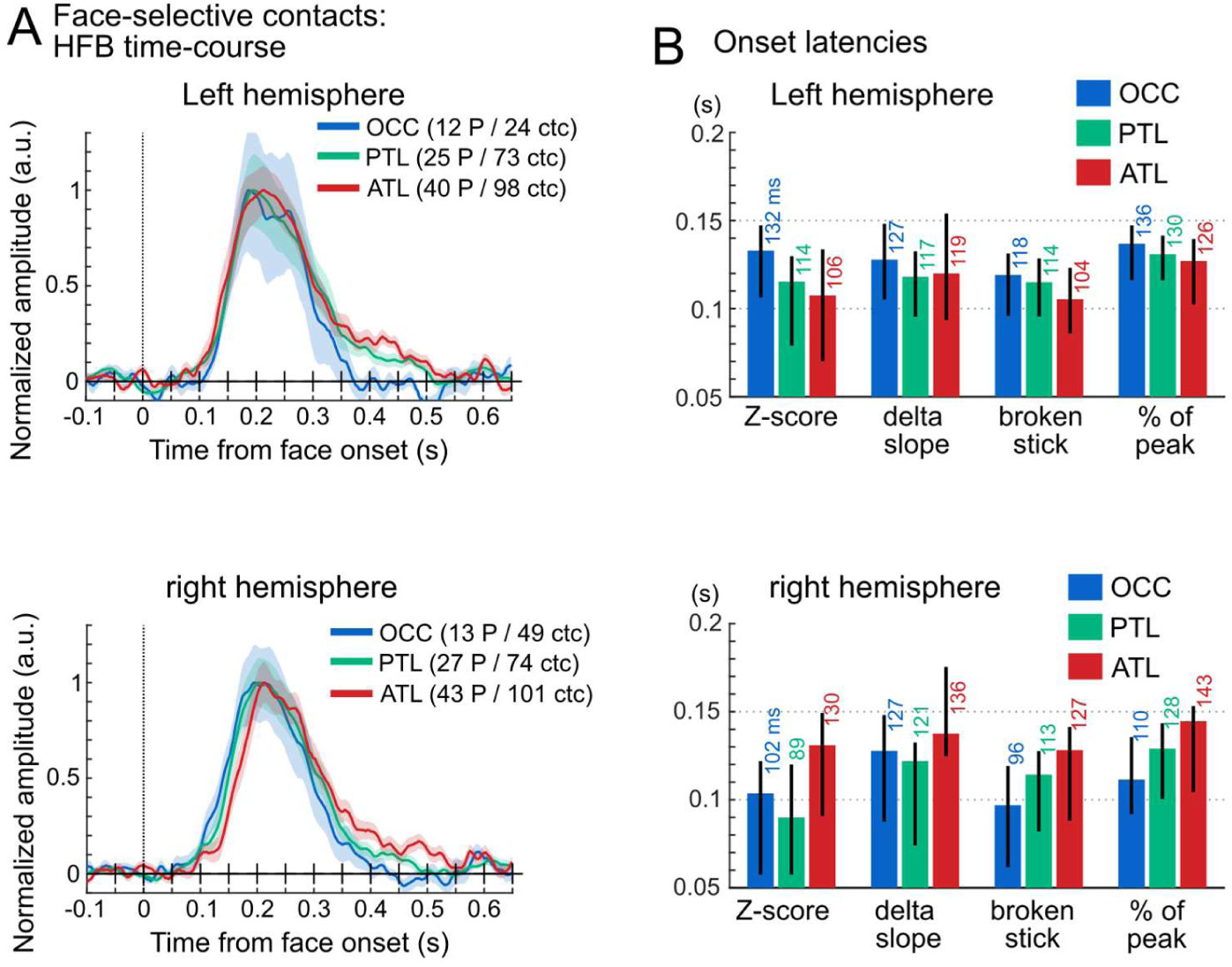
Time-course face-selective contacts split by hemispheres. Response timing in face-selective contacts grouped by main region separately for left (top) and right (bottom) hemispheres. **A.** Mean time-domain face-selective HFB activity. HFB time-series were notch-filtered to remove the general visual activity at 6 Hz and harmonics, leaving face-selective signals only. The maximum amplitude of each averaged waveform was normalized to 1 for visualization purposes only. Shaded area represents the standard error of the mean between participants. Number of participants and contacts are indicated in the legend. **B**. Onset latency for each VOTC main region and for 4 latency estimation methods, together with 95% confidence interval (percentile bootstrap).

**Supplementary Figure 7:**
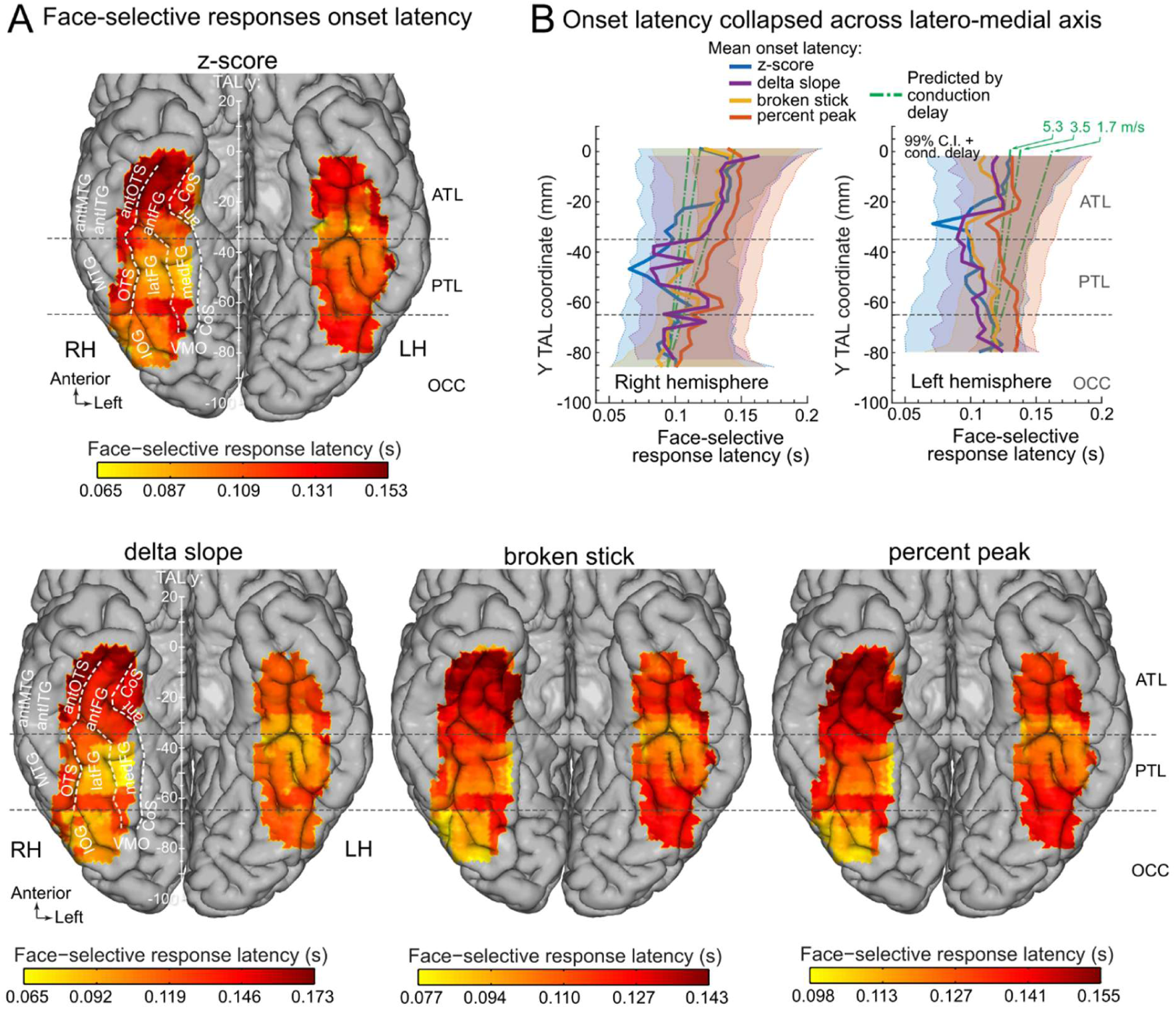
Mapping response onset latency of face-selective contacts split by hemispheres. A. Face-selective onset latency map across VOTC, displayed for 4 different methods to estimate response onset latency. We collected HFB time-series from contiguous face-selective contacts located within voxels of 20 x 20 x 100 mm (swept across VOTC in steps of 3 x 3 x 100 mm), averaged collected time-series across contacts within-participant and then across participants, and computed onset latency of face-selective response within the current voxels. B. Variation of face-selective response latency along the postero-anterior axis for 4 estimation methods and split by hemisphere. For each method, each data point represents the onset latency measured from the time-series averaged over contacts collapsed across the medio-lateral X dimension within 20 mm segments (in the Y dimension). Thick lines are estimated onset latencies and shaded areas show the 99% confidence intervals expected under the null hypothesis that the postero-anterior location has no influence on the onset latency, accounting for the expected conduction delays between posterior and anterior VOTC. Green lines show the expected increase in response onset latency based on simple axonal conduction delays(Lemarechal et al., 2022; van Blooijs et al., 2023) due to increasing distance from the occipital region, with reference to the latency averaged over the OCC region for the ‘z-score’ method. Lines show mean expected conduction velocity for direct cortico-cortical connections (∼3.5 m/s) and +/- 1 std (1.7 and 5.3 m/s).

**Supplementary Table 2 :**
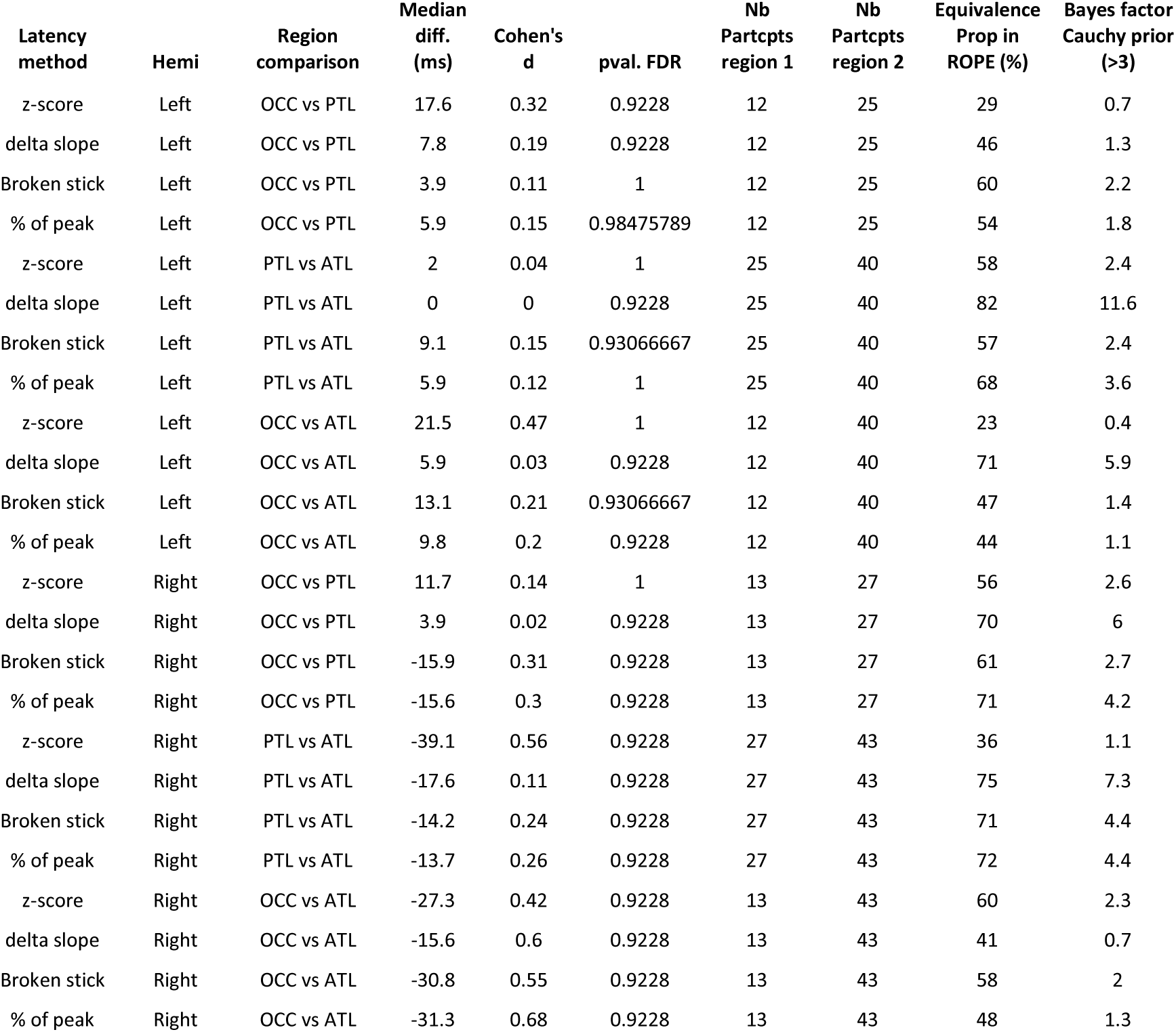
Statistical comparisons of face-selective onset latencies across regions (OCC vs PTL, PTL vs ATL, OCC vs ATL) in each hemisphere using permutation tests et equivalence testing (proportion of difference in ROPE and Bayes factor) for 4 different onset estimation methods. Table contains the median latency difference between regions (Median diff.), effect size (Cohen’s d) of the difference, p-value of the permutation test (pval. FDR), number of participants included in each region.

**Supplementary Figure 8:**
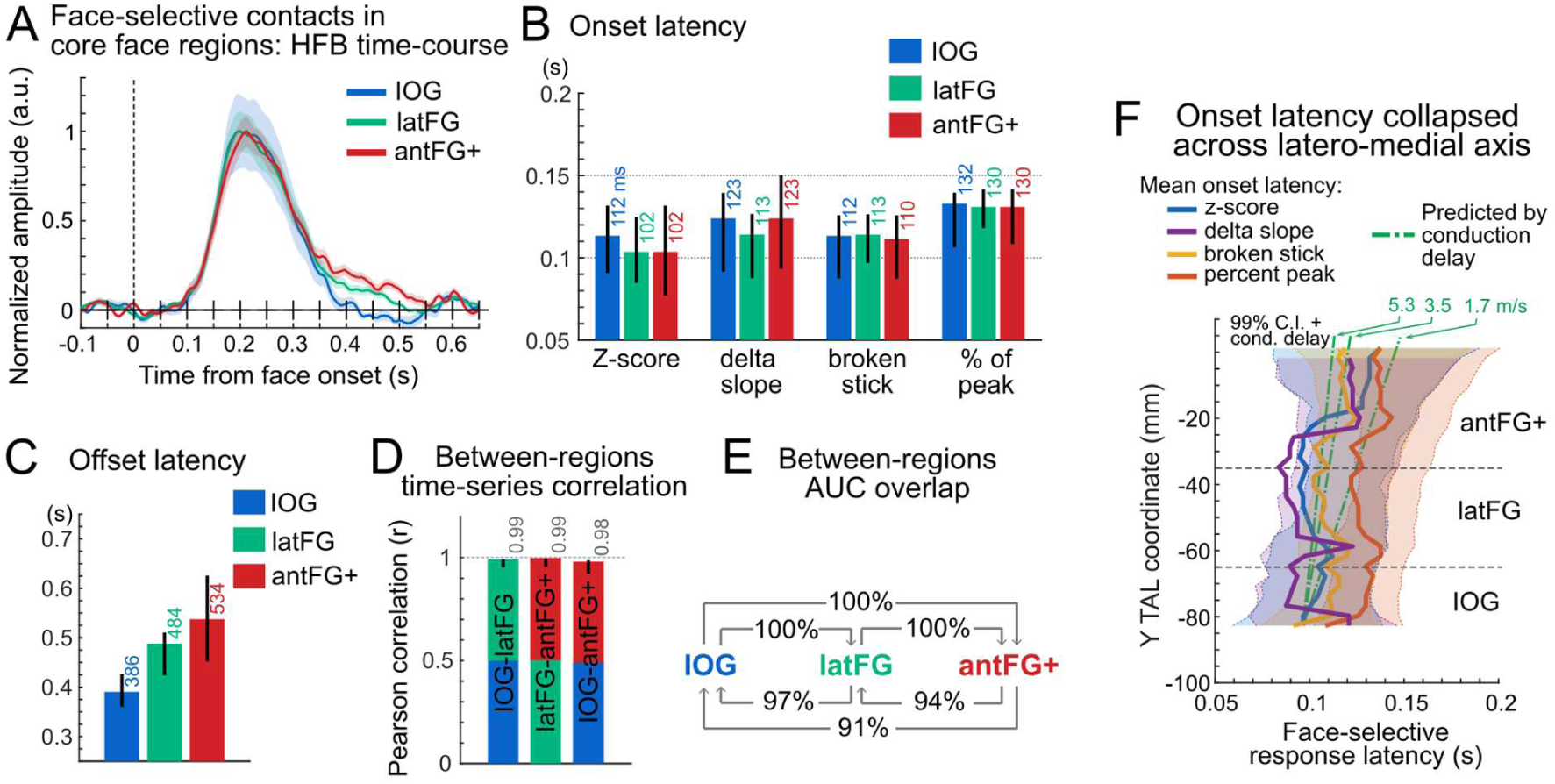
**Response timing in face-selective contacts in core VOTC face-selective regions.** Response timing in face-selective contacts showing response increase in core VOTC face-selective regions: Inferior Occipital Gyrus (IOG, (N=65), lateral Fusiform Gyrus and OTS (latFG, N=123) and antFG+ (anterior fusiform Gyrus, Anterior Occipitotemporal Sulcus, Anterior Collateral Sulcus, N=184). **A.** Mean time-domain face-selective HFB activity in each core face-selective VOTC region collapsed across hemisphere. HFB time-series were filtered to remove the general visual response at 6 Hz and harmonics so only face-selective signal remains. The maximum amplitude of each averaged waveform was normalized to 1 for visualization purposes only. Shaded area represents the standard error of the mean between participants. **B.** Onset latency for each VOTC main region and for 4 latency estimation methods, together with 95% confidence interval (percentile bootstrap). **C.** Offset latency in each region. **D.** Correlation of time-series between regions and 95% confidence intervals. **E.** Between-region area under the curve (AUC) overlap. The arrow shows the directionality of the computed overlap. For instance, the arrow from ATL to OCC indicates the percentage of the total AUC of ATL (measured between onset and offset latencies) occupied by the AUC of its temporal overlap with OCC (determined between max(onset(OCC,ATL)) and min(offset(OCC,ATL))). **F.** Variation of face-selective response latency along the postero-anterior axis for 4 estimation methods. For each method, each data point represents the onset latency measured from the time-series averaged over contacts collapsed across the medio-lateral X dimension within 20 mm segments (in the Y dimension). Thick lines are estimated onset latencies and shaded areas show the 99% confidence intervals expected under the null hypothesis that the postero-anterior location has no influence on the onset latency, accounting for the expected conduction delays between posterior and anterior VOTC. Green lines show the expected increase in response onset latency based on simple axonal conduction delays (Lemarechal et al., 2022; van Blooijs et al., 2023)due to increasing distance from the occipital region, with reference to the latency averaged over the OCC region for the ‘z-score’ method. Lines show mean expected conduction velocity for direct cortico-cortical connections (∼3.5 m/s) and +/- 1 std (1.7 and 5.3 m/s).

**Supplementary Table 3 :**
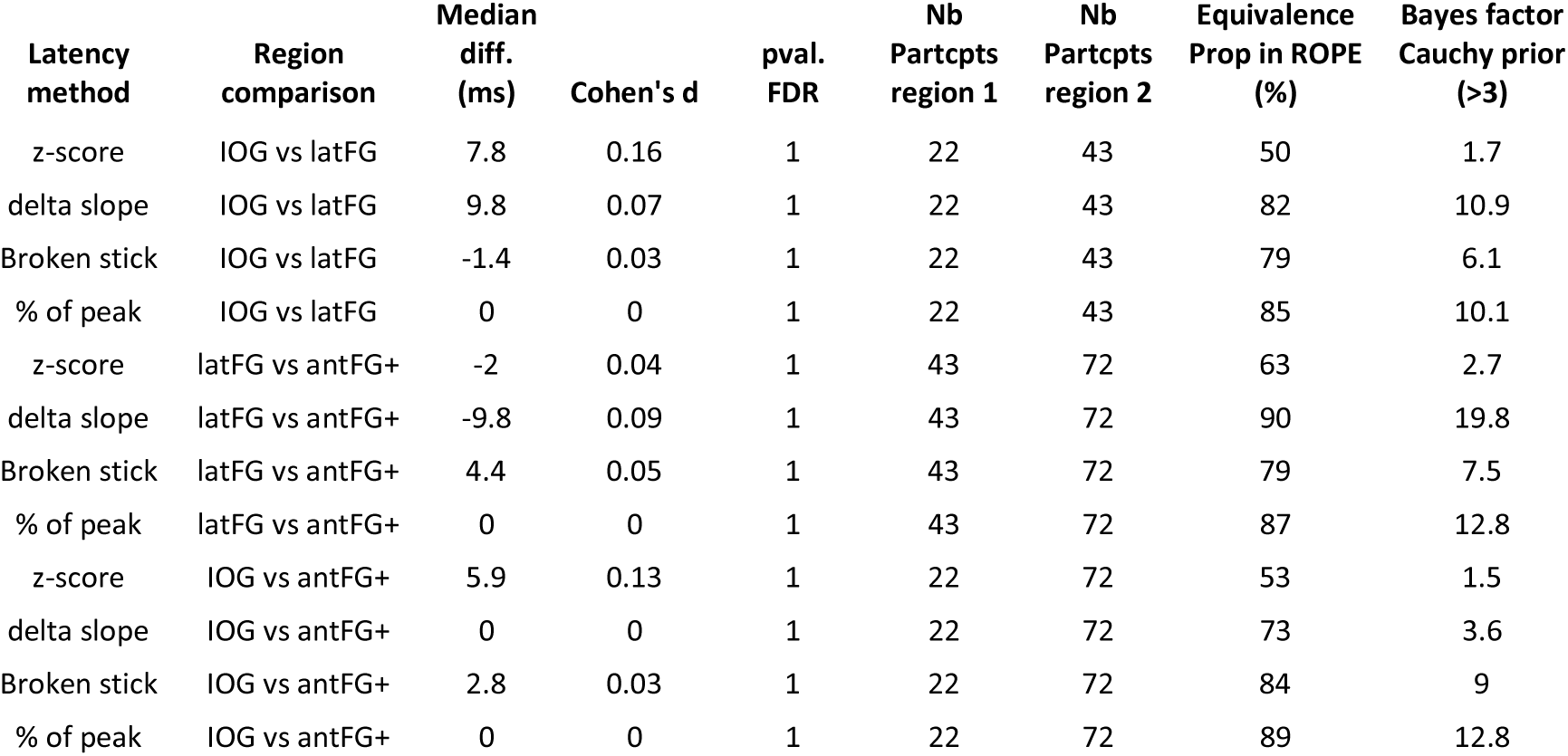
Statistical comparisons of face-selective onset latencies measured in **HFB signal** across **core face-selective regions** (IOG, latFG, antFG) using permutation tests et equivalence testing (proportion of difference in ROPE and Bayes factor) for 4 different onset estimation methods (same as in main). Table contains the median latency difference between regions (Median diff.), effect size (Cohen’s d) of the difference, p-value of the permutation test (pval. FDR), number of participants included in each region.

**Supplementary Figure 9:**
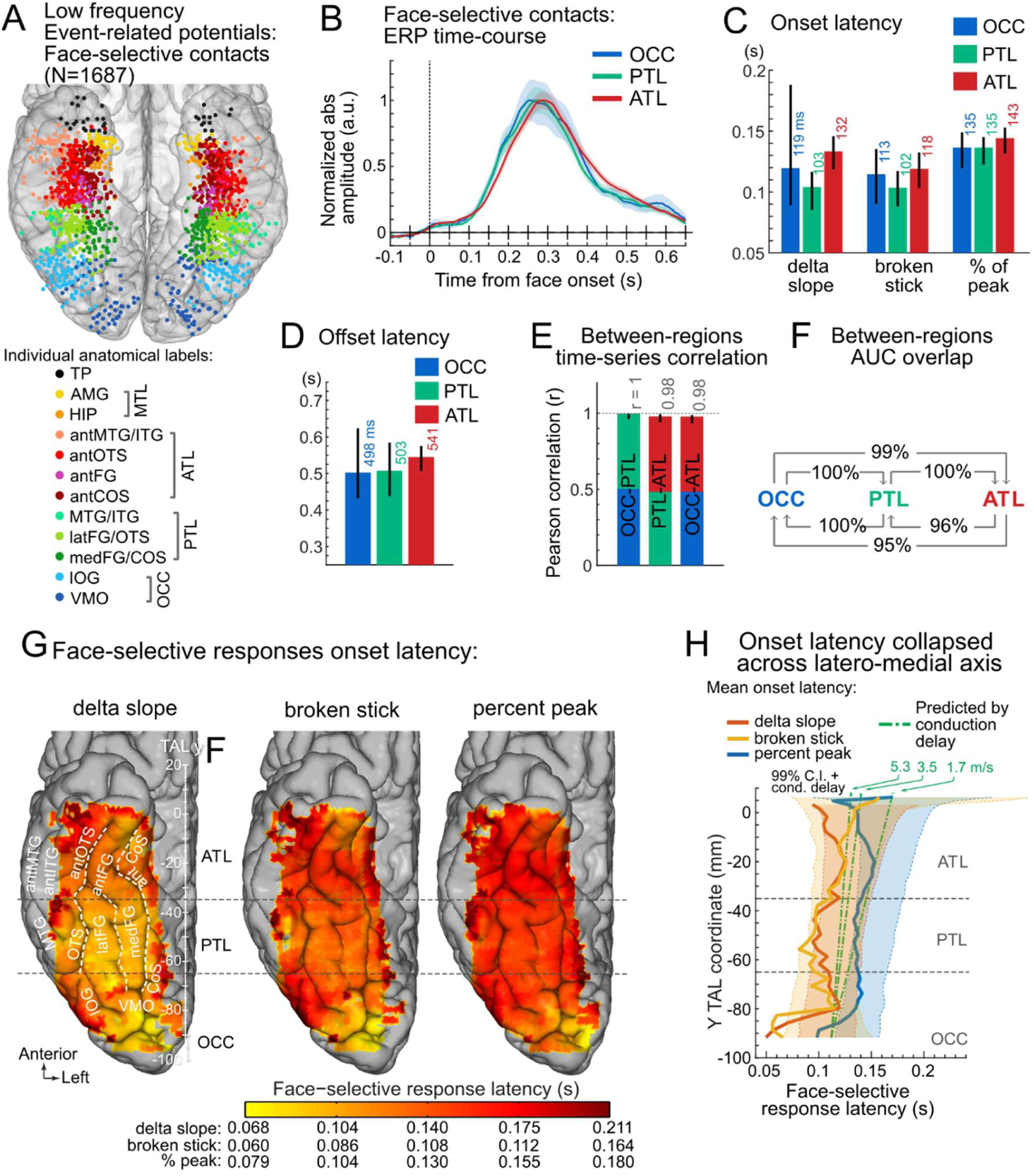
Face-selective activity, low frequency event-related potentials (ERPs). **A.** Map of all VOTC contacts showing a significant (Z>3.1) face-selective response in low-frequency ERPs, displayed in the Talairach space in a transparent reconstructed cortical surface of the Colin27 brain (ventral view). Each circle represents a single recording contact. Contacts are color-coded according to their anatomical location in the original individual anatomy. Low frequency signal is the electrophysiological response measured at each recording contact after bipolar re-referencing (Jacques et al., 2022), dominated by low frequency event-related voltage variations. Only contacts with split-half correlation in the time-domain (i.e. reliability) > 0.66 (i.e. 75% of contacts) were kept to unsure reliable latency estimations. **B.** Grand average face-selective ERP in each face-selective VOTC region collapsed across hemispheres. ERP time-series were filtered to remove the general visual response at 6 Hz and harmonics, leaving face-selective signals only. Averaged ERP were baseline-corrected relative to a [-0.166 0]s window before face onset and the absolute value was computed to allow averaging across contacts/participants despite variability in ERP polarity and morphology. ERP at individual recording contacts were averaged by participants and then across participants in each region. The maximum amplitude of each averaged waveform was normalized to 1 for visualization purposes only. Shaded area represents the standard error of the mean between participants. **C**. Onset latency for each VOTC main region and for 3 latency estimation methods (same as in main text except for ‘z-score’ method, which is unusable with low frequencies), together with 95% confidence interval (percentile bootstrap). **D**. Offset latency in each region. **E**. Correlation of time-series between regions and 95% confidence intervals. **F**. Between-region area under the curve (AUC) overlap. The arrow shows the directionality of the computed overlap. For instance, the arrow from ATL to OCC indicates the percentage of the total AUC of ATL (measured between onset and offset latencies) occupied by the AUC of its temporal overlap with OCC (determined between max(onset(OCC,ATL)) and min(offset(OCC,ATL))). **G.** Face-selective onset latency map across VOTC, collapsed across hemispheres and displayed for 3 different methods to estimate response onset latency. **H.** Variation of face-selective response latency along the postero-anterior axis for 3 estimation methods. For each method, each data point represents the onset latency measured from the time-series averaged over contacts collapsed across the medio-lateral X dimension within 20 mm segments (in the Y dimension). Thick lines are estimated onset latencies and shaded areas show the 99% confidence intervals expected under the null hypothesis that the postero-anterior location has no influence on the onset latency, accounting for the expected conduction delays between posterior and anterior VOTC. Green lines show the expected increase in response onset latency based on axonal conduction delays(Lemarechal et al., 2022; van Blooijs et al., 2023) due to increasing distance from the occipital region, with reference to the latency averaged over the OCC region for the ‘z-score’ method. Lines show mean expected conduction velocity for direct cortico-cortical connections (∼3.5 m/s) and +/- 1 std (1.7 and 5.3 m/s). Note that response latency from few recording contacts located posterior to y = -85 (around and posterior/medial to the posterior transverse Collateral Sulcus), likely in early visual cortex (V1,V2v,V3v,hV4, (Brewer et al., 2005; Winawer and Witthoft, 2015)) and potentially driven by low-level cues (Or et al., 2019), is up to 40 ms earlier than responses measured elsewhere in the VOTC, where little difference is observed.

**Supplementary Table 4 :**
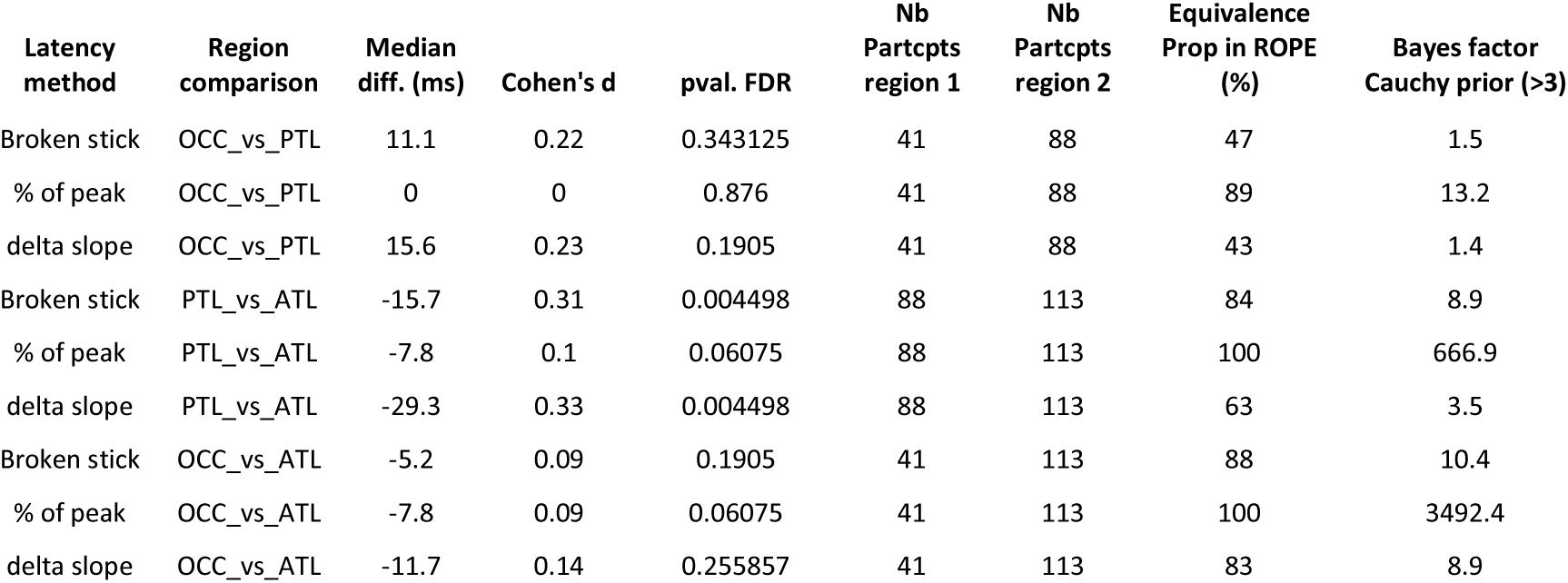
Statistical comparisons of face-selective onset latencies measured in **low frequency event-related potentials** across regions (OCC vs PTL, PTL vs ATL, OCC vs ATL) using permutation tests et equivalence testing (proportion of difference in ROPE and Bayes factor) for 3 different onset estimation methods (same as in main text except for ‘z-score’ method, which is unusable with low frequencies). Table contains the median latency difference between regions (Median diff.), effect size (Cohen’s d) of the difference, p-value of the permutation test (pval. FDR), number of participants included in each region. Despite significantly later ATL than PTL onset latencies (with small effect sizes), the difference is still within bounds of equivalence when taking into account estimated axonal conduction delays between the two regions.

**Supplementary Figure 10:**
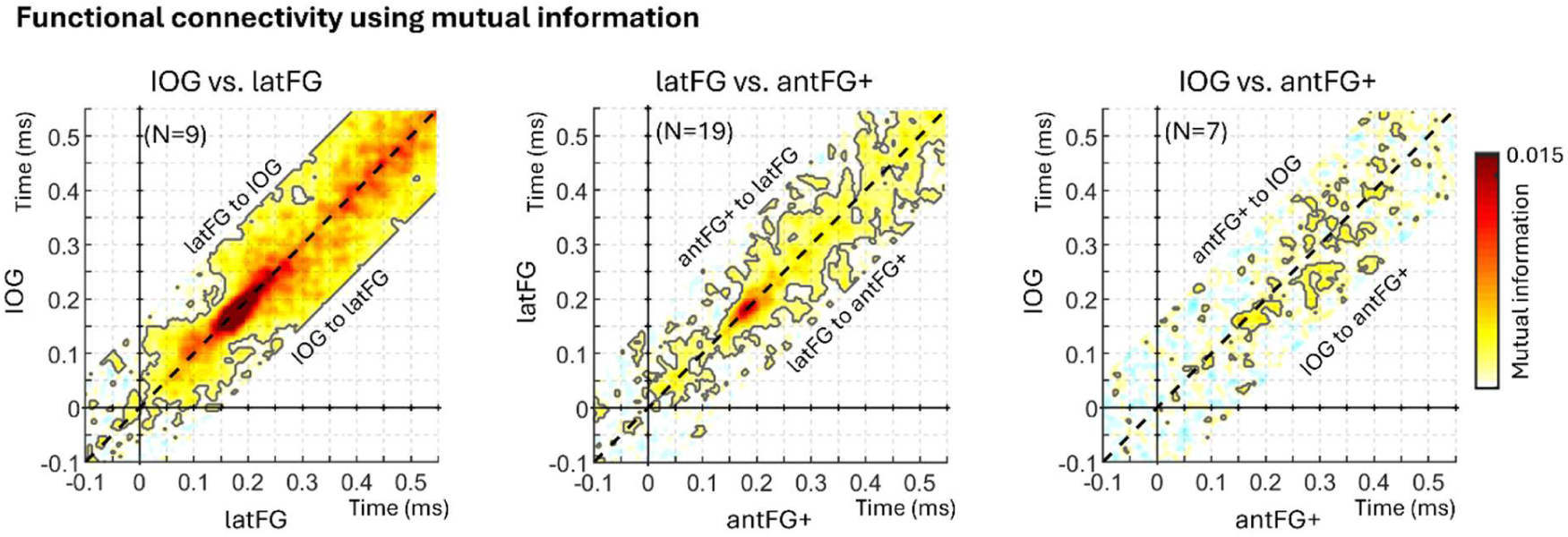
Concurrent functional connectivity between face-selective regions computed using mutual information. Each plot shows the group-averaged mutual information (MI) between single-trials face-selective amplitude measured at two distinct face-selective region (left: IOG and latFG; middle: latFG and antFG+; right: IOG and antFG+) computed at each time point, representing functional connectivity between pairs of regions. MI was computed using the method and Matlab code described in (Ince et al., 2017). Statistics were performed using permutation tests as for Pearson correlations presented in the main manuscript. Black contour lines indicate significant MI (*p* < 0.01, fdr-corrected). MI was computed across -150 to 150 ms lags between regions to infer direction of connectivity. The black dashed diagonal line represents a 0 ms time-lag between regions. MI centered above the diagonal would indicate that face-selective activity in the more anterior region (e.g. latFG in left panel) is coupled with but precedes activity in the posterior region (e.g., IOG in left panel), suggesting an information flow from anterior to posterior, and the reverse for MI centered below the diagonal.

**Supplementary Figure 11:**
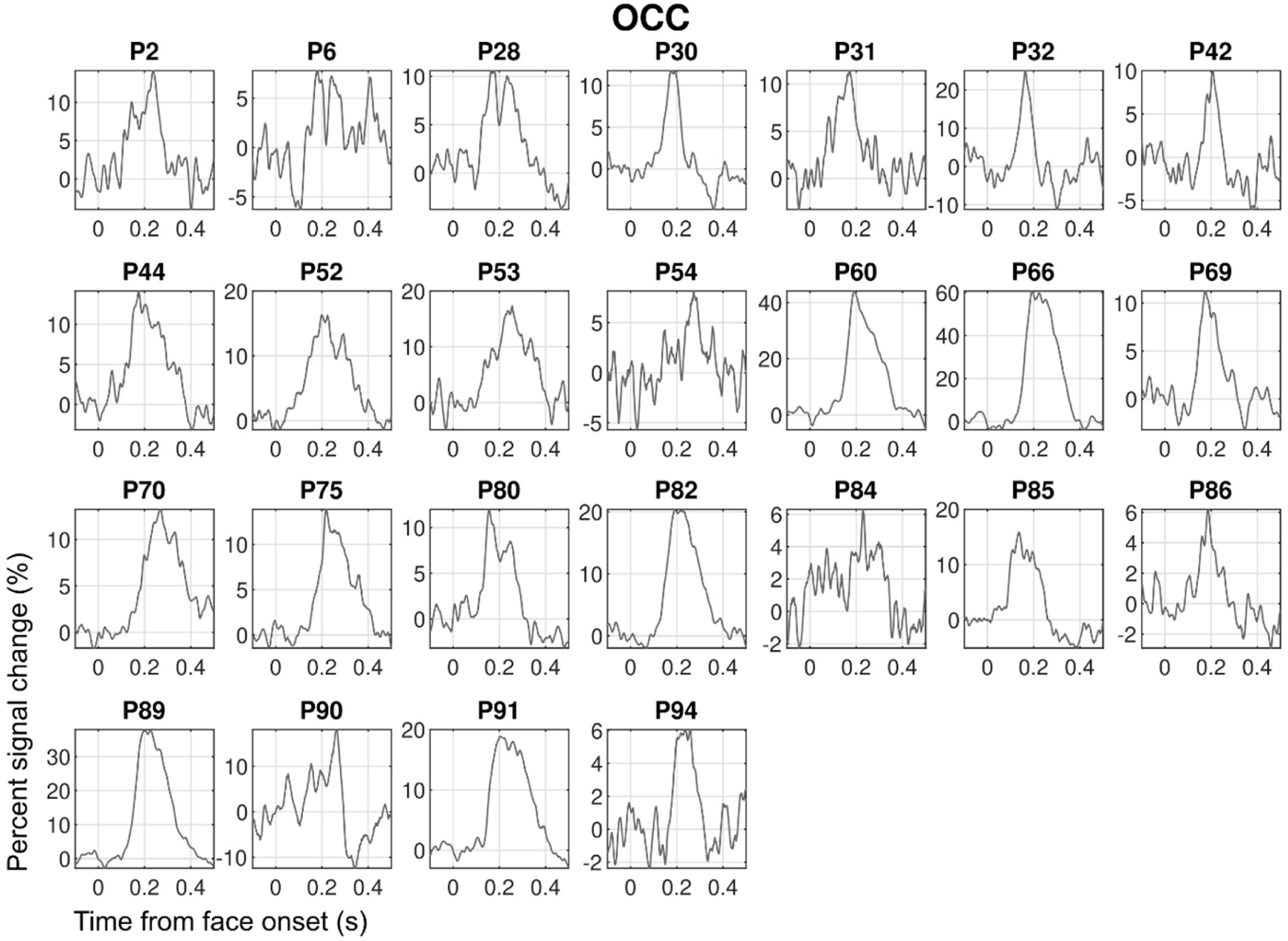
HFB time-domain face-selective response in individual participants for each main VOTC region. Time courses are averaged across contacts within participants. The 6 Hz response to non-face objects has been filtered-out. Part 1: responses in the occipital cortex (OCC).

**Supplementary Figure 11:**
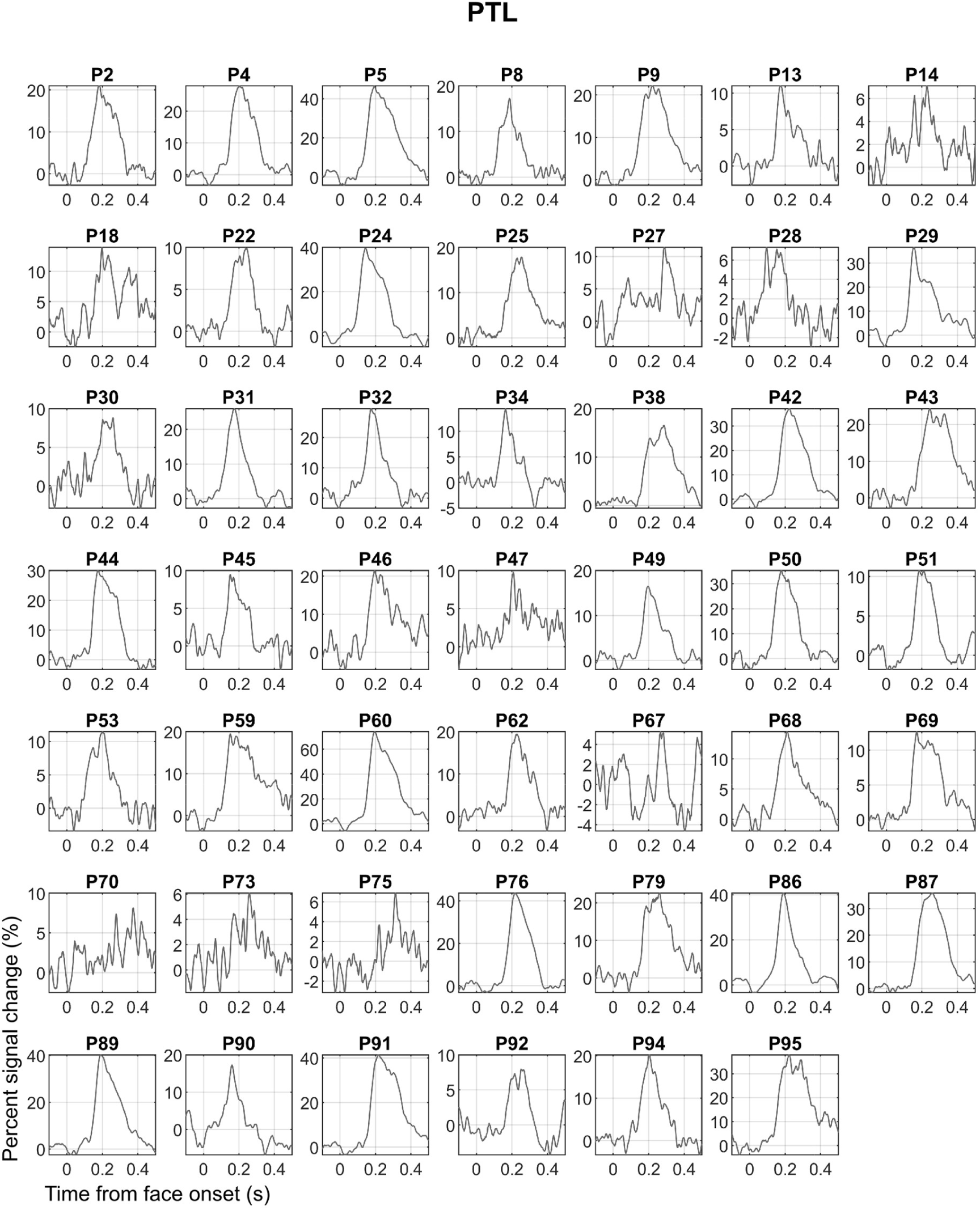
HFB time-domain face-selective response in individual participants for each main VOTC region. Time courses are averaged across contacts within participants. The 6 Hz response to non-face objects has been filtered-out. Part 2: responses in the posterior temporal lobe (PTL).

**Supplementary Figure 11:**
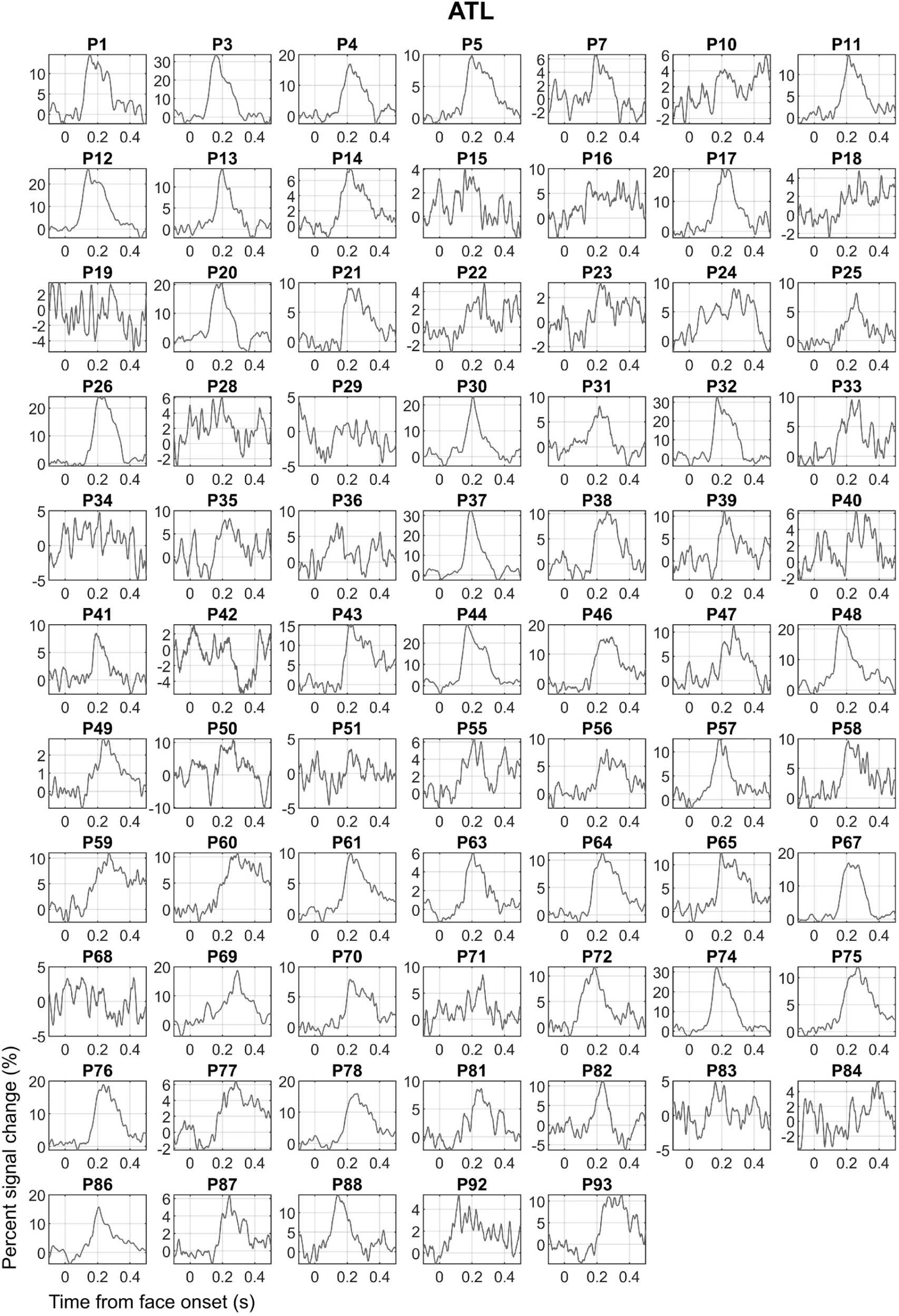
HFB time-domain face-selective response in individual participants for each main VOTC region. Time courses are averaged across contacts within participants. The 6 Hz response to non-face objects has been filtered-out. Part 3: responses in the anterior temporal lobe (ATL).

