## supplemental_material for "Concurrent category-selective neural activity across the ventral occipito-temporal cortex supports a non-hierarchical view of human visual recognition"

**This file includes:**

Figures S1 to S11

Tables S1 - S4

| | Face-selective contacts : response increase | |
|---|---|---|
| | **Left hemisphere** | **Right hemisphere** |
| **VMO** | **6 (4)** | **3 (3)** |
| **IOG** | **19 (10)** | **46 (12)** |
| **medFG** | **7 (6)** | **7 (6)** |
| **latFG** | **58 (21)** | **65 (25)** |
| **MTG/ITG** | **7 (6)** | **2 (2)** |
| **antCOS** | **18 (13)** | **26 (16)** |
| **antFG** | **10 (9)** | **10 (8)** |
| **antOTS** | **63 (29)** | **57 (34)** |
| **antMTG/ITG** | **7 (6)** | **8 (6)** |
| **AMG** | **8 (6)** | **6 (6)** |

**Supplementary Table 1: Number of contacts showing significant face-selective activity increase in each anatomical region.** The corresponding number of participants in which these contacts were found is indicated in parenthesis. Acronyms: VMO: ventro-medial occipital cortex; IOG: inferior occipital gyrus; medFG: medial fusiform gyrus and collateral sulcus; latFG: lateral fusiform gyrus and occipito-temporal sulcus; MTG/ITG: inferior and middle temporal gyri; antCoS: anterior collateral sulcus; antOTS: anterior occipito-temporal sulcus; antFG: anterior fusiform gyrus; antMTG/ITG: anterior middle and inferior temporal gyri.

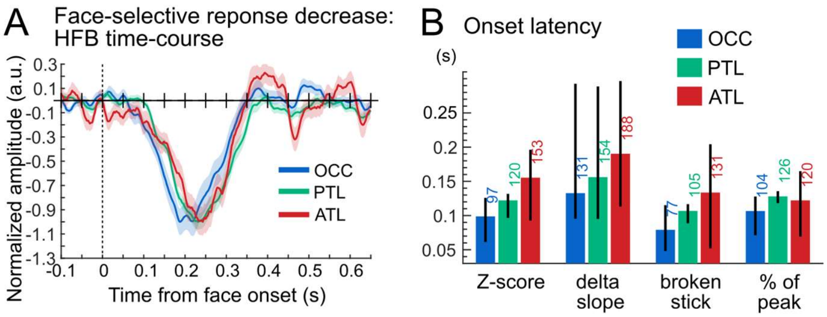


**Supplementary Figure 1: Response timing in contacts showing face-selective response decrease. A.** Time-domain face-selective HFB activity averaged by main VOTC region (OCC, PTL, ATL). HFB time-series were notch-filtered to remove the general visual response at 6 Hz and harmonics, leaving only face-selective signals. The maximum amplitude of each averaged waveform was normalized to -1 for visualization purposes only. Shaded area represents the standard error of the mean between participants. **B**. Onset latency for each VOTC main region and for 4 latency estimation methods, together with 95% confidence interval (percentile bootstrap).

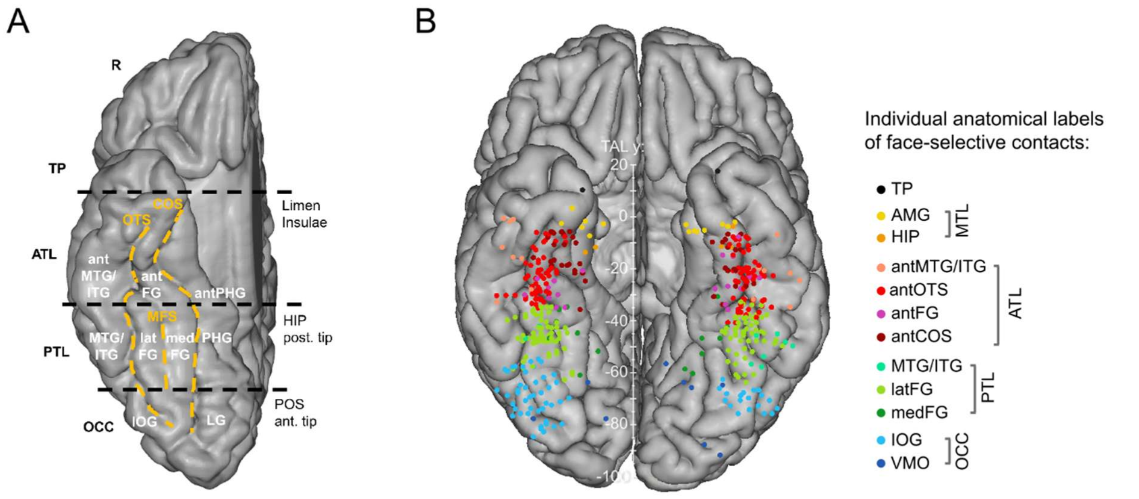


**Supplementary Figure 2: VOTC anatomical parcellation scheme and face-selective recording contacts anatomical labels. A**. Anatomical regions were defined in each individual hemisphere according to major anatomical landmarks. The ventral temporal sulci (COS, OTS, and midfusiform sulcus, i.e., MFS) serve as medial/lateral borders of regions, whereas three coronal reference planes containing anatomical landmarks (OCC-PTL: anterior tip of the parieto-occipital sulcus, i.e., POS, PTL-ATL: posterior tip of the hippocampus, i.e., HIP, ATL-TP: limen insulae) serve as an anterior/posterior boundary for each region. Electrode contacts were considered to be in the ATL if they were located anteriorly to the posterior tip of the hippocampus and posteriorly to the limen insulae. The schematic locations of these anatomical structures are shown on a reconstructed cortical surface of the Colin27 brain. Acronyms: TP: temporal pole; ATL: anterior temporal lobe; PTL: posterior temporal lobe; OCC: occipital lobe; PHG: parahippocampal gyrus; CoS: collateral sulcus; FG: fusiform gyrus; ITG: inferior temporal gyrus; MTG: middle temporal gyrus; OTS: occipito-temporal sulcus; CS: calcarine sulcus; IOG: inferior occipital gyrus; LG: lingual gyrus; ant: anterior; lat: lateral; med: medial. **B.** Map of all face-selective recording contacts and displayed in the Talairach space. Each circle represents a single face-selective contact color-coded according to its anatomical location in the original individual anatomy (see legend on the right).

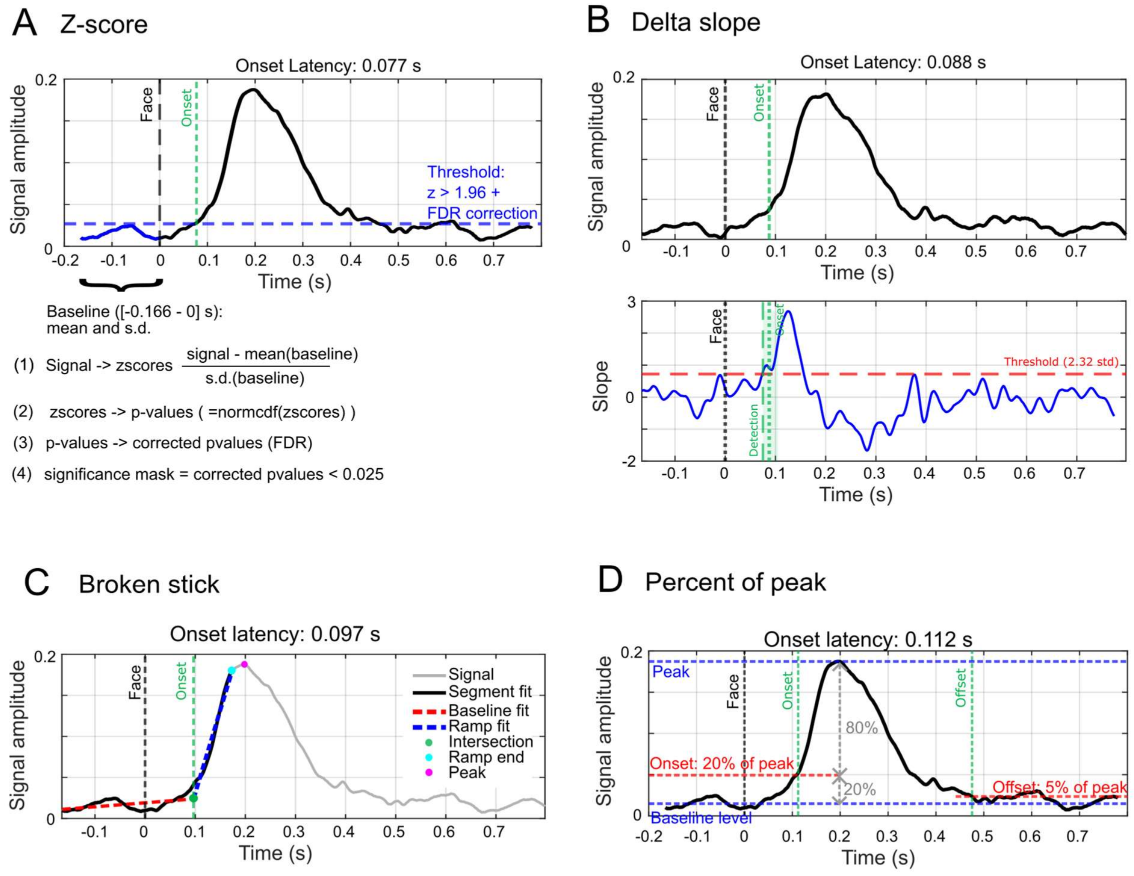


**Supplementary Figure 3: Estimating Onset and offset latencies.** The onset latencies of face-selective activity were characterized using 4 different methods. The offset latency was characterized with one methods (i.e., percent of peak). **A**. In the *z-score* method, the HFB time-series was converted to z-score values by subtracting the mean amplitude in the baseline window of the time-series (i.e. before face onset: [-0.166 to 0s]) and dividing by the standard deviation of the amplitude in the same baseline window. Z-scores were converted to p-values which were FDR-corrected (Benjamini and Hochberg, 1995). Onset latency was the first time-point after 40ms (which was taken as a lower bound for physiologically plausible response latencies after face onset) at which $p < 0.05$ (two-tailed), for at least 30 ms. **B**. In the *delta slope* method, the slopes of the variation in HFB amplitude are computed at each time point within a 25 ms sliding window (see bottom plot). Onset latency is defined as the time (with a lower bound of 40 ms) at which the slope exceeds 2.32 times the mean and standard deviation of the slopes in the baseline window (i.e. an 'acceleration' of the electrophysiological signal manifesting the onset of the neural response), for a duration of at least 30 ms. **C**. In the *broken stick* method, a regression-based method (Mordkoff and Gianaros, 1999), onset latency is defined as the intersection point (lower bound set to 40 ms) of two line segments (with variable slopes) best fitted (least square error fit) to the HFB signal between the start of the baseline window (-166ms) and the end of the ramping-up portion of the signal before the first peak. **D**. In the *percent peak* method, onset latency is defined as the first point, after 40 ms, that rises above 20% of the amplitude difference between the baseline window and the peak (maximum amplitude between 0 and 0.8 s), for at least 30 ms. Offset latency was defined using the 'percent peak' approach, as the point in time, after the onset latency, where the signal reaches below 5% of the amplitude difference between baseline and peak.

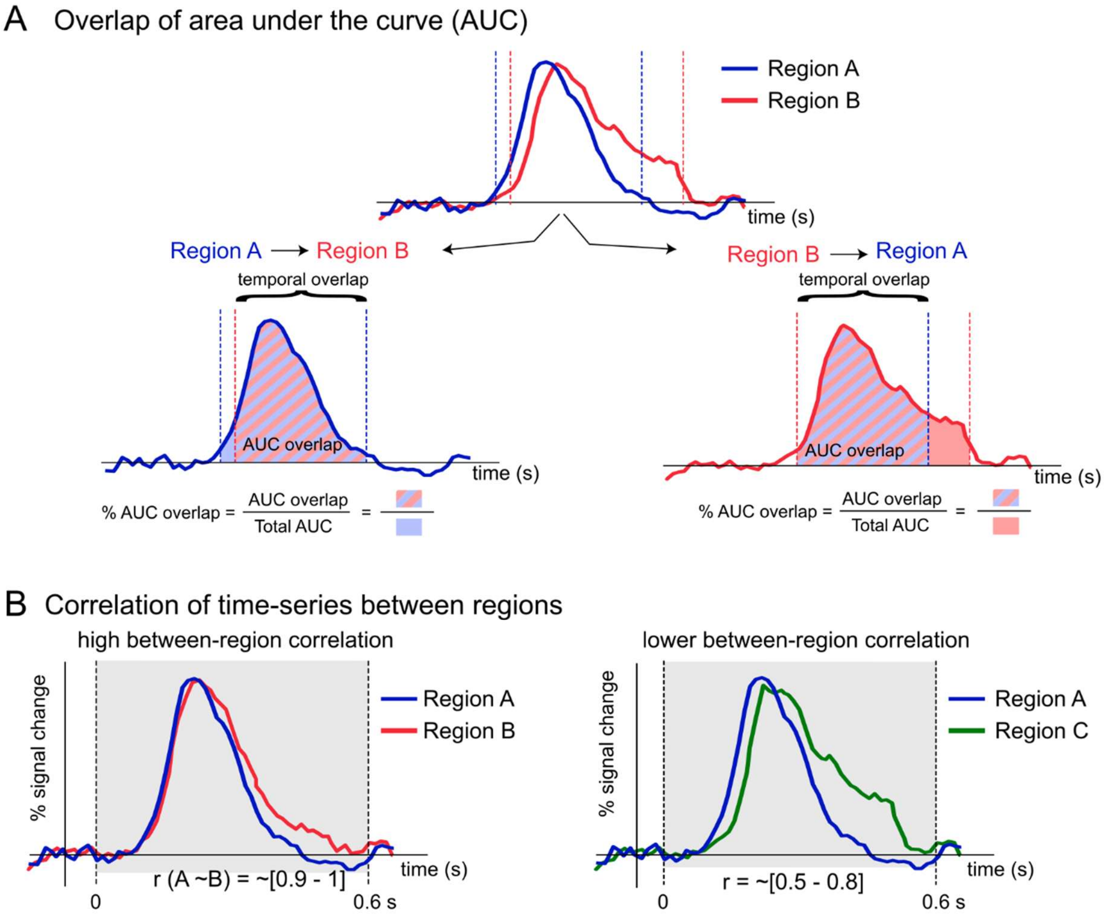


**Supplementary Figure 4: Additional parameters for timing analyses.** The timing of face-selective activity was also characterized by computing response overlap and response correlation across VOTC regions. **A**. A parameter estimated the overlap between time-series for pairs of regions (e.g., regions A and B). The overlap is asymmetrical and is calculated separately for region A and region B as the ratio between the area under the curve (AUC) of the overlap between regions A and region B (i.e., summing the amplitude between the maximum of onset A and B and the minimum of offset A and B) and the total AUC for region A (for overlap of region A to B) or for region B (for overlap of region B to A). **B**. For the last timing parameter, Pearson correlations were computed between the time-series of pairs of regions using data between 0 and 0.6 s relative to face onset (gray shaded area). This provided an estimate of the similarity between the response function of the two regions compared. The shape of the response is more similar between region A and B than between region A and C. Between region correlations were compared against within-region correlations.

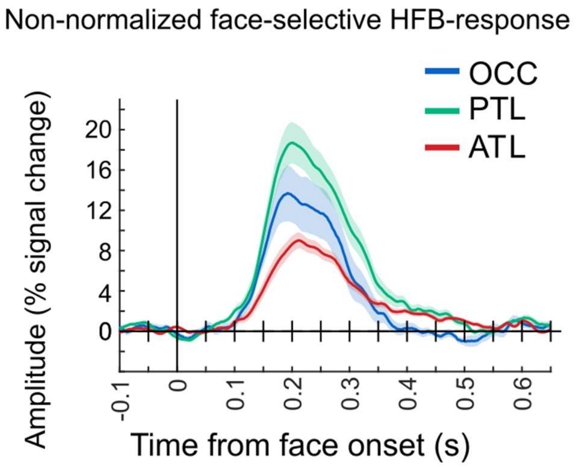


**Supplementary Figure 5: non-normalized time-courses.** Time-domain face-selective HFB activity in contacts showing response increase, averaged by main VOTC region (OCC, PTL, ATL) across hemisphere. HFB time-series were notch-filtered to remove the general visual response at 6 Hz and harmonics, leaving only face-selective signals. Time-series are original non-normalized versions of Figure 4 of the main text. Shaded area represents the standard error of the mean between participants.

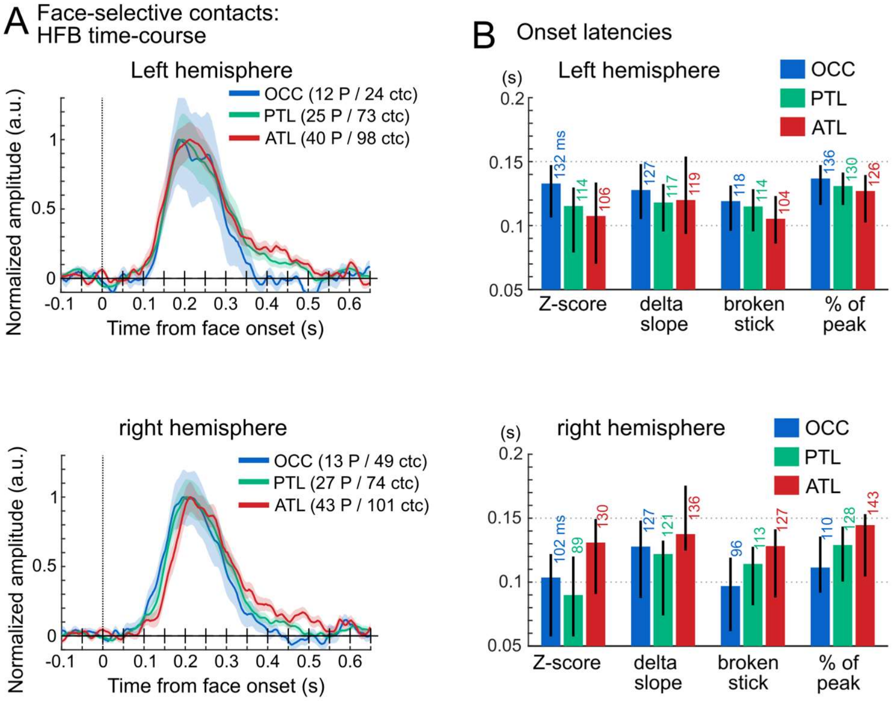


**Supplementary Figure 6: Time-course face-selective contacts split by hemispheres.** Response timing in face-selective contacts grouped by main region separately for left (top) and right (bottom) hemispheres. **A.** Mean time-domain face-selective HFB activity. HFB time-series were notch-filtered to remove the general visual activity at 6 Hz and harmonics, leaving face-selective signals only. The maximum amplitude of each averaged waveform was normalized to 1 for visualization purposes only. Shaded area represents the standard error of the mean between participants. Number of participants and contacts are indicated in the legend. **B**. Onset latency for each VOTC main region and for 4 latency estimation methods, together with 95% confidence interval (percentile bootstrap).

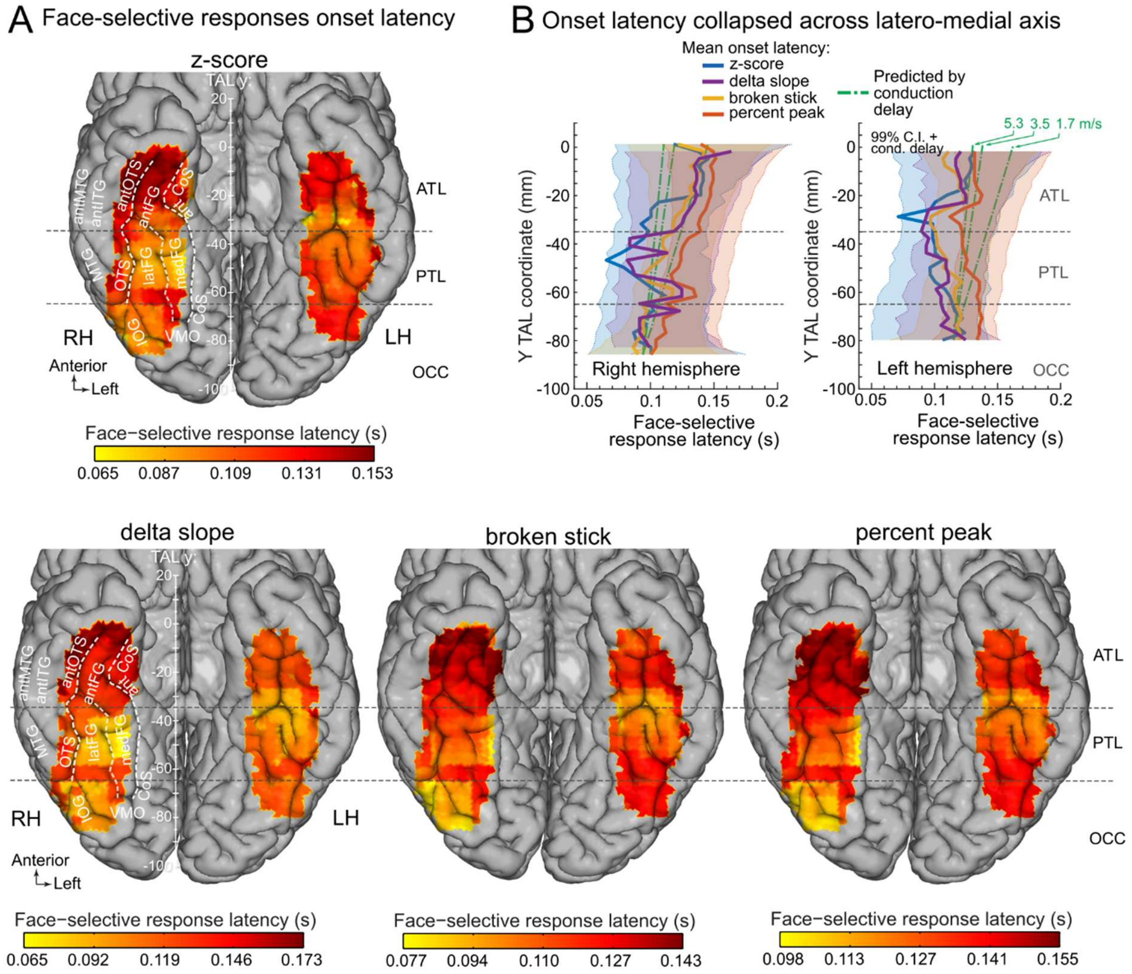


**Supplementary Figure 7: Mapping response onset latency of face-selective contacts split by hemispheres.** A. Face-selective onset latency map across VOTC, displayed for 4 different methods to estimate response onset latency. We collected HFB time-series from contiguous face-selective contacts located within voxels of 20 x 20 x 100 mm (swept across VOTC in steps of 3 x 3 x 100 mm), averaged collected time-series across contacts within-participant and then across participants, and computed onset latency of face-selective response within the current voxels. **B.** Variation of face-selective response latency along the postero-anterior axis for 4 estimation methods and split by hemisphere. For each method, each data point represents the onset latency measured from the time-series averaged over contacts collapsed across the medio-lateral X dimension within 20 mm segments (in the Y dimension). Thick lines are estimated onset latencies and shaded areas show the 99% confidence intervals expected under the null hypothesis that the postero-anterior location has no influence on the onset latency, accounting for the expected conduction delays between posterior and anterior VOTC. Green lines show the expected increase in response onset latency based on simple axonal conduction delays(Lemarechal et al., 2022; van Blooijs et al., 2023) due to increasing distance from the occipital region, with reference to the latency averaged over the OCC region for the 'z-score' method. Lines show mean expected conduction velocity for direct cortico-cortical connections (~3.5 m/s) and +/- 1 std (1.7 and 5.3 m/s).

| Latency method | Hemi | Region comparison | Median diff. (ms) | Cohen's d | pval. FDR | Nb Partcpts region 1 | Nb Partcpts region 2 | Equivalence Prop in ROPE (%) | Bayes factor Cauchy prior (>3) |
|---|---|---|---|---|---|---|---|---|---|
| z-score | Left | OCC vs PTL | 17.6 | 0.32 | 0.9228 | 12 | 25 | 29 | 0.7 |
| delta slope | Left | OCC vs PTL | 7.8 | 0.19 | 0.9228 | 12 | 25 | 46 | 1.3 |
| Broken stick | Left | OCC vs PTL | 3.9 | 0.11 | 1 | 12 | 25 | 60 | 2.2 |
| % of peak | Left | OCC vs PTL | 5.9 | 0.15 | 0.98475789 | 12 | 25 | 54 | 1.8 |
| z-score | Left | PTL vs ATL | 2 | 0.04 | 1 | 25 | 40 | 58 | 2.4 |
| delta slope | Left | PTL vs ATL | 0 | 0 | 0.9228 | 25 | 40 | 82 | 11.6 |
| Broken stick | Left | PTL vs ATL | 9.1 | 0.15 | 0.93066667 | 25 | 40 | 57 | 2.4 |
| % of peak | Left | PTL vs ATL | 5.9 | 0.12 | 1 | 25 | 40 | 68 | 3.6 |
| z-score | Left | OCC vs ATL | 21.5 | 0.47 | 1 | 12 | 40 | 23 | 0.4 |
| delta slope | Left | OCC vs ATL | 5.9 | 0.03 | 0.9228 | 12 | 40 | 71 | 5.9 |
| Broken stick | Left | OCC vs ATL | 13.1 | 0.21 | 0.93066667 | 12 | 40 | 47 | 1.4 |
| % of peak | Left | OCC vs ATL | 9.8 | 0.2 | 0.9228 | 12 | 40 | 44 | 1.1 |
| z-score | Right | OCC vs PTL | 11.7 | 0.14 | 1 | 13 | 27 | 56 | 2.6 |
| delta slope | Right | OCC vs PTL | 3.9 | 0.02 | 0.9228 | 13 | 27 | 70 | 6 |
| Broken stick | Right | OCC vs PTL | -15.9 | 0.31 | 0.9228 | 13 | 27 | 61 | 2.7 |
| % of peak | Right | OCC vs PTL | -15.6 | 0.3 | 0.9228 | 13 | 27 | 71 | 4.2 |
| z-score | Right | PTL vs ATL | -39.1 | 0.56 | 0.9228 | 27 | 43 | 36 | 1.1 |
| delta slope | Right | PTL vs ATL | -17.6 | 0.11 | 0.9228 | 27 | 43 | 75 | 7.3 |
| Broken stick | Right | PTL vs ATL | -14.2 | 0.24 | 0.9228 | 27 | 43 | 71 | 4.4 |
| % of peak | Right | PTL vs ATL | -13.7 | 0.26 | 0.9228 | 27 | 43 | 72 | 4.4 |
| z-score | Right | OCC vs ATL | -27.3 | 0.42 | 0.9228 | 13 | 43 | 60 | 2.3 |
| delta slope | Right | OCC vs ATL | -15.6 | 0.6 | 0.9228 | 13 | 43 | 41 | 0.7 |
| Broken stick | Right | OCC vs ATL | -30.8 | 0.55 | 0.9228 | 13 | 43 | 58 | 2 |
| % of peak | Right | OCC vs ATL | -31.3 | 0.68 | 0.9228 | 13 | 43 | 48 | 1.3 |

**Supplementary Table 2** : Statistical comparisons of face-selective onset latencies across regions (OCC vs PTL, PTL vs ATL, OCC vs ATL) in each hemisphere using permutation tests et equivalence testing (proportion of difference in ROPE and Bayes factor) for 4 different onset estimation methods. Table contains the median latency difference between regions (Median diff.), effect size (Cohen’s d) of the difference, p-value of the permutation test (pval. FDR), number of participants included in each region.

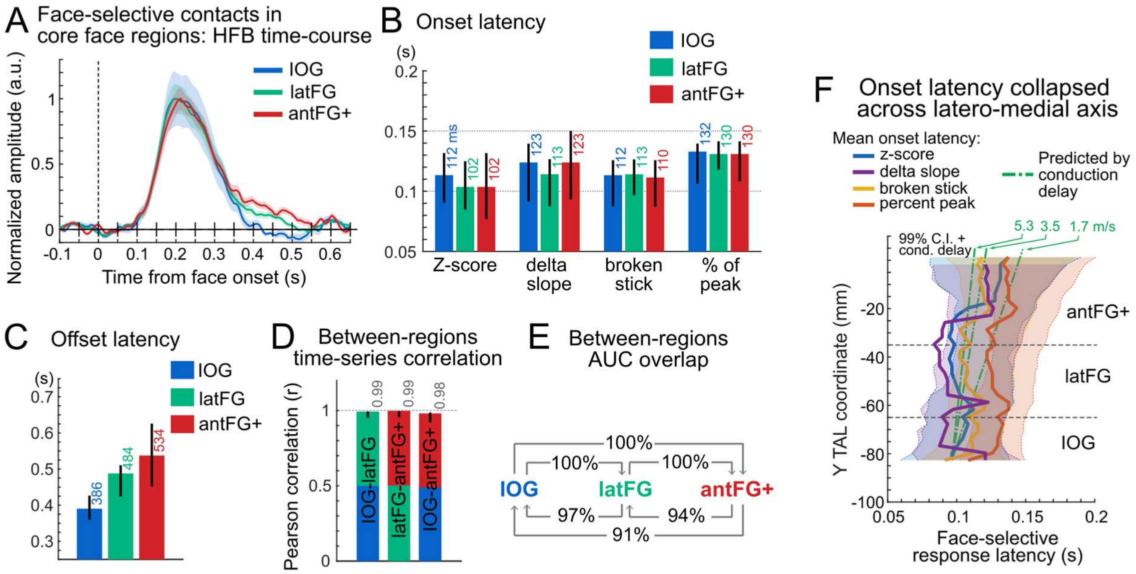


**Supplementary Figure 8: Response timing in face-selective contacts in core VOTC face-selective regions.** Response timing in face-selective contacts showing response increase in core VOTC face-selective regions: Inferior Occipital Gyrus (IOG, (N=65), lateral Fusiform Gyrus and OTS (latFG, N=123) and antFG+ (anterior fusiform Gyrus, Anterior Occipitotemporal Sulcus, Anterior Collateral Sulcus, N=184). **A.** Mean time-domain face-selective HFB activity in each core face-selective VOTC region collapsed across hemisphere. HFB time-series were filtered to remove the general visual response at 6 Hz and harmonics so only face-selective signal remains. The maximum amplitude of each averaged waveform was normalized to 1 for visualization purposes only. Shaded area represents the standard error of the mean between participants. **B**. Onset latency for each VOTC main region and for 4 latency estimation methods, together with 95% confidence interval (percentile bootstrap). **C**. Offset latency in each region. **D**. Correlation of time-series between regions and 95% confidence intervals. **E**. Between-region area under the curve (AUC) overlap. The arrow shows the directionality of the computed overlap. For instance, the arrow from ATL to OCC indicates the percentage of the total AUC of ATL (measured between onset and offset latencies) occupied by the AUC of its temporal overlap with OCC (determined between max(onset(OCC,ATL)) and min(offset(OCC,ATL)) ). **F.** Variation of face-selective response latency along the postero-anterior axis for 4 estimation methods. For each method, each data point represents the onset latency measured from the time-series averaged over contacts collapsed across the medio-lateral X dimension within 20 mm segments (in the Y dimension). Thick lines are estimated onset latencies and shaded areas show the 99% confidence intervals expected under the null hypothesis that the postero-anterior location has no influence on the onset latency, accounting for the expected conduction delays between posterior and anterior VOTC. Green lines show the expected increase in response onset latency based on simple axonal conduction delays (Lemarechal et al., 2022; van Blooijs et al., 2023)due to increasing distance from the occipital region, with reference to the latency averaged over the OCC region for the ‘z-score’ method. Lines show mean expected conduction velocity for direct cortico-cortical connections (~3.5 m/s) and +/- 1 std (1.7 and 5.3 m/s).

| Latency method | Region comparison | Median diff. (ms) | Cohen's d | pval. FDR | Nb Partcpts region 1 | Nb Partcpts region 2 | Equivalence Prop in ROPE (%) | Bayes factor Cauchy prior (>3) |
|---|---|---|---|---|---|---|---|---|
| z-score | IOG vs latFG | 7.8 | 0.16 | 1 | 22 | 43 | 50 | 1.7 |
| delta slope | IOG vs latFG | 9.8 | 0.07 | 1 | 22 | 43 | 82 | 10.9 |
| Broken stick | IOG vs latFG | -1.4 | 0.03 | 1 | 22 | 43 | 79 | 6.1 |
| % of peak | IOG vs latFG | 0 | 0 | 1 | 22 | 43 | 85 | 10.1 |
| z-score | latFG vs antFG+ | -2 | 0.04 | 1 | 43 | 72 | 63 | 2.7 |
| delta slope | latFG vs antFG+ | -9.8 | 0.09 | 1 | 43 | 72 | 90 | 19.8 |
| Broken stick | latFG vs antFG+ | 4.4 | 0.05 | 1 | 43 | 72 | 79 | 7.5 |
| % of peak | latFG vs antFG+ | 0 | 0 | 1 | 43 | 72 | 87 | 12.8 |
| z-score | IOG vs antFG+ | 5.9 | 0.13 | 1 | 22 | 72 | 53 | 1.5 |
| delta slope | IOG vs antFG+ | 0 | 0 | 1 | 22 | 72 | 73 | 3.6 |
| Broken stick | IOG vs antFG+ | 2.8 | 0.03 | 1 | 22 | 72 | 84 | 9 |
| % of peak | IOG vs antFG+ | 0 | 0 | 1 | 22 | 72 | 89 | 12.8 |

**Supplementary Table 3** : Statistical comparisons of face-selective onset latencies measured in **HFB signal** across **core face-selective regions** (IOG, latFG, antFG) using permutation tests et equivalence testing (proportion of difference in ROPE and Bayes factor) for 4 different onset estimation methods (same as in main). Table contains the median latency difference between regions (Median diff.), effect size (Cohen's d) of the difference, p-value of the permutation test (pval. FDR), number of participants included in each region.

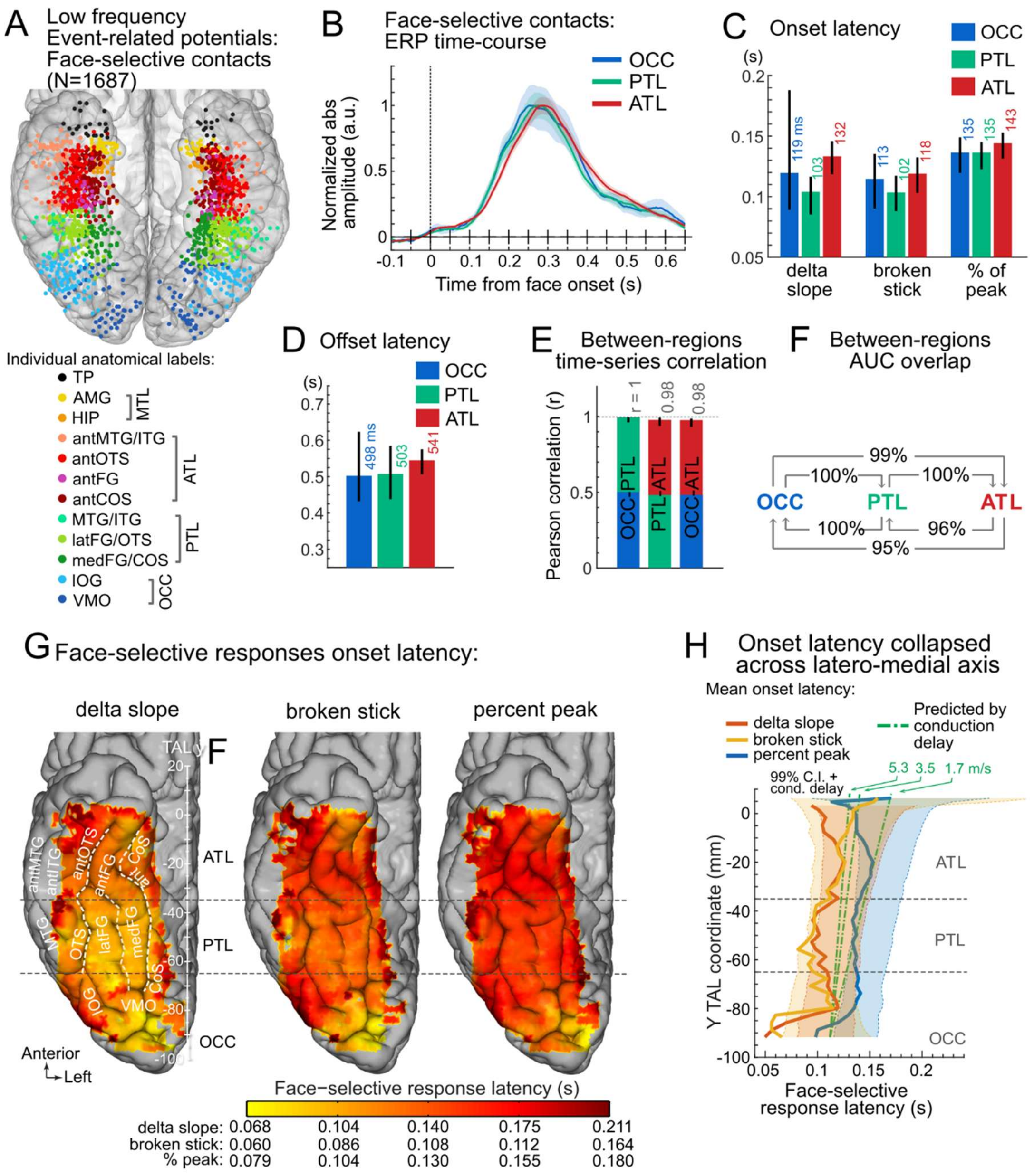


**Supplementary Figure 9: Face-selective activity, low frequency event-related potentials (ERPs)**. **A.** Map of all VOTC contacts showing a significant (Z>3.1) face-selective response in low-frequency ERPs, displayed in the Talairach space in a transparent reconstructed cortical surface of the Colin27 brain (ventral view). Each circle represents a single recording contact. Contacts are color-coded according to their anatomical location in the original individual anatomy. Low frequency signal is the electrophysiological response measured at each recording contact after bipolar re-referencing (Jacques et al., 2022), dominated by low frequency event-related voltage variations. Only contacts with split-half correlation in the time-domain (i.e. reliability) > 0.66 (i.e. 75% of contacts) were kept to unsure reliable latency estimations. **B.** Grand average face-selective ERP in each face-selective VOTC region collapsed across hemispheres. ERP time-series were filtered to remove the general visual response at 6 Hz and harmonics, leaving face-selective signals only. Averaged ERP were baseline-corrected relative to a [-0.166 0]s window before face onset and the absolute value was computed to allow averaging across contacts/participants despite variability in ERP polarity and morphology. ERP at individual recording contacts were averaged by participants and then across participants in each region. The maximum amplitude of each averaged waveform was normalized to 1 for visualization purposes only. Shaded area represents the standard error of the mean between participants. **C**. Onset latency for each VOTC main region and for 3 latency estimation methods (same as in main text except for 'z-score' method, which is unusable with low frequencies), together with 95% confidence interval (percentile bootstrap). **D**. Offset latency in each region. **E**. Correlation of time-series between regions and 95% confidence intervals. **F**. Between-region area under the curve (AUC) overlap. The arrow shows the directionality of the computed overlap. For instance, the arrow from ATL to OCC indicates the percentage of the total AUC of ATL (measured between onset and offset latencies) occupied by the AUC of its temporal overlap with OCC (determined between max(onset(OCC,ATL)) and min(offset(OCC,ATL))). **G.** Face-selective onset latency map across VOTC, collapsed across hemispheres and displayed for 3 different methods to estimate response onset latency. **H.** Variation of face-selective response latency along the postero-anterior axis for 3 estimation methods. For each method, each data point represents the onset latency measured from the time-series averaged over contacts collapsed across the medio-lateral X dimension within 20 mm segments (in the Y dimension). Thick lines are estimated onset latencies and shaded areas show the 99% confidence intervals expected under the null hypothesis that the postero-anterior location has no influence on the onset latency, accounting for the expected conduction delays between posterior and anterior VOTC. Green lines show the expected increase in response onset latency based on axonal conduction delays(Lemarechal et al., 2022; van Blooijs et al., 2023) due to increasing distance from the occipital region, with reference to the latency averaged over the OCC region for the 'z-score' method. Lines show mean expected conduction velocity for direct cortico-cortical connections (~3.5 m/s) and +/- 1 std (1.7 and 5.3 m/s). Note that response latency from few recording contacts located posterior to y = -85 (around and posterior/medial to the posterior transverse Collateral Sulcus), likely in early visual cortex (V1,V2v,V3v,hV4, (Brewer et al., 2005; Winawer and Witthoft, 2015)) and potentially driven by low-level cues (Or et al., 2019), is up to 40 ms earlier than responses measured elsewhere in the VOTC, where little difference is observed.

| Latency method | Region comparison | Median diff. (ms) | Cohen's d | pval. FDR | Nb Partcpts region 1 | Nb Partcpts region 2 | Equivalence Prop in ROPE (%) | Bayes factor Cauchy prior (>3) |
|---|---|---|---|---|---|---|---|---|
| Broken stick | OCC_vs_PTL | 11.1 | 0.22 | 0.343125 | 41 | 88 | 47 | 1.5 |
| % of peak | OCC_vs_PTL | 0 | 0 | 0.876 | 41 | 88 | 89 | 13.2 |
| delta slope | OCC_vs_PTL | 15.6 | 0.23 | 0.1905 | 41 | 88 | 43 | 1.4 |
| Broken stick | PTL_vs_ATL | -15.7 | 0.31 | 0.004498 | 88 | 113 | 84 | 8.9 |
| % of peak | PTL_vs_ATL | -7.8 | 0.1 | 0.06075 | 88 | 113 | 100 | 666.9 |
| delta slope | PTL_vs_ATL | -29.3 | 0.33 | 0.004498 | 88 | 113 | 63 | 3.5 |
| Broken stick | OCC_vs_ATL | -5.2 | 0.09 | 0.1905 | 41 | 113 | 88 | 10.4 |
| % of peak | OCC_vs_ATL | -7.8 | 0.09 | 0.06075 | 41 | 113 | 100 | 3492.4 |
| delta slope | OCC_vs_ATL | -11.7 | 0.14 | 0.255857 | 41 | 113 | 83 | 8.9 |

**Supplementary Table 4** : Statistical comparisons of face-selective onset latencies measured in **low frequency event-related potentials** across regions (OCC vs PTL, PTL vs ATL, OCC vs ATL) using permutation tests et equivalence testing (proportion of difference in ROPE and Bayes factor) for 3 different onset estimation methods (same as in main text except for ‘z-score’ method, which is unusable with low frequencies). Table contains the median latency difference between regions (Median diff.), effect size (Cohen’s d) of the difference, p-value of the permutation test (pval. FDR), number of participants included in each region. Despite significantly later ATL than PTL onset latencies (with small effect sizes), the difference is still within bounds of equivalence when taking into account estimated axonal conduction delays between the two regions.

**Functional connectivity using mutual information**

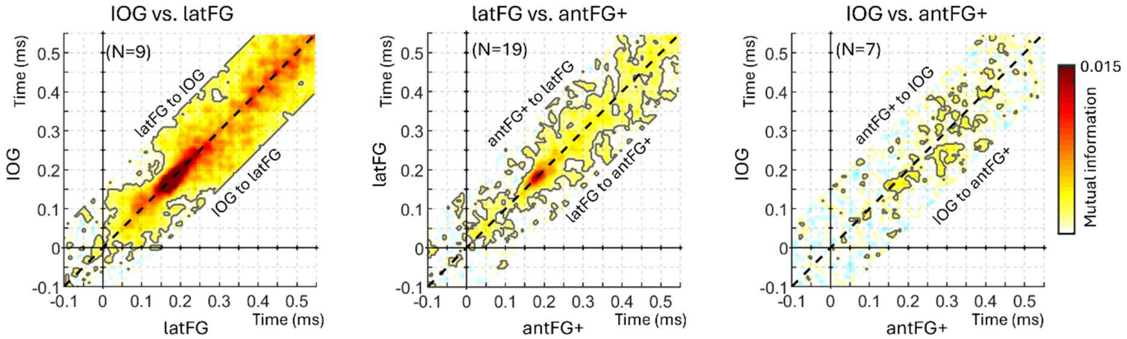


**Supplementary Figure 10: Concurrent functional connectivity between face-selective regions computed using mutual information.** Each plot shows the group-averaged mutual information (MI) between single-trials face-selective amplitude measured at two distinct face-selective region (left: IOG and latFG; middle: latFG and antFG+; right: IOG and antFG+) computed at each time point, representing functional connectivity between pairs of regions. MI was computed using the method and Matlab code described in (Ince et al., 2017). Statistics were performed using permutation tests as for Pearson correlations presented in the main manuscript. Black contour lines indicate significant MI ($p$ < 0.01, fdr-corrected). MI was computed across -150 to 150 ms lags between regions to infer direction of connectivity. The black dashed diagonal line represents a 0 ms time-lag between regions. MI centered above the diagonal would indicate that face-selective activity in the more anterior region (e.g. latFG in left panel) is coupled with but precedes activity in the posterior region (e.g., IOG in left panel), suggesting an information flow from anterior to posterior, and the reverse for MI centered below the diagonal.

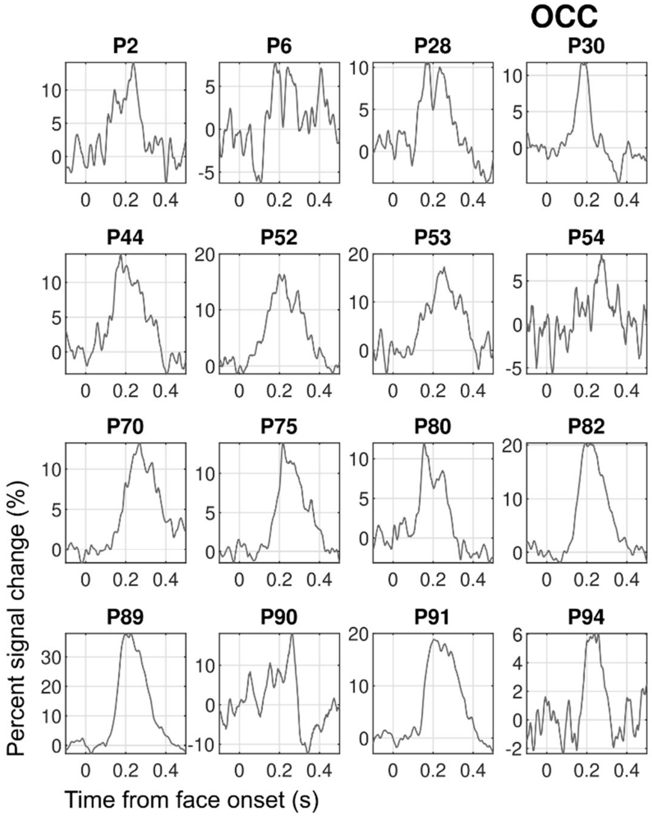


**Supplementary Figure 11: HFB time-domain face-selective response in individual participants for each main VOTC region.** Time courses are averaged across contacts within participants. The 6 Hz response to non-face objects has been filtered-out. Part 1: responses in the occipital cortex (OCC).

### PTL

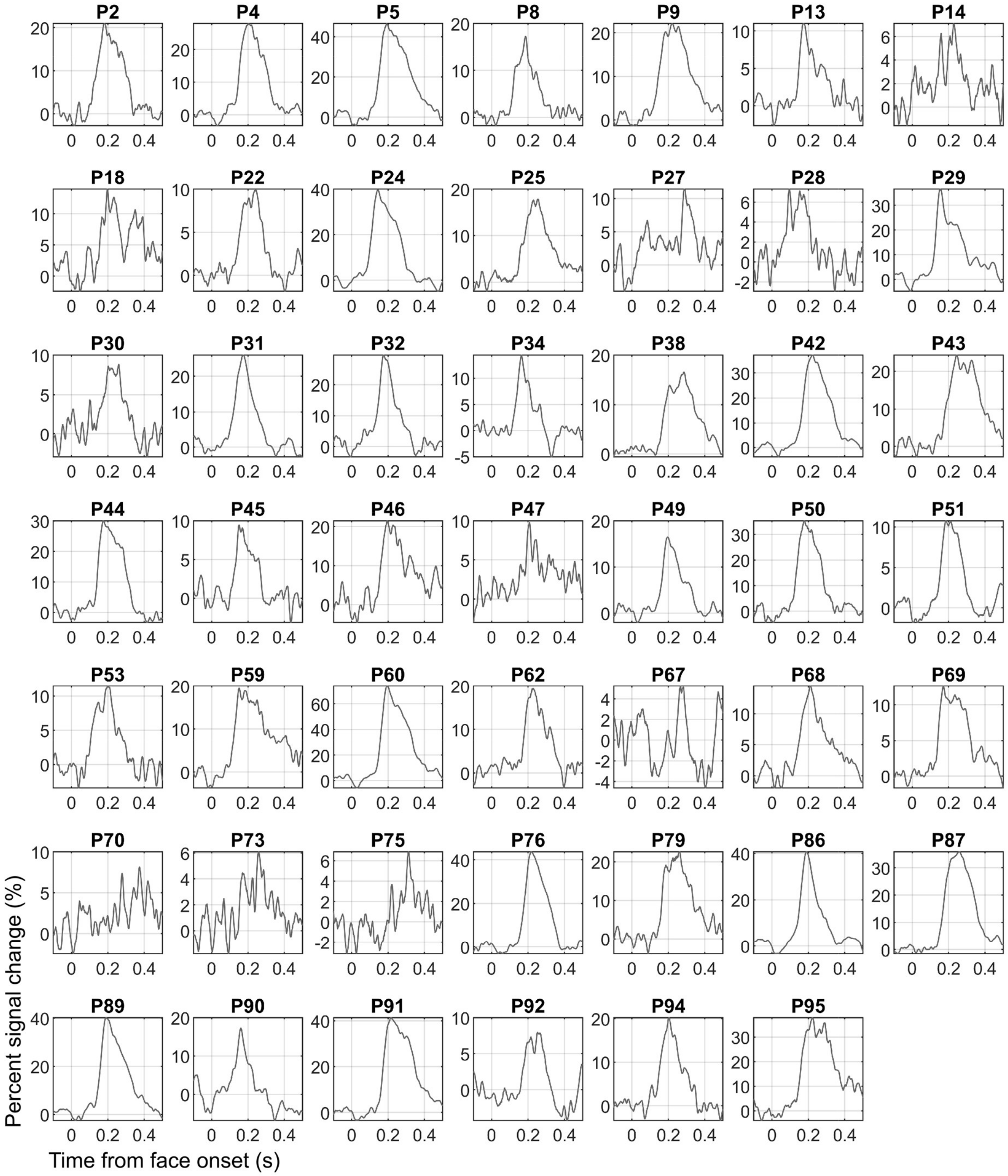


**Supplementary Figure 11: HFB time-domain face-selective response in individual participants for each main VOTC region.** Time courses are averaged across contacts within participants. The 6 Hz response to non-face objects has been filtered-out. Part 2: responses in the posterior temporal lobe (PTL).

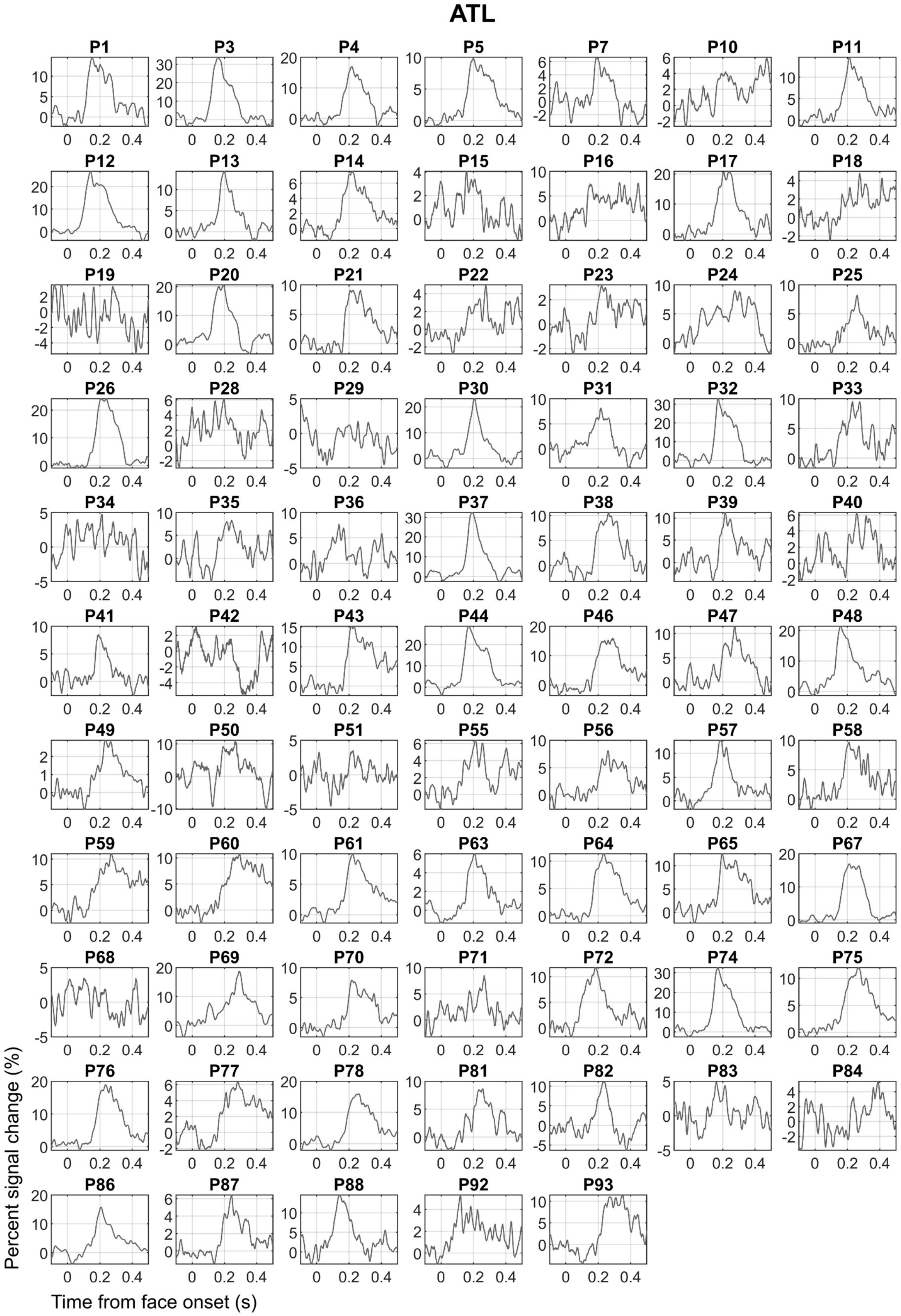


**Supplementary Figure 11: HFB time-domain face-selective response in individual participants for each main VOTC region.** Time courses are averaged across contacts within participants. The 6 Hz response to non-face objects has been filtered-out. Part 3: responses in the anterior temporal lobe (ATL).

**Supplemental Material references**:


Brewer AA, Liu J, Wade AR, Wandell BA. 2005. Visual field maps and stimulus selectivity in human ventral occipital cortex. Nature Neuroscience **8**:1102–1109. DOI: https://doi.org/10.1038/nn1507, PMID: 16025108

Ince RAA, Giordano BL, Kayser C, Rousselet GA, Gross J, Schyns PG. 2017. A statistical framework for neuroimaging data analysis based on mutual information estimated via a gaussian copula. Human brain mapping **38**:1541–1573. DOI: https://doi.org/10.1002/HBM.23471, PMID: 27860095

Jacques C, Jonas J, Colnat-Coulbois S, Maillard L, Rossion B. 2022. Low and high frequency intracranial neural signals match in the human associative cortex. eLife **11**:e76544. DOI: https://doi.org/10.7554/eLife.76544, PMID: 36074548

Lemarechal JD, Jedynak M, Trebaul L, Boyer A, Tadel F, Bhattacharjee M, Deman P, Tuyisenge V, Ayoubian L, Hugues E, Chanteloup-Forêt B, Saubat C, Zouglech R, Reyes Mejia GC, Tourbier S, Hagmann P, Adam C, Barba C, Bartolomei F, Blauwblomme T, Curot J, Dubeau F, Francione S, Garces M, Hirsch E, Landre E, Liu S, Maillard L, Metsähonkala EL, Mindruta I, Nica A, Pail M, Petrescu AM, Rheims S, Rocamora R, Schulze-Bonhage A, Szurhaj W, Taussig D, Valentin A, Wang H, Kahane P, George N, David O, Adam C, Navarro V, Biraben A, Nica A, Menard D, Brazdil M, Kuba R, Kočvarová J, Pail M, Dolealová I, Dubeau F, Gotman J, Ryvlin P, Isnard J, Catenoix H, Montavont A, Rheims S, Trebuchon A, Mcgonigal A, Zhou W, Wang H, Liu S, Wei Z, Dan Z, Qiang G, Xiangshu H, Hua L, Gang H, Wensheng W, Xi M, Yigang F, Nabbout R, Bourgeois M, Kaminska A, Blauwblomme T, Garces M, Valentin A, Singh R, Metsähonkala L, Gaily E, Lauronen L, Peltola M, Chassoux F, Landre E, Derambure P, Szurhaj W, Chochois M, Hirsch E, Paola Valenti M, Scholly J, Valton L, Denuelle M, Curot J, Rocamora R, Principe A, Ley M, Mindruta I, Barborica A, Francione S, Mai R, Nobili L, Sartori I, Tassi L, Maillard L, Vignal JP, Jonas J, Tyvaert L, Chipaux M, Taussig D, Kahane P, Minotti L, Job AS, Michel V, De Montaudoin M, Aupy J, Bouilleret V, Maria Petrescu A, Masnou P, Dussaule C, Quirins M, Taussig D, Guerrini R, Lenge M, Nacci E. 2022. A brain atlas of axonal and synaptic delays based on modelling of cortico-cortical evoked potentials. Brain **145**:1653–1667. DOI: https://doi.org/10.1093/brain/awab362

Mordkoff JT, Gianaros PJ. 1999. Detecting the onset of the lateralized readiness potential: A comparison of available methods and procedures.

Or CCF, Retter TL, Rossion B. 2019. The contribution of color information to rapid face categorization in natural scenes. Journal of vision **19**:1–20. DOI: https://doi.org/10.1167/19.5.20, PMID: 31112241

van Blooijs D, van den Boom MA, van der Aar JF, Huiskamp GM, Castegnaro G, Demuru M, Zweiphenning WJEM, van Eijsden P, Miller KJ, Leijten FSS, Hermes D. 2023. Developmental trajectory of transmission speed in the human brain. Nature Neuroscience **26**:537–541. DOI: https://doi.org/10.1038/s41593-023-01272-0, PMID: 36894655

Winawer J, Witthoft N. 2015. Human V4 and ventral occipital retinotopic maps. Visual neuroscience **32**:E020. DOI: https://doi.org/10.1017/S0952523815000176, PMID: 26241699
